# Rational Design of a Photoactivated Fluorescent Probe for Thiol-ene Mediated Protein Labelling

**DOI:** 10.64898/2026.10.13.758905

**Authors:** Ben H. McLoughlin, Conor Williams, Sean M. McKenna, Kosta Besermenji, Ivana Drienovská, Marinda Westerveld, Inés Rabadán González, Susannah H. Calvert, Joanna F. McGouran, Eoin M. Scanlan, Mark D. Nolan, Rita Petracca

**Affiliations:** Trinity Biomedical Sciences Institute, Trinity College Dublin, 152-160 Pearse Street, Dublin 2, Ireland; Department of Pharmaceutical Sciences, Faculty of Science, Utrecht University, David De Wied Building, Universiteitsweg 99, 3584 CG Utrecht, The Netherlands; Department of Chemistry and Pharmaceutical Sciences, Vrije Universiteit Amsterdam, De Boelelaan 1105, 1081HV Amsterdam, The Netherlands; Department of Chemistry, Princeton University, Princeton, NJ, USA

**Keywords:** “Fluorescent Probe”, “Thiol–ene”, “Labelling”, “Radical”, “Cell Lysate”

## Abstract

Fluorescent labelling of biomolecules enables the visualization and tracking of molecular processes in cells, providing insights into cellular function and disease mechanisms. Despite recent advances in bioorthogonal labelling, efficient and site-selective strategies for targeting alkene functionalities remain limited. Here, we report AbzSH, a compact aminobenzamide-based fluorescent thiol probe for selective alkene labelling through thiol–ene ligation. Photoinduced ligation proceeds under mild, radical-mediated conditions in aqueous media and exhibits broad substrate compatibility, including carbohydrate, amino acid, and steroid scaffolds. The method is readily extended to protein labelling, enabling selective modification of both chemically functionalized proteins and engineered proteins containing genetically encoded alkene motifs. Efficient and chemoselective labelling was further demonstrated in complex cell lysates, highlighting the robustness of the approach in biologically relevant environments. Collectively, these results establish AbzSH as a versatile fluorescent probe for alkene-directed biomolecular labelling and provide a broadly applicable platform for chemical biology, molecular imaging, proteomics, and drug discovery.

**TOC Graphic:** Thiol-ene mediated ‘click’ labelling of an alkene-bearing protein in complex cell lysate using a fluorescent thiol probe.

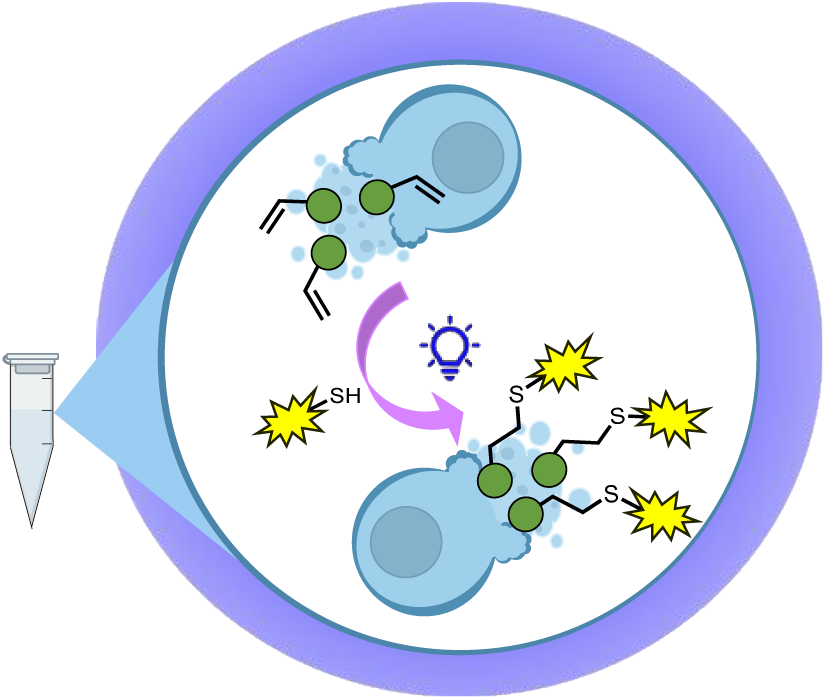

---

Dysfunction in biomolecular structure or localization is implicated in numerous disease states.^1–4^ Accordingly, the ability to selectively probe biomolecules within complex biological environments is central to understanding disease pathophysiology and advancing therapeutic development. Fluorescent labelling enables visualization and tracking of dynamic biomolecular processes, including uptake, localization, metabolism and aggregation in real time.^5–8^ Established fluorescent labelling strategies often rely on highly selective bioorthogonal transformations, including azide-alkyne cycloaddition, Staudinger ligation and inverse electron demand Diels-Alder.^9–11^ While exceptionally robust, these approaches often require prior installation of bulky non-native chemical handles, which can necessitate additional synthetic or genetic manipulation and, depending on the system, may perturb molecular behaviour or conformation.^12,13^ Furthermore, certain labelling strategies, such as copper-catalysed azide-alkyne cycloaddition, present additional challenges due to the potential toxicity of the catalyst.^14^ The development of selective fluorescent labelling methods that exploit intrinsic or minimally perturbing functionalities therefore remains of considerable interest.

A particularly attractive reaction for alkene-directed labelling is the radical-mediated, thiol–ene ‘click’ (TEC) reaction.^15^ This process enables the efficient addition of thiol moieties across carbon–carbon double bonds, to furnish stable thioether linkages under mild conditions. The TEC reaction is radical-mediated, can be initiated *via* mild photochemical or thermal methods and is compatible with a broad range of unsaturated residues, including both terminal and internal alkenes.^15,16^ The rapid kinetics, broad functional group tolerance, and compatibility with aqueous conditions renders TEC particularly well suited to chemical biology applications.^15,17–19^ In addition, photochemical initiation provides precise spatiotemporal control, allowing ligation to be activated only upon irradiation. Developments in recent years have demonstrated TEC under increasingly mild initiation conditions, including the use of blue or green LEDs, further increasing biocompatibility and allowing for labelling applications involving living cells.^20,21^ Previous studies of TEC-based biological labelling have largely focused on the modification of biological thiols using alkene-based probes, for example, cysteine-directed ligation and protein crosslinking.^17,18,22,23^ However, achieving selective modification of individual cysteine residues remains challenging due to the diversity of cysteine reactivity across the proteome.^24^ In contrast, the direct radical labelling of alkene motifs with fluorescent thiol probes remains significantly less explored, despite its clear potential for the visualisation of naturally occurring unsaturations, such as dehydro-amino acids generated by protein aging.^25–27^

Only a limited number of fluorescent thiol probes suitable for alkene-directed TEC have been reported to date. Li *et al.* demonstrated the use of dansyl cysteamine and bi-dansyl cysteamine derivatives for site-selective protein labelling through genetically incorporated alkenes, thereby establishing the feasibility of thiol-based fluorescent ligation in biological systems.^28^ This work achieved selective labelling of alkene containing proteins with ∼50% labelling efficiency. Similarly, Lo Conte *et al.* reported a fluorescein-based thiol system for labelling of alkyne-functionalised monosaccharides and serum albumin conjugates *via* the related thiol-yne reaction, highlighting the wide applicability of sulfur-centred radical ligation strategies in biomolecular settings.^29^ Although these studies elegantly established proof-of-concept, the available probe architectures remain comparatively bulky and, in some instances, synthetically complex, thus limiting broader utility. Moreover, examples that can competitively and efficiently tag unsaturated moieties within complex biological environments, such as cell lysate, remain scarce.^30^ Taken together, these precedents underscore both the considerable promise of thiol-based fluorescent probes and the opportunity to develop more streamlined platforms for selective biomolecular labelling.

Herein, we present a compact, synthetically accessible fluorescent thiol probe capable of selectively targeting alkene moieties in both small molecules and biological systems *via* the radical-mediated TEC reaction. The aminobenzamide-derived thiol probe **1**, termed AbzSH, offers several key advantages: compact size, intrinsic fluorescence and facile synthesis from commercially available precursors. These features render AbzSH particularly attractive as a minimally perturbing labelling reagent for sensitive biomolecular substrates, while retaining the reactivity required for radical-mediated TEC. In practice, this fluorescent probe is suitable for labelling both synthetically and genetically incorporated alkenes, even within complex reaction media such as cell lysates. Thus, the AbzSH probe expands the chemistry toolbox for probing alkene containing biomolecules, allowing for improved understanding of the cellular makeup in disease states.

Initially, we had explored the use of the reported dansyl and fluorescein groups for labelling, given their existing precedent.^28,29^ However, in our initial screening conditions using a model alkene, we observed no reactivity under the conditions screened. As such, we moved to investigate the relatively simple Abz fluorophore, AbzSH **1**, which was synthesised in a good overall yield of 78% across a concise 3-step route, starting from commercially available anthranilic acid (**Scheme S1**). Initial studies focused on evaluating AbzSH reactivity under standard photochemical TEC conditions using a model alkene substrate, a terminal alkene-bearing OAc-glucosamine **2**, herein termed GlcNAc analogue **2** (**Figure 1a**).^31,32^ Initiation was achieved using UV-A irradiation and 2,2-Dimethoxy-2-phenylacetophenone (DPAP) as a photoinitiator. An initial test reaction in DMSO-d6 showed quantitative consumption of the alkene by NMR (**Figure 1b**). The formation of product **4** was also confirmed via HRMS analysis (**Figure 1b**). However, unexpectedly, the TEC reaction failed to proceed in a range of conventional organic solvents, including dichloromethane (DCM), ethyl acetate (EtOAc), acetonitrile (ACN) and dimethylformamide (DMF) (**Figure 1c, Entries 1–4**). In many cases, the probe remained unreacted, while under prolonged irradiation the corresponding disulfide was observed as the major product. Disulfide formation is postulated to have occurred through a radical mechanism,^33,34^ indicating formation of a thiyl radical intermediate. Thus, extensive screening of irradiation wavelength, photoinitiator, reaction time and thiol stoichiometry were explored (**Table S1**). However, this screening process did not result in measurable product formation. Since primary amines are a known inhibitory factor in thiyl radical mediated reactions, we investigated if the aromatic amine of the AbzSH probe was detrimental to the radical chain process.^35^ Therefore, an amine-deficient AbzSH analogue **3** was synthesised; however, when **3** was exposed to the reaction conditions previously explored, a similar lack of reactivity was observed (**Figure 1c, Entry 7**). These findings suggest that the diminished reactivity is unlikely to arise from interference of the amine functionality, and instead may reflect solvent-dependent self-association or aggregation phenomena, potentially driven by aromatic *π* − *π* interactions, which would reduce effective thiol accessibility.^36,37^ This observation led us to pursue subsequent optimisation studies in other solvent systems, where these aggregation effects could be suppressed. Reactivity in purely aqueous conditions could not be achieved due to solubility of the reagents and therefore methanol was incorporated as a suitable co-solvent to fully solubilise the reactants. Implementation of an aqueous solvent system gave rise to formation of product **4** in a low yield of only15%. In an effort to enhance this yield, the use of additives was pursued. Initially, 6 M of Gdn•HCl, a chaotropic agent, was incorporated into the reaction mixture to disrupt hydrogen bonding effects.^38^ This furnished the desired product **4** in 22% yield (**Figure 1c, Entry 5**), indicating that intermolecular effects were indeed influencing the reaction outcome. Further optimisation led to the use of buffered solutions, with reaction in HEPES-buffer/MeOH solvent mixtures offering a markedly improved isolated yield of 80% (**Figure 1c, Entry 6**). Further, reaction of **3** under these optimised conditions furnished the desired product **5** in a yield of 76% (**Figure 1c, Entry 8**). The enhanced performance in water and emerging hypothesis that solvent environment plays a decisive role in probe activity is also consistent with literature precedents for protein thiol–ene ligation, where buffered aqueous conditions are routinely employed for efficient radical-mediated conjugation.^28,29,39,40^

**Figure 1:**
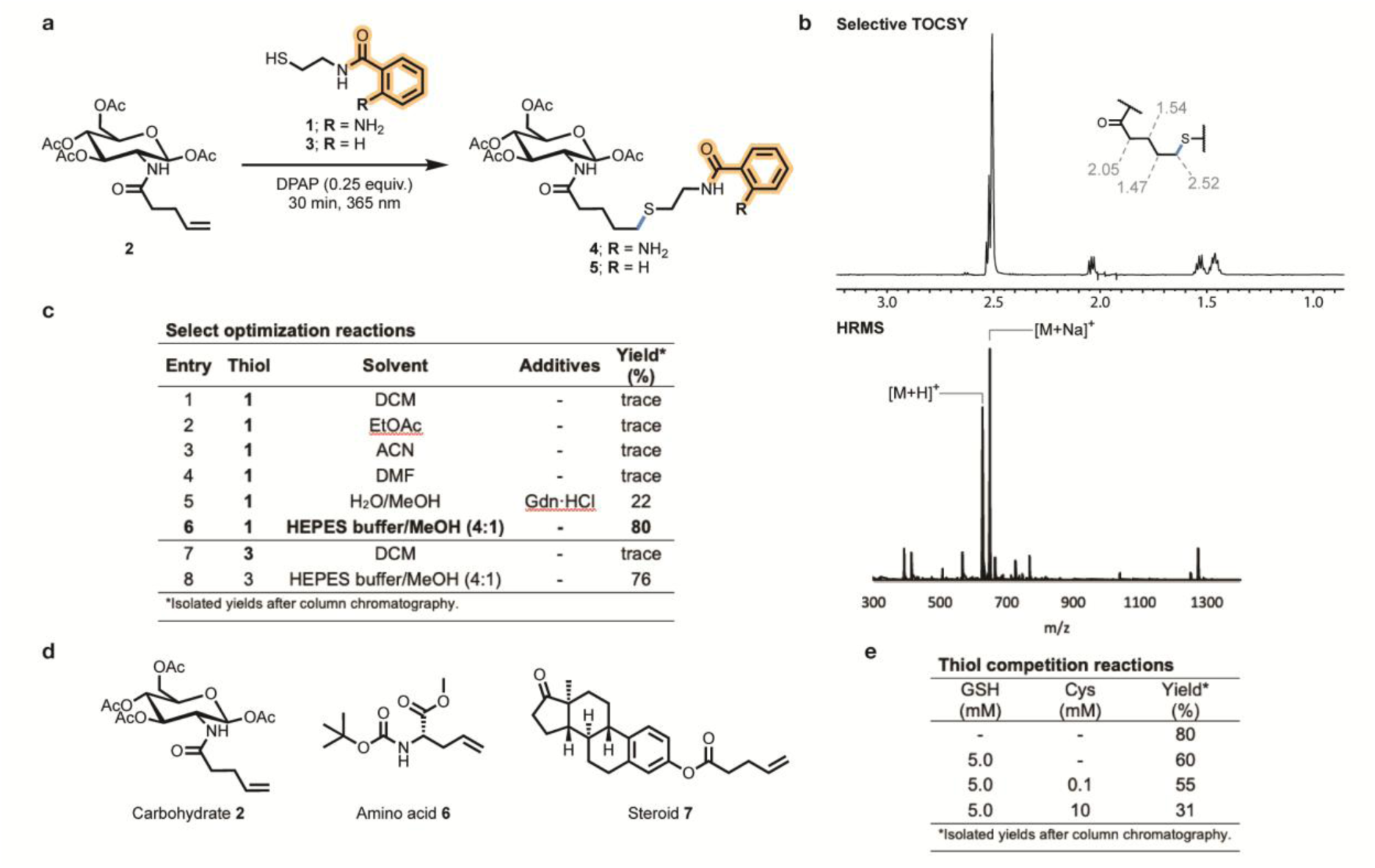
Initial optimisation and investigation of AbzSH labelling of unsaturated biomolecules. (a) AbzSH mediated labelling of GlcNAc analogue **2**. (b) TOCSY NMR and HRMS analysis of the fluorescent-labelled product. (c) Optimisation table for AbzSH labelling. (d) Structures of alkenes labelled using AbzSH. (e) Effect of additional thiols on isolated yield of labelled GlcNAc from AbzSH reaction.

Having successfully labelled GlcNAc analogue **2** to furnish thioether **4** in an isolated yield of 80%, we next sought to expand the substrate scope across a range of structurally diverse alkene-containing biomolecules (**Figure 1d**). Boc-allylglycine methyl ester **6** underwent efficient thiol–ene coupling to furnish the corresponding fluorescent conjugate with an excellent isolated yield of 84%, highlighting the significant potential of the methodology for peptide labelling. Reaction of an alkene functionalised estrone **7** furnished the desired thioether product with only a 10% isolated yield, likely due to solubility challenges associated with the steroidal substrate under the standard conditions. Nevertheless, successful labelling of three major classes of biomolecule highlights the significant potential of the AbzSH thiol–ene platform for fluorescent labelling.

Having demonstrated ligation across small-molecule, biomimetic substrates, we next set out to investigate the capabilities of the AbzSH probe **1** within complex biological environments. A key challenge for thiol-based ligation within biological environments is the presence of native thiol residues, which can act as a competing thiol source.^41,42^ Although thiols are involved in multiple biological processes, concentrations of both protein thiols (PSH), and non-protein thiols (NPSH), are tightly regulated, with single-cell thiol profiling revealing very low quantities of detectable thiols.^43–45^ However, a notable exception is reduced glutathione (GSH), the most abundant intracellular low-molecular-weight thiol, which is typically present at intracellular concentrations between 1-10 mM.^46,47^ GSH has the potential to influence reactions of thiol-based probes through radical quenching or competitive thiol–ene addition, thereby attenuating labelling efficiency.^48^ On the other hand, GSH can partake in chain-transfer processes which may actually enhance efficiency.^49^ Indeed, previous alkene-directed, thiol–ene protein labelling studies have explicitly incorporated glutathione into their reaction systems to mimic intracellular conditions, highlighting its recognised importance in assessing biological compatibility.^48^

To evaluate the tolerance of the AbzSH platform towards biologically relevant thiol concentrations, a control experiment was performed in the presence of glutathione (GSH, 10 mM), representing the upper range of intracellular levels. Using GlcNAc analogue **2** as a model substrate, the desired labelled product was obtained in 60% isolated yield under standard conditions (AbzSH, 10 mM), a modest decrease relative to the 80% yield in the absence of a competing thiol and demonstrating that AbzSH-mediated thiol–ene coupling remains highly effective, even under physiologically relevant GSH concentrations (**Figure 1e**). To further probe chemoselectivity, cysteine was introduced as a second competing thiol. Gratifyingly, at biologically relevant cysteine concentrations (∼100 μM), and in the presence of GSH (10 mM) the AbzSH probe (10 mM) gave a 55% yield of the desired thioether product – indicating minimal additional impact on labelling efficiency. Remarkably, under deliberately stringent conditions (10 mM AbzSH, 10 mM GSH and 10 mM cysteine, a 100-fold increase over physiological cysteine levels), productive labelling was still achieved, affording the desired product in 31% yield. Taken together, these results demonstrate that although endogenous thiols can attenuate selectivity, they do not preclude alkene labelling by AbzSH, which outcompetes the screened endogenous thiols. Importantly, for practical protein-labelling applications, the probe can be employed in excess relative to the alkene-containing target, further favouring productive ligation in thiol-rich environments. This observation strongly supports the successful application of AbzSH in lysate-based and cellular settings and enhances the biorthogonal credentials of thiol–ene ligation.

We next sought to evaluate the AbzSH probe for protein labelling, where selective ligation in complex macromolecular environments is considerably more challenging. As an initial proof-of-concept, bovine serum albumin (BSA) was selected as a model protein scaffold due to its well-characterised structure and multiple, solvent-accessible lysine residues. To generate an alkene-bearing model protein, the 36 solvent-exposed lysine residues of BSA were chemically acylated using alkene-functionalised *N*-hydroxysuccinimide ester **S5**, affording alkene-BSA C. Two control proteins were prepared in parallel: an alkane-modified analogue (alkane-BSA B), obtained using the corresponding saturated NHS ester **S6**, and an unmodified BSA control (BSA A), generated by treatment with DMSO in the absence of an NHS ester reagent.

Fluorescent labelling of the three BSA variants was investigated under photochemical TEC conditions using DPAP (0.7 mM) as a photoinitiator with varying concentrations of AbzSH probe (2.3 µM–2.3 mM). Reactions were performed in aqueous buffer, akin to the optimal solvent systems previously identified. 4′-methoxyacetophenone (MAP, 4.9 mM) was included as a photosensitiser to increase the rate of the radical reaction.^50^ Following UV irradiation for 5 minutes, labelling efficiency was assessed by in-gel fluorescence at 302 nm. A weak fluorescence signal was observed for alkene-BSA treated with 1 mM AbzSH, while increasing the probe concentration to 2.3 mM resulted in a clear and intense fluorescence band, consistent with successful covalent labelling of the alkene-functionalised protein (**Figure 2, lane 6**). In contrast, neither unmodified BSA A nor alkane-BSA B displayed detectable fluorescence under identical conditions (**Figure 2, lanes 4 and 5**), strongly indicating that probe attachment proceeds selectively through thiol–ene reaction with the alkene functionality, rather than through non-specific protein modification.

**Figure 2:**
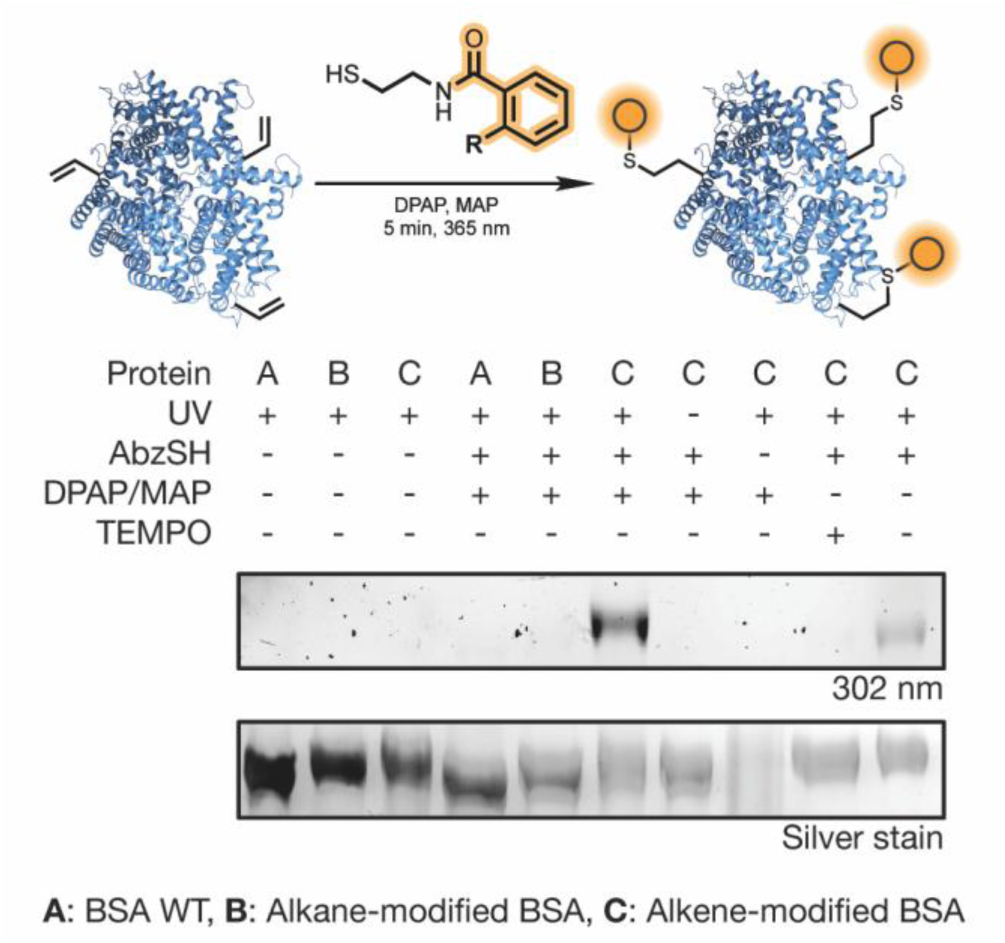
TEC labelling of alkene modified BSA variants (4 μg) with AbzSH. Assay visualised by silver staining and fluorescent imaging at 302 nm. Stability of proteins A, B and C to UV irradiation confirmed (lanes 1-3) and alkene-free proteins A and B tested as negative controls (lanes 4,5). Clear AbzSH labelling observed in lane 6. Sequential removal of reaction components (lanes 7-10) confirms a radical mechanism.

To further validate the mode of labelling, a series of mechanistic control experiments were performed using alkene-BSA. No fluorescent signal was detected in the absence of UV irradiation (**Figure 2, lane 7**), confirming that light activation is essential for productive conjugation. Inclusion of the radical scavenger TEMPO fully inhibited labelling, indicating that the reaction indeed proceeds *via* a radical-mediated reaction mechanism (**Figure 2, lane 9**). Interestingly, weak labelling was still observed under UV exposure in the absence of DPAP and MAP (**Figure 2, lane 10**), suggesting that the thiyl radical can be readily generated directly upon irradiation, potentially *via* atmospheric oxygen mediated initiation.^51^ As expected, no labelling was observed in the absence of AbzSH (**Figure 2, lane 8**). Under these conditions, however, some protein degradation was apparent. This was not the case in samples that did not contain DPAP and MAP (**Figure 2, lanes 1-3**). Protein degradation was likely induced by carbon-centred radical species generated by DPAP photolysis.^22,52^

Although chemically modified BSA provided an effective model protein, a homogeneous protein system displaying a single, defined alkene handle was required to more rigorously assess the chemoselectivity and site-specificity of AbzSH as a fluorescent probe. We employed genetic code expansion to site-selectively incorporate an alkene-containing non-canonical amino acid (ncAA) into a model protein scaffold *via* amber codon suppression.^53–55^ Commercially available *S*-allyl-cysteine (SAC) was selected based on previous reports by Budisa *et al.*, who demonstrated its compatibility with orthogonal translation machinery.^56^ SAC was then incorporated into Lactococcal multidrug resistance Regulator (LmrR).

Four different synthetase/tRNA orthogonal pairs (OPs) were screened for their ability to selectively incorporate SAC. OP2 was identified as the most effective synthetase/tRNA pair, exhibiting a strong concentration-dependent sfGFP fluorescence response, indicative of successful SAC incorporation (**SI Figure S1**). Using site-directed mutagenesis, an amber stop codon (TAG) was then introduced into wild-type LmrR constructs at the solvent-exposed V15 position. Large-scale protein expression was carried out in E. Coli BL21 (DE3). LmrR V15SAC was successfully expressed with a yield of 14 mg L^-1^ and successful ncAA incorporation was confirmed by mass spectrometry (**SI Figure S2**).

Fluorescent labelling of LmrR V15SAC with AbzSH was subsequently performed under conditions analogous to those employed for alkene-BSA, with increased protein loading to allow for the presence of only a single alkene residue per protein molecule. Gratifyingly, a clear fluorescence signal was observed only when all reaction components were present (**Figure 3a, lane 10**), confirming successful, site-selective protein labelling. No signal was detected in the absence of either photoinitiator cocktail or UV irradiation (**Figure 3a, lanes 8 and 9**), demonstrating that productive ligation requires full TEC activation. Importantly, unmodified wild-type LmrR remained unlabelled under identical conditions (**Figure 3a, lanes 4 and 5**), confirming the excellent chemoselectivity of AbzSH towards the genetically installed alkene handle and highlighting that non-specific labelling of the protein backbone does not occur under these conditions. Gratifyingly, the relative intensity of the protein bands remained constant under all reaction conditions screened (**SI Figure S5 and S6**), in contrast to the results observed utilising modified BSA, which displayed more variable reactivity due to the presence of 36 distinct reactive sites.

**Figure 3:**
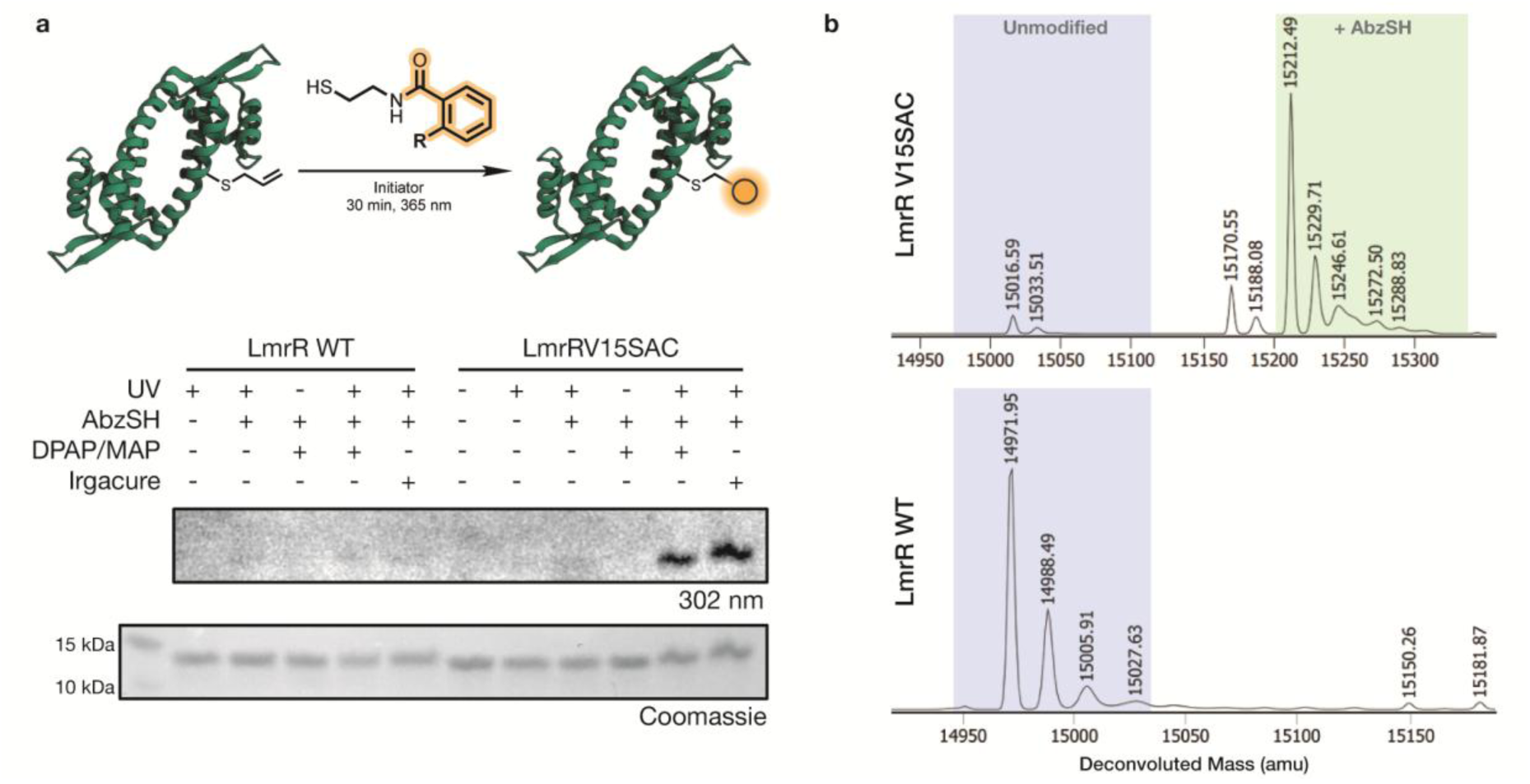
TEC initiated by DPAP/MAP and Irgacure photoinitiators shown in lanes 10 and 11 respectively. TEC labelling of LmrR V15SAC (6 μg) with AbzSH (2 mM). (a) Assay visualised by Coomassie Blue staining and fluorescent imaging at 305 nm. Wild type LmrR used as a negative control (lanes 1-5). Sequential removal of reaction components, tested against LmrR V15SAC (lanes 6-9) confirms a radical mechanism. TEC initiated by DPAP/MAP and Irgacure photoinitiators shown in lanes 10 and 11 respectively. (b) MS analysis of products of labelling reactions on LmrRV15SAC substrate and LmrR WT non-substrate.

In addition to SDS-PAGE and in-gel fluorescence, the success of the ligation reaction and the stoichiometry of probe attachment was also determined by LC-MS analysis (**Figure 3b**), revealing near-complete conversion of LmrRV15SAC with attachment of a single AbzSH moiety per protein molecule, providing compelling evidence for precise and stoichiometric alkene-directed ligation. No conversion was detected on unmodified LmrR under identical conditions. Although the DPAP/MAP photoinitiator system consistently delivered efficient fluorescent labelling, we sought to further investigate if the AbzSH-mediated TEC platform was compatible with alternative radical initiation systems. Broad initiator compatibility would significantly enhance the practical utility of the method, particularly for biological applications where solvent compatibility and phototoxicity must be carefully considered. To this end, water-soluble Irgacure 2959 was investigated as an alternative photoinitiator under otherwise identical conditions to label LmrRV15SAC.^57^ Gratifyingly, clear fluorescent labelling of LmrRV15SAC was observed by in-gel fluorescence analysis, with no signal detected for unmodified LmrR under the same conditions (**Figure 3, lanes 5 and 11**). LC-MS analysis confirmed covalent attachment of a single AbzSH moiety to the protein, consistent with site-specific alkene ligation (**Figure S3**). Although minor low-intensity peaks corresponding to unidentified side products were observed in the mass spectrum, these did not correspond to AbzSH or any known Irgacure-derived adducts. Taken together, these findings demonstrate that the AbzSH probe is compatible with alternate radical initiation platforms, highlighting the broader methodological flexibility of this alkene-labelling strategy.

Following successful site-selective protein labelling, we next set out to further optimise reagent loading against our genetically expressed model to improve efficiency, while minimising probe consumption and limiting unnecessary exposure of the protein to radical forming conditions. A concentration screening was therefore undertaken, using AbzSH concentrations ranging from 0.1 to 2.0 mM, with DPAP and MAP maintained in proportional quantities and all other reaction parameters held constant (**Figure S5**). In-gel fluorescence analysis revealed a clear dependence of labelling intensity on reagent concentration, with progressively stronger fluorescence signals observed at higher probe loadings. Importantly, labelling remained detectable even at 0.1 mM AbzSH, the lowest concentration tested, demonstrating the high sensitivity of the system and the intrinsic efficiency of alkene capture by the probe. This result is particularly significant from a chemical biology perspective, as it demonstrates that effective labelling can be achieved using relatively modest probe concentrations, a key feature for fluorescent labelling strategies both *in vitro* and *in vivo*.

To further refine the labelling conditions, a two-dimensional optimisation study was performed in which both probe concentration and irradiation time were systematically varied (**Figure S6**). AbzSH concentrations of 0.2, 0.5, 1.0, and 2.0 mM were evaluated in combination with UV irradiation times ranging from 2.5 to 30 minutes. Encouragingly, detectable labelling was observed after only 5 minutes across all tested concentrations, underscoring the rapid kinetics of the radical-mediated TEC process.^58–60^ At 1.0 and 2.0 mM AbzSH, maximal fluorescence intensity was achieved after 10 minutes of irradiation, whereas 0.5 mM required 30 minutes to reach comparable signal intensity. At the lowest concentration tested (0.2 mM), signal intensity continued to increase beyond 30 minutes, indicating incomplete reaction under these conditions. On the basis of these studies, the optimal protein labelling conditions were identified as 1.0 mM AbzSH, 0.4 mM DPAP, and 2.5 mM MAP with 10 minutes of UV irradiation. These conditions offered an excellent balance between reagent economy, reaction efficiency, and minimisation of photodamage.

Having established robust labelling conditions in purified protein systems, we next evaluated whether the AbzSH platform could function in a more biologically complex environment and in the presence of both endogenous thiols and alkenes. Cell-free E. coli lysates expressing LmrRV15SAC were prepared to provide a heterogeneous, protein-rich milieu, representative of biological conditions.^61,62^ While the purpose of this experiment is to investigate probe reactivity in a complex lysate, it can also serve as a readout for confirmation of successful incorporation of the ncAA within the target protein during expression. Under the optimised conditions, efficient fluorescent labelling of LmrRV15SAC was observed across all tested AbzSH concentrations (0.5, 1.0, and 2.0 mM), with the strongest signal obtained at 1.0 mM, in excellent agreement with the purified protein optimisation studies (**Figure 4, lanes 8, 9, 10 and 11**). Importantly, Coomassie staining revealed no evidence of global protein degradation following irradiation, indicating that the reaction conditions are well tolerated even in complex lysate environments. The ability to selectively label alkene-containing proteins in crude lysate strongly supports the practical utility of AbzSH as a chemical biology probe, particularly for proteomic profiling and functional screening applications.

**Figure 4:**
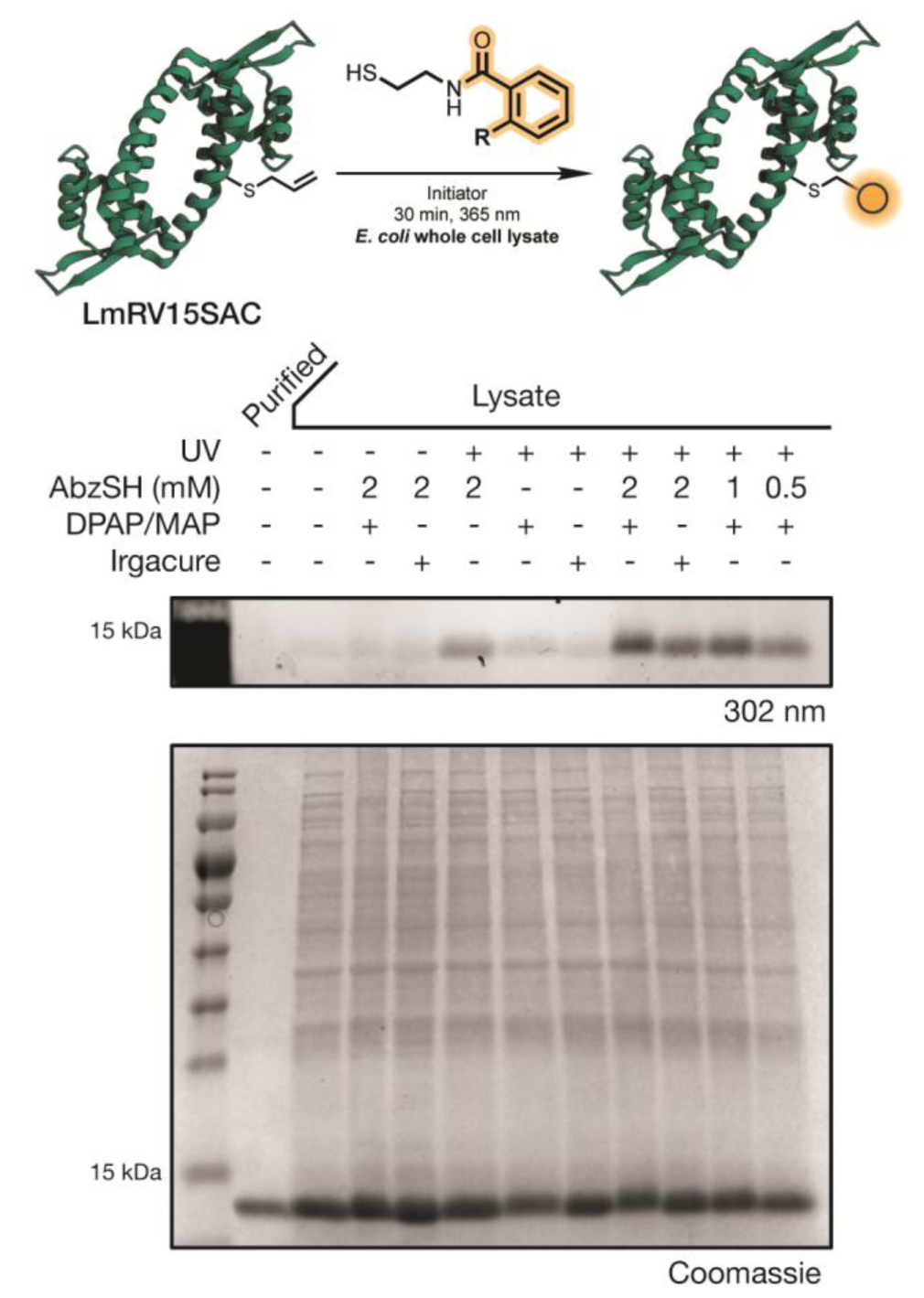
TEC labelling of cell lysate containing LmrR V15SAC with AbzSH. Assay visualised by Coomassie Blue staining and fluorescent imaging at 305 nm. Unlabelled LmrR V15SAC shown in lane 1 for reference with lanes 2-4 and 6-7 providing negative controls. UV only, DPAP/MAP or irgacure initiation shown in lane 5 and 8-11 with varying concentrations of AbzSH.

In conclusion, we report a novel and readily accessible fluorescent thiol probe, AbzSH, for the selective labelling of alkene-containing biomolecules *via* the radical-mediated thiol–ene ‘click’ (TEC) reaction. The AbzSH probe was shown to efficiently label a diverse range of non-native alkene functionalities on biomolecular scaffolds, including carbohydrate, steroid and amino acid substrates, and displayed high tolerance to physiologically relevant glutathione competition. The utility of the probe was further extended to biologically relevant protein systems, including proteins bearing chemically and genetically incorporated alkenes, both of which were successfully labelled with AbzSH. Under optimised aqueous photochemical conditions, near-complete conversion of LmrRV15SAC was achieved with selective attachment of a single AbzSH moiety per protein, as confirmed by in-gel fluorescence and LC–MS analysis. This demonstrates the remarkable efficacy and selectivity of AbzSH, with only 1.0 mM of thiol label and 10 min reaction times required under optimal conditions. Crucially, these conditions remained effective in complex cell lysate, demonstrating the robustness and chemoselectivity of the platform in highly complex, biologically relevant environments. Taken together, this operationally simple and highly efficient methodology establishes AbzSH as a versatile fluorescent labelling platform for probing unsaturated residues in complex biological systems. We anticipate that this approach will provide a valuable addition to the chemical biology toolbox, with potential for applications in biomolecular imaging, proteomics, biological screening, and future drug discovery efforts.

## ASSOCIATED CONTENT

### Supporting Information

The Supporting Information is available free of charge.

Experimental sections, analytical characterisation data (NMR spectra), biological procedures, additional optimisation data and supplementary figures. (PDF)

## AUTHOR INFORMATION

### Author Contributions

The manuscript was written through contributions of all authors. All authors have given approval to the final version of the manuscript.

## Noetes

## ACKNOWLEDGMENT

We are grateful to Dr John O’Brien and Dr Manuel Ruether for assistance with NMR spectroscopy and Dr Gary Hessman for assistance with mass spectrometry.

## ABBREVIATIONS

AbzSH: 2-amino-*N*-(2-mercaptoethyl)benzamide;
BSA: bovine serum albumin;
DPAP: 2,2-dimethoxy-2-phenylacetophenone;
Gdn•HCl: guanidine hydrochloride;
GSH: glutathione;
HEPES: 4-(2-hydroxyethyl)-1-piperazineethanesulfonic acid;
LmrR: lactococcal multidrug resistance regulator;
MAP: 4′-methoxyacetophenone;
MeOH: methanol;
ncAA: non-canonical amino acid;
NHS: *N*-hydroxysuccinimide;
NPSH: non-protein thiols;
PSH: protein thiols;
SAC: *S-*allyl-cysteine;
TEC: thiol–ene ‘click’;
TEMPO: 2,2,6,6-Tetramethylpiperidin-1-oxyl.

## References

(1) Jeon, S.; Jeon, Y.; Lim, J.-Y.; Kim, Y.; Cha, B.; Kim, W. Emerging Regulatory Mechanisms and Functions of Biomolecular Condensates: Implications for Therapeutic Targets. Signal Transduct. Target. Ther. 2025, 10 (1), 4. 10.1038/s41392-024-02070-1

(2) Han, B.; Li, B.; Wang, X.; Wang, L. Biomolecular Condensates in Disease: Decoding the Material State and Engineering Precision Modulators. Int. J. Mol. Sci. 2026, 27 (2), 837. 10.3390/ijms27020837

(3) Akter, R.; Cao, P.; Noor, H.; Ridgway, Z.; Tu, L.-H.; Wang, H.; Wong, A. G.; Zhang, X.; Abedini, A.; Schmidt, A. M.; Raleigh, D. P. Islet Amyloid Polypeptide: Structure, Function, and Pathophysiology. J. Diabetes Res. 2016, 2016 (1), 2798269. 10.1155/2016/2798269

(4) Moya-Gudiño, V.; Altamirano-Bustamante, N. F.; Revilla-Monsalve, C.; Altamirano-Bustamante, M. M. Decoding the Contribution of IAPP Amyloid Aggregation to Beta Cell Dysfunction: A Systematic Review and Epistemic Meta-Analysis of Type 1 Diabetes. Int. J. Mol. Sci. 2025, 26 (2), 767. 10.3390/ijms26020767

(5) Specht, E. A.; Braselmann, E.; Palmer, A. E. A Critical and Comparative Review of Fluorescent Tools for Live-Cell Imaging. Annu. Rev. Physiol. 2017, 79, 93–117. 10.1146/annurev-physiol-022516-034055

(6) Choi, N.-E.; Lee, J.-Y.; Park, E.-C.; Lee, J.-H.; Lee, J. Recent Advances in Organelle-Targeted Fluorescent Probes. Molecules 2021, 26 (1), 217. 10.3390/molecules26010217

(7) Knaack, J. I. H.; Meier, C. Out of the Dark, into the Light: Metabolic Fluorescent Labeling of Nucleic Acids. ChemMedChem 2024, 19 (15), e202400160. 10.1002/cmdc.202400160

(8) Shimogawa, M.; Petersson, E. J. New Strategies for Fluorescently Labeling Proteins in the Study of Amyloids. Curr. Opin. Chem. Biol. 2021, 64, 57–66. 10.1016/j.cbpa.2021.04.011

(9) Sletten, E. M.; Bertozzi, C. R. Bioorthogonal Chemistry: Fishing for Selectivity in a Sea of Functionality. Angew. Chem. Int. Ed. 2009, 48 (38), 6974–6998. 10.1002/anie.200900942

(10) Li, Y.; Fu, H. Bioorthogonal Ligations and Cleavages in Chemical Biology. ChemistryOpen 2020, 9 (8), 835–853. 10.1002/open.202000128

(11) Oliveira, B. L.; Guo, Z.; Bernardes, G. J. L. Inverse Electron Demand Diels–Alder Reactions in Chemical Biology. Chem. Soc. Rev. 2017, 46 (16), 4895–4950. 10.1039/C7CS00184C

(12) Scache, J.; Rigolot, V.; Lion, C.; Mortuaire, M.; Lefebvre, T.; Biot, C.; Vercoutter-Edouart, A.-S. Switching Azide and Alkyne Tags on Bioorthogonal Reporters in Metabolic Labeling of Sialylated Glycoconjugates: A Comparative Study. Sci. Rep. 2022, 12 (1), 22129. 10.1038/s41598-022-26521-3

(13) George, J. T.; Srivatsan, S. G. Bioorthogonal Chemistry-Based RNA Labeling Technologies: Evolution and Current State. Chem. Commun. 2020, 56 (82), 12307–12318. 10.1039/d0cc05228k

(14) Amiri, A.; Abedanzadeh, S.; Khodabandehloo, M. H.; Shaabani, A.; Moosavi-Movahedi, A. A. Ligand Tricks for Faster Clicks: Bioconjugation via the CuAAC Reaction. Chem. Commun. 2025, 61 (100), 19810–19827. 10.1039/d5cc04435a

(15) Hoyle, C. E.; Bowman, C. N. Thiol–Ene Click Chemistry. Angew. Chem. Int. Ed. 2010, 49 (9), 1540–1573. 10.1002/anie.200903924

(16) Povie, G.; Tran, A.-T.; Bonnaffé, D.; Habegger, J.; Hu, Z.; Le Narvor, C.; Renaud, P. Repairing the Thiol-Ene Coupling Reaction. Angew. Chem. Int. Ed. 2014, 53 (15), 3894–3898. 10.1002/anie.201309984

(17) McLean, J. T.; Benny, A.; Nolan, M. D.; Swinand, G.; Scanlan, E. M. Cysteinyl Radicals in Chemical Synthesis and in Nature. Chem. Soc. Rev. 2021, 50 (19), 10857–10894. 10.1039/D1CS00254F

(18) Nolan, M. D.; Scanlan, E. M. Applications of Thiol-Ene Chemistry for Peptide Science. Front. Chem. 2020, 8, 583272. 10.3389/fchem.2020.583272

(19) Prieto-Castañeda, A.; Martin, H.; Manna, T.; Beswick, L.; McLean, J. T.; Pongener, I.; González, I. R.; Twamley, B.; Miller, G. J.; Scanlan, E. M. Photoinitiated Thiol–Ene Mediated Functionalisation of 4,5-Enoses. Org. Biomol. Chem. 2025, 23 (22), 5332–5338. 10.1039/D5OB00507H

(20) Choi, H.; Kim, M.; Jang, J.; Hong, S. Visible-Light-Induced Cysteine-Specific Bioconjugation: Biocompatible Thiol–Ene Click Chemistry. Angew. Chem. Int. Ed. 2020, 59 (50), 22514–22522. 10.1002/anie.202010217

(21) Lechner, V. M.; Nappi, M.; Deneny, P. J.; Folliet, S.; Chu, J. C. K.; Gaunt, M. J. Visible-Light-Mediated Modification and Manipulation of Biomacromolecules. Chem. Rev. 2022, 122 (2), 1752–1829. 10.1021/acs.chemrev.1c00357

(22) Campaniҫo, A.; Baran, M.; Bowie, A. G.; Longley, D. B.; Harrison, T.; McGouran, J. F. Chemical-and Photo-Activation of Protein-Protein Thiol-Ene Coupling for Protein Profiling. Commun. Chem. 2025, 8 (1), 25. 10.1038/s42004-025-01412-6

(23) Bellaizac-Riascos, J.; Ortiz-Serna, P.; Pérez-Ruiz, R.; Encinas, S.; Jiménez, M. C. Synthesis of Cysteine–Fluorophore Conjugates by Visible-Light-Promoted Thiol–Ene Coupling Using Organic Photocatalysts. Eur. J. Org. Chem. 2026, 29 (38), e70823. 10.1002/ejoc.70823

(24) Weerapana, E.; Wang, C.; Simon, G. M.; Richter, F.; Khare, S.; Dillon, M. B. D.; Bachovchin, D. A.; Mowen, K.; Baker, D.; Cravatt, B. F. Quantitative Reactivity Profiling Predicts Functional Cysteines in Proteomes. Nature 2010, 468 (7325), 790–795. 10.1038/nature09472

(25) Wang, Z.; Lyons, B.; Truscott, R. J. W.; Schey, K. L. Human Protein Aging: Modification and Crosslinking through Dehydroalanine and Dehydrobutyrine Intermediates. Aging Cell 2014, 13 (2), 226–234. 10.1111/acel.12164

(26) Markovich, S. W.; Frey, B. L.; Scalf, M.; Shortreed, M. R.; Smith, L. M. Dehydroamino Acids and Their Crosslinks in Alzheimer’s Disease Aggregates. Brain Commun. 2025, 7 (1), fcaf019. 10.1093/braincomms/fcaf019

(27) Jones, L. H. Dehydroamino Acid Chemical Biology: An Example of Functional Group Interconversion on Proteins. *RSC Chem*. Biol. 2020, 1 (5), 298–304. 10.1039/d0cb00174k

(28) Li, Y.; Yang, M.; Huang, Y.; Song, X.; Liu, L.; Chen, P. R. Genetically Encoded Alkenyl–Pyrrolysine Analogues for Thiol–Ene Reaction Mediated Site-Specific Protein Labeling. Chem. Sci. 2012, 3 (9), 2766–2770. 10.1039/C2SC20433A

(29) Conte, M. L.; Staderini, S.; Marra, A.; Sanchez-Navarro, M.; Davis, B. G.; Dondoni, A. Multi-Molecule Reaction of Serum Albumin Can Occur through Thiol-Yne Coupling. Chem. Commun. 2011, 47 (39), 11086–11088. 10.1039/C1CC14402B

(30) Yang, T.; Tang, S.; Kuo, S.-T.; Freitas, D.; Edwards, M.; Wang, H.; Sun, Y.; Yan, X. Lipid Mass Tags via Aziridination for Probing Unsaturated Lipid Isomers and Accurate Relative Quantification. Angew. Chem. Int. Ed. 2022, 61 (39), e202207098. 10.1002/anie.202207098

(31) Nolan, M. D.; Schüttel, M.; Scanlan, E. M.; Nielsen, A. L. Nanomole-Scale Photochemical Thiol-Ene Chemistry for High-Throughput Late-Stage Diversification of Peptide Macrocycles. Pept. Sci. 2024, 116 (1), e24310. 10.1002/pep2.24310

(32) Nolan, M. D.; Mezzetta, A.; Guazzelli, L.; Scanlan, E. M. Radical-Mediated Thiol–Ene ‘Click’ Reactions in Deep Eutectic Solvents for Bioconjugation. Green Chem. 2022, 24 (4), 1456–1462. 10.1039/D1GC03714E

(33) Izmailov, S. A.; Podkorytov, I. S.; Skrynnikov, N. R. Simple MD-Based Model for Oxidative Folding of Peptides and Proteins. Sci. Rep. 2017, 7 (1), 9293. 10.1038/s41598-017-09229-7

(34) Winterbourn, C. C.; Hampton, M. B. Thiol Chemistry and Specificity in Redox Signaling. Free Radic. Biol. Med. 2008, 45 (5), 549–561. 10.1016/j.freeradbiomed.2008.05.004

(35) Love, D. M.; Kim, K.; Goodrich, J. T.; Fairbanks, B. D.; Worrell, B. T.; Stoykovich, M. P.; Musgrave, C. B.; Bowman, C. N. Amine Induced Retardation of the Radical-Mediated Thiol–Ene Reaction via the Formation of Metastable Disulfide Radical Anions. J. Org. Chem. 2018, 83 (5), 2912–2919. 10.1021/acs.joc.8b00143

(36) Bera, S.; Dong, X.; Krishnarjuna, B.; Raab, S. A.; Hales, D. A.; Ji, W.; Tang, Y.; Shimon, L. J. W.; Ramamoorthy, A.; Clemmer, D. E.; Wei, G.; Gazit, E. Solid-State Packing Dictates the Unexpected Solubility of Aromatic Peptides. Cell Rep. Phys. Sci. 2021, 2 (4), 100391. 10.1016/j.xcrp.2021.100391

(37) Heo, J.; Murale, D. P.; Yoon, H. Y.; Arun, V.; Choi, S.; Kim, E.; Lee, J.-S.; Kim, S. Recent Trends in Molecular Aggregates: An Exploration of Biomedicine. Aggregate 2022, 3 (2), e159. 10.1002/agt2.159

(38) Benny, A.; Scanlan, E. M. Synthesis of Macrocyclic Thiolactone Peptides *via* Photochemical Intramolecular Radical Acyl Thiol–Ene Ligation. Chem. Commun. 2024, 60 (61), 7950–7953. 10.1039/D4CC02442G

(39) Hurst, R. D.; Nieves, A.; Brichacek, M. Expanding Glycomic Investigations through Thiol-Derivatized Glycans. Molecules 2023, 28 (4), 1956. 10.3390/molecules28041956

(40) Williams, C.; Raabe, K.; Tully, D.; Windisch, D.; O’Brien, J.; Trujillo, C.; McGouran, J. F.; Muttenthaler, M.; Scanlan, E. M. Synthesis of Macrocyclic Peptides via Photochemical Radical Thiol–Yne Reaction. J. Am. Chem. Soc. 2026, 148 (31), 33399–33410. 10.1021/jacs.6c08113

(41) Su, J. Thiol-Mediated Chemoselective Strategies for In Situ Formation of Hydrogels. Gels 2018, 4 (3), 72. 10.3390/gels4030072

(42) Cianfoni, G.; Pisano, L.; Varouhaki, D. M.; Centioni, G.; Calcaterra, A.; Ghirga, F.; Athanassopoulos, C. M.; Botta, B.; Cammarone, S.; Baiocco, P.; Quaglio, D. Tunable Linkers for Dynamic Thiol-Based Bioconjugation Strategies. Biomacromolecules 2026, 27 (2), 1024–1072. 10.1021/acs.biomac.5c02149

(43) Yang, F.; Xiao, Y.; Ding, J.-H.; Jin, X.; Ma, D.; Li, D.-Q.; Shi, J.-X.; Huang, W.; Wang, Y.-P.; Jiang, Y.-Z.; Shao, Z.-M. Ferroptosis Heterogeneity in Triple-Negative Breast Cancer Reveals an Innovative Immunotherapy Combination Strategy. Cell Metab. 2023, 35 (1), 84–100. 10.1016/j.cmet.2022.09.021

(44) Yamamoto, H.; Fujiwara, T.; Funatsu, T.; Tsunoda, M. Quantification of Intracellular Thiols by HPLC-Fluorescence Detection. Molecules 2021, 26 (8), 2365. 10.3390/molecules26082365

(45) Miao, D.; Li, Q.; Zhang, Y.; Qin, S.; Wang, Y.; Liu, X.; Bai, Y. Single-Cell Thiol Profiling Enabled by Live-Cell Labeling Reveals Metabolic Heterogeneity in Ferroptosis. Nat. Commun. 2026, 17 (1), 3378. 10.1038/s41467-026-70336-z

(46) Raj Rai, S.; Bhattacharyya, C.; Sarkar, A.; Chakraborty, S.; Sircar, E.; Dutta, S.; Sengupta, R. Glutathione: Role in Oxidative/Nitrosative Stress, Antioxidant Defense, and Treatments. ChemistrySelect 2021, 6 (18), 4566–4590. 10.1002/slct.202100773

(47) Franco, R.; Cidlowski, J. A. Apoptosis and Glutathione: Beyond an Antioxidant. Cell Death Differ. 2009, 16 (10), 1303–1314. 10.1038/cdd.2009.107

(48) Healy, J.; Rasmussen, T.; Miller, S.; Booth, I. R.; Conway, S. J. The Photochemical Thiol–Ene Reaction as a Versatile Method for the Synthesis of Glutathione S-Conjugates Targeting the Bacterial Potassium Efflux System Kef. Org. Chem. Front. 2016, 3 (4), 439–446. 10.1039/C5QO00436E

(49) Venneti, N. M.; Samala, G.; Morsy, R. M. I.; Mendoza, L. G.; Isidro-Llobet, A.; Tom, J. K.; Mukherjee, S.; Kopach, M. E.; Stockdill, J. L. Phosphine-Dependent Photoinitiation of Alkyl Thiols under Near-UV Light Facilitates User-Friendly Peptide Desulfurization. J. Am. Chem. Soc. 2023, 145 (2), 1053–1061. 10.1021/jacs.2c10625

(50) Durchschlag, H.; Fochler, C.; Feser, B.; Hausmann, S.; Seroneit, T.; Swientek, M.; Swoboda, E.; Winklmair, A.; Wlček, C.; Zipper, P. Effects of X-and UV-Irradiation on Proteins. Radiat. Phys. Chem. 1996, 47 (3), 501–505. 10.1016/0969-806X(95)00138-N

(51) Dénès, F.; Pichowicz, M.; Povie, G.; Renaud, P. Thiyl Radicals in Organic Synthesis. Chem. Rev. 2014, 114 (5), 2587–2693. 10.1021/cr400441m

(52) Xu, H.; Casillas, J.; Krishnamoorthy, S.; Xu, C. Effects of Irgacure 2959 and Lithium Phenyl-2,4,6-Trimethylbenzoylphosphinate on Cell Viability, Physical Properties, and Microstructure in 3D Bioprinting of Vascular-like Constructs. Biomed. Mater. 2020, 15 (5), 055021. 10.1088/1748-605X/ab954e

(53) Noren, C. J.; Anthony-Cahill, S. J.; Griffith, M. C.; Schultz, P. G. A General Method for Site-Specific Incorporation of Unnatural Amino Acids into Proteins. Science 1989, 244 (4901), 182–188. 10.1126/science.2649980

(54) Dumas, A.; Lercher, L.; Spicer, C. D.; Davis, B. G. Designing Logical Codon Reassignment – Expanding the Chemistry in Biology. Chem. Sci. 2014, 6 (1), 50–69. 10.1039/C4SC01534G

(55) Huang, Y.; Zhang, P.; Wang, H.; Chen, Y.; Liu, T.; Luo, X. Genetic Code Expansion: Recent Developments and Emerging Applications. Chem. Rev. 2025, 125 (2), 523–598. 10.1021/acs.chemrev.4c00216

(56) Koch, N. G.; Goettig, P.; Rappsilber, J.; Budisa, N. Engineering Pyrrolysyl-tRNA Synthetase for the Incorporation of Non-Canonical Amino Acids with Smaller Side Chains. Int. J. Mol. Sci. 2021, 22 (20), 11194. 10.3390/ijms222011194

(57) Rabadán González, I.; McLean, J. T.; Ostrovitsa, N.; Fitzgerald, S.; Mezzetta, A.; Guazzelli, L.; O’Shea, D. F.; Scanlan, E. M. A Thiol–Ene Mediated Approach for Peptide Bioconjugation Using ‘Green’ Solvents under Continuous Flow. Org. Biomol. Chem. 2024, 22 (11), 2203–2210. 10.1039/d4ob00122b

(58) Northrop, B. H.; Coffey, R. N. Thiol–Ene Click Chemistry: Computational and Kinetic Analysis of the Influence of Alkene Functionality. J. Am. Chem. Soc. 2012, 134 (33), 13804–13817. 10.1021/ja305441d

(59) Höller, R. J.; Sivun, D.; Gvindzhiliia, G.; Müller, S. M.; Haiden, L.; Klar, T. A.; Jacak, J.; Schlögl, S.; Griesser, T. Spatial Control of Curing Kinetics in Thiol-Ene-Systems through Antagonistic Photoreactions. Nat. Commun. 2025, 16 (1), 8487. 10.1038/s41467-025-63407-0

(60) Love, D.; Fairbanks, B.; Bowman, C. Reaction Environment Effect on the Kinetics of Radical Thiol–Ene Polymerizations in the Presence of Amines and Thiolate Anions. ACS Macro Lett. 2020, 9 (2), 174–179. 10.1021/acsmacrolett.9b00960

(61) Ishihama, Y.; Schmidt, T.; Rappsilber, J.; Mann, M.; Hartl, F. U.; Kerner, M. J.; Frishman, D. Protein Abundance Profiling of the Escherichia Coli Cytosol. BMC Genomics 2008, 9, 102. 10.1186/1471-2164-9-102

(62) Soufi, B.; Krug, K.; Harst, A.; Macek, B. Characterization of the E. Coli Proteome and Its Modifications during Growth and Ethanol Stress. Front. Microbiol. 2015, 6, 103. 10.3389/fmicb.2015.00103

