## Supplementary Information for "Rational Design of a Photoactivated Fluorescent Probe for Thiol-ene Mediated Protein Labelling"

### 1. Supplementary Discussion

The AbzSH probe **1** was synthesised through a concise multi-step route from commercially available anthranilic acid. Boc protection of the free amine **S1** to furnish **S2** was achieved quantitatively, followed by a simple amide coupling to give protected fluorophore **S3** in 84% yield. Global deprotection of both protecting groups yielded the final free-thiol fluorophore **1** quantitatively. This synthetic route afforded the target fluorescent thiol in an excellent overall yield of 78% across 3 steps, avoiding the need for complex chemistries (***SI Scheme 1***).

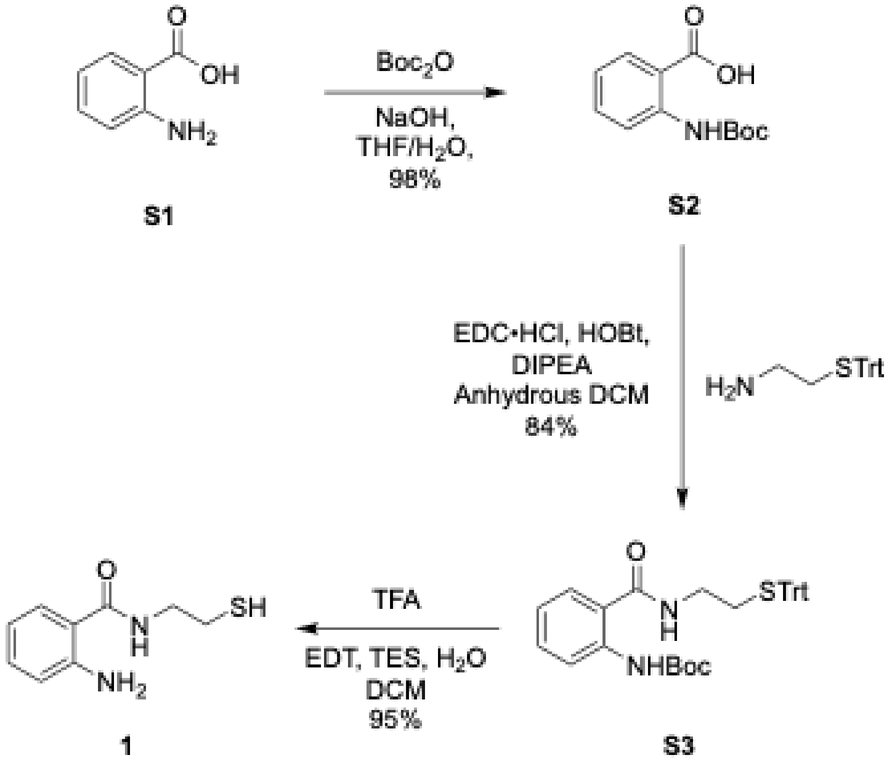

Scheme S1: Synthesis of AbzSH (1)

| **Table S1: Optimisation of thiol–ene reaction conditions** | | | | | | | | |
|---|---|---|---|---|---|---|---|---|
| ***Entry*** | ***Alkene 2 (equiv.)*** | ***Thiol 1 (equiv.)*** | ***Photoinitiator*** | ***Light Source Wavelength (nm)*** | ***Reaction Time (min)*** | ***Solvent*** | ***Additives*** | ***Yield (%)*** |
| 1 | 1.0 | 1.0 | DPAP | 365 | 30 | DCM | - | trace |
| 2 | 1.0 | 1.2 | DPAP | 365 | 30 | DCM | - | trace |
| 3 | 1.0 | 1.5 | DPAP | 365 | 30 | DCM | - | trace |
| 4 | 1.0 | 2.0 | DPAP | 365 | 30 | DCM | - | trace |
| 5 | 1.0 | 3.0 | DPAP | 365 | 30 | DCM | - | trace |
| 6 | 1.0 | 5.0 | DPAP | 365 | 30 | DCM | - | trace |
| 7 | 1.0 | 1.2 | DPAP | 254 | 30 | DCM | - | trace |
| 8 | 1.0 | 1.2 | Irgacure 2959 | 365 | 30 | DCM | - | trace |
| 9 | 1.0 | 1.2 | Eosin Y | 525 | 30 | DCM | - | trace |
| 10 | 1.0 | 1.0 | Cumene Hydroperoxide | [a] | 180 | Toluene | - | trace |
| 11 | 1.0 | 5.0 | Cumene Hydroperoxide | [a] | 180 | Toluene | - | trace |
| 12 | 1.0 | 1.0 | DPAP | Dark[b] | 60 | DCM | - | No reaction |
| 13 | 1.0 | 1.2 | DPAP | 365 | 5 | DCM | - | trace |
| 14 | 1.0 | 1.2 | DPAP | 365 | 15 | DCM | - | trace |
| 15 | 1.0 | 1.2 | DPAP | 365 | 30 | DCM | - | trace |
| 16 | 1.0 | 1.2 | DPAP | 365 | 60 | DCM | - | trace |
| 17 | 1.0 | 1.2 | DPAP | 365 | 90 | DCM | - | trace |
| 18 | 1.0 | 1.2 | DPAP | 365 | 120 | DCM | - | trace |
| 19 | 1.0 | 1.2 | DPAP | 365 | 30 | DCM | - | trace |
| 20 | 1.0 | 1.2 | DPAP | 365 | 30 | EtOAc | - | trace |
| 21 | 1.0 | 1.2 | DPAP | 365 | 30 | Hexane | - | trace |
| 22 | 1.0 | 1.2 | DPAP | 365 | 30 | ACN | - | trace |
| 23 | 1.0 | 1.2 | DPAP | 365 | 30 | DMSO | - | trace |
| 24 | 1.0 | 1.2 | DPAP | 365 | 30 | DMF | - | trace |
| 25 | 1.0 | 1.2 | DPAP | 365 | 30 | $H_2O$/MeOH | - | 15 |
| 26 | 1.0 | 1.2 | DPAP | 365 | 30 | $H_2O$/MeOH | Gdn.HCl | 22 |
| **27** | **1.0** | **1.2** | **DPAP** | **365** | **30** | **HEPES Buffered $H_2O$/MeOH** | **-** | **80** |

[a] No irradiation. Reaction was carried out under thermal conditions (reflux at 80 °C).

[b] Reaction stirred in the absence of light as a negative control for photochemical radical initiation.

### 2. General Experimental Information

All commercially available chemicals were purchased from Merck KGaA, Fluorochem Ltd or Tokyo Chemical Industry (TCI) and used without further purification unless otherwise stated. Solvents for synthesis and purification were supplied by Sigma-Aldrich at HPLC grade. Room temperature (rt) was ~20 °C. All UV reactions were carried out in a Luzchem photoreactor, LZC-EDU (110 V/60 Hz), containing 14 UV-A lamps centred at 352 nm. All blue LED reactions were carried out using a Kessil PR160 Standard mounting system, KSPRRM03 fitted with 2 Kessil LED PhotoReaction Lights (KSPR160L-440) centred at 440 nm. Green LED reactions were carried out using the same set-up but fitted with 1 Kessil LED PhotoReaction Light (KSPR160L-525) centred at 525 nm.

**Nuclear Magnetic Resonance (NMR) Spectroscopy**

NMR spectra were recorded using Bruker 400HD (400.13 MHz for $^{1}$H NMR and 100.61 MHz for $^{13}$C NMR), Bruker AV 400 (400.13 MHz for $^{1}$H NMR and 100.61 MHz for $^{13}$C NMR), Agilent MR400 (400.13 MHz for $^{1}$H NMR and 100.61 MHz for $^{13}$C NMR) or Bruker AV 600 (600.13 MHz for $^{1}$H NMR and 150.90 MHz for $^{13}$C NMR) instruments. Deuterated solvents for NMR were purchased from Sigma Aldrich (Merck). Chemical shifts, δ, are in ppm and referenced to the internal solvent signals for $CDCl_3$ (δH 7.26 and δC 77.16) or DMSO (δH 2.50 and δC 39.52). NMR descriptions: s, singlet; d, doublet; t, triplet; m, multiplet; br., broad; quint. quintuplet; sext. Sextuplet; app. apparent. Coupling constants, *J*, are reported in Hertz (Hz). NMR data were processed using TopSpin software (version 4.0.9) by Bruker BioSpin.

**High Resolution Mass Spectrometry (HRMS)**

Electrospray ionisation (ESI) mass spectra were acquired using a Bruker micrOTOF-Q III spectrometer interfaced to a Dionex UltiMate 3000 LC in positive and negative modes as required. Agilent ESI-L Low Concentration Tuning Mix was used to calibrate the system, and this was also used as an internal lock mass. Masses were recorded over the range 100-2000 m/z. Operating conditions were as follows: end-plate offset 500 V, capillary 4500 V, nebuliser 2.0 bar, dry gas 8.0 L/min, and dry temperature 180 °C. Compass HyStar version 3.2 and micrOTOF Control version 3.2 software were used to analyse the data.

### 3. General Procedures

#### General procedure A: AbzSH labelling

AbzSH **1** (1 eq.), DPAP (0.25 eq.) and alkene containing substrate were dissolved in HEPES buffer solution* forming a 0.1 M solution of alkene substrate. MeOH (<30% v/v) was added as necessary to dissolve residual organics and the reaction mixture sonicated for 5 min to ensure homogeneity. The reaction mixture was irradiated under UV-A irradiation for 1 h with vigorous stirring. The solvent was evaporated and the mixture re-suspended in DCM : $H_2O$ (1:1). The organic layer was separated, dried over $MgSO_4$, filtered and solvent evaporated and purified by flash column chromatography.

*HEPES buffer: HEPES (1.19 g) and NaCl (0.88 g) dissolved in deionised water (80 mL). pH was adjusted to pH 7.4 utilising 0.2 M NaOH as necessary and the solution made up to a total volume of 100 mL with deionised water.

### 4. Fluorophore Synthesis

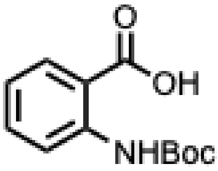

**2-((*tert*-butoxycarbonyl)amino)benzoic acid[1] (S2)**

To a suspension of anthranilic acid (5.00 g, 36.46 mmol) in THF/$H_2O$ (1:1, 80 mL), NaOH was added until pH 8 was achieved. $Boc_2O$ (8.76 g, 40.10 mmol) was then added and the reaction stirred overnight. The THF was then removed *in vacuo* and the mixture acidified with citric acid (10% w/v) to give a white precipitate. The mixture was then extracted with DCM (3 x 150 mL) and the combined organics washed with citric acid (10% w/v, 3 x 150 mL), dried over $MgSO_4$, filtered, and concentrated *in vacuo* to yield the product as an off-white solid (8.48 g, 98%).

**$R_f$** = 0.07 (15% EtOAc/Hexane (v/v)).

**$^1$H NMR (400 MHz, DMSO-d6)** $\delta_H$ 10.51 (br. s, 1H, NH), 8.47 (dd, *J* = 8.5, 1.0 Hz, 1H, ArH), 8.08 (dd, *J* = 8.0, 1.6 Hz, 1H, ArH), 7.56 (td, *J* = 8.0, 1.6 Hz, 1H, ArH), 7.03 (td, *J* = 8.0, 1.1 Hz, 1H, ArH), 1.54 (s, 9H, Boc 3 x $CH_3$) ppm.

***m/z* HRMS (ESI$^+$)** calcd. for $C_{12}H_{15}NO_4Na$ ([M+Na]$^+$): 260.0893; found: 260.0900.

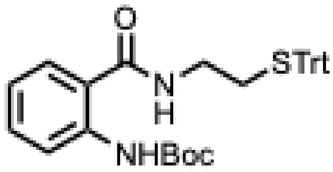

**tert-butyl (2-((2-(tritylthio)ethyl)carbamoyl)phenyl)carbamate (S3)**

Under $N_2$, 2-((*tert*-butoxycarbonyl)amino)benzoic acid **S2** (1.00 g, 4.21 mmol), EDC•HCl (885 mg, 4.63 mmol), HOBt (625 mg, 4.63 mmol) and DIPEA (1.47 mL, 8.42 mmol) were dissolved in dry DCM (25 mL) at 0 °C. A solution of 2-tritylthio-1-ethylamine hydrochloride (1.48 g, 4.64 mmol) in dry DCM (15 mL) was added dropwise to this stirred suspension, which was then allowed to warm to rt and stirred for 15 h. The reaction mixture was diluted with DCM (20 mL) and washed with 0.1 M aq. HCl solution (20 mL), $H_2O$ (20 mL), sat. aq. $NaHCO_3$ solution (20 mL), brine (20 mL), dried over $Na_2SO_4$, filtered and concentrated *in vacuo* to afford an off-white foam. The product was then purified by silica gel flash chromatography using DCM to afford the desired product as a white solid (1.96 g, 3.64 mmol, 86%).

$\mathbf{R_f}$ = 0.48 (20% EtOAc/Hexane (v/v)).

**$^1$H NMR (400 MHz, $CDCl_3$)** $\delta_H$ 10.07 (br. s, 1H, NHBoc), 8.36 (d, *J* = 8.3 Hz, 1H, ArH), 7.47-7.16 (m, 17H, 2 x ArH, 15 x Trt ArH), 6.97 (t, *J* = 7.5 Hz, 1H, ArH), 6.25 (br. s, 1H, CONH), 3.26 (dt, *J* = 6.3, 6.1 Hz, 2H, CONH$\underline{CH_2}$), 2.53 (t, *J* = 6.1 Hz, 2H, $\underline{CH_2}$S), 1.51 (s, 9H, Boc 3 x $CH_3$) ppm.

**$^{13}$C NMR (151 MHz, $CDCl_3$)** $\delta_C$ 168.7 (CONH), 153.1 (Boc C=O), 144.5 (Trt ArC), 140.4 (ArC), 132.6 (ArH), 129.5 (Trt ArH), 128.0 (Trt ArH), 126.9 (Trt ArH), 126.6 (ArC), 121.3 (ArH), 119.8 (ArH), 119.5 (ArH), 80.2 (Trt ArH), 67.0 (Trt ArC), 38.3 (CONH$CH_2$), 32.0 ($CH_2$S), 28.4 (Boc $CH_3$) ppm.

***m/z* HRMS (ESI$^+$)** calcd. for $C_{33}H_{34}N_2O_3SNa$ ([M+Na]$^+$): 561.2182; found: 561.2195.

$\mathbf{\nu_{max}}$ (film), cm$^{-1}$: 3309 (N-H stretch), 1696, 1641 (C=O stretch).

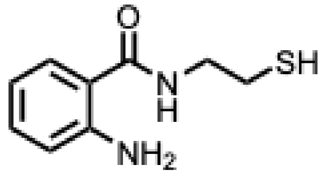

**2-amino-*N*-(2-mercaptoethyl)benzamide[2] (1)**

Tert-butyl (2-((2-(tritylthio)ethyl)carbamoyl)phenyl)carbamate **S3** (2.00 g, 3.71 mmol) was dissolved in a DCM/TFA/$H_2O$/TES/EDT cocktail (44:50:2.5:1.0:2.5, 18.6 mL). The mixture was stirred for 1 h, following which the TFA was removed under $N_2$ flow. The product was then purified by silica gel flash chromatography using 25-40% EtOAc/Hexane (v/v) to yield the desired product as a yellow wax (716 mg, 3.65 mmol, 98%).

$\mathbf{R_f}$ = 0.40 (50% EtOAc/Hexane (v/v)).

**$^1$H NMR (400 MHz, CDCl$_3$)** $\delta_H$ 7.40 (dd, *J* = 7.8, 1.3 Hz, 1H ArH), 7.27 (ddd, 1H, *J* = 8.4, 7.0, 1.3 Hz, ArH), 6.81-6.73 (m, 2H, 2 x ArH), 6.66 (br. s, 1H, CONH), 6.23 (br. s, 2H, NH$_2$), 3.61 (dt, *J* = 6.3, 6.2 Hz, 2H, CONH<u>CH$_2$</u>), 2.79 (dt, *J* = 8.3, 6.3 Hz 2H, <u>CH$_2$</u>SH), 1.44 (t, *J* = 8.4 Hz, 1H, SH) ppm.

***m/z* HRMS (ESI$^+$)** calcd. for $C_9H_{13}N_2OS$ ([M+H]$^+$): 197.0743; found: 197.0753.

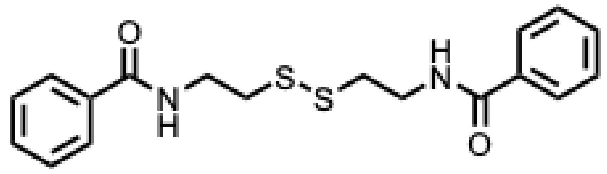

***N,N*'-(disulfanediylbis(ethane-2,1-diyl))dibenzamide[3] (S4)**

Cysteamine dichloride (249 mg, 1.10 mmol) and $Et_3N$ (0.7 mL, 5.00 mmol) were dissolved in anhydrous DCM (20 mL) and cooled to 0 °C. Benzoyl chloride (0.26 mL, 2.21 mmol) was slowly added and the reaction mixture was allowed to warm to rt. After 16 h, the reaction mixture was diluted with DCM (2 × 20 mL). The combined organic layers were washed with $H_2O$ (20 mL), sat. aq. $NaHCO_3$ (20 mL) and brine (20 mL). The product was then purified by silica gel flash chromatography using 50% Hex/EtOAc (v/v) to yield the desired product as a white solid (335 mg, 0.93 mmol, 84%).

**R$_f$** = 0.22 (50% Hex/EtOAc (v/v)).

**$^1$H NMR (400 MHz, CDCl$_3$)** $\delta_H$ 7.82 (d, *J* = 7.7 Hz 4H, ArH), 7.49 (t, *J* = 7.4 Hz, 2H, ArH), 7.41 (t, *J* = 7.4 Hz, 4H, ArH), 7.05 (br. s, 2H, CONH), 3.79 (app. q, *J* = 6.2, 6.1 Hz, 4H, NH<u>CH$_2$</u>), 2.98 (t, *J* = 6.1 Hz, 4 H, <u>CH$_2$</u>S) ppm.

***m/z* HRMS (ESI$^+$)** calcd. for 383.0864 ([M+Na]$^+$): ; found: 383.0858.

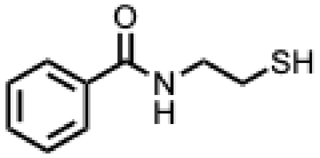

***N*-(2-mercaptoethyl)benzamide[4] (3)**

TCEP•HCl (133 mg, 0.47 mmol) was dissolved in $H_2O$ (3.6 mL), degassed and adjusted to pH 7 with sat. aq. $NaHCO_3$. This solution was added to a solution of *N,N*'-(disulfanediylbis(ethane-2,1-diyl))dibenzamide **S4** (129 mg, 0.36 mmol) in MeOH (18 mL) and stirred at rt for 1 h. MeOH was then evaporated under reduced pressure. The aqueous residue was diluted with DCM (2 x 25 mL), washed with $H_2O$ (50 mL) and brine (50 mL). The organic phase was dried over $MgSO_4$, filtered and concentrated *in vacuo*. The product was isolated as a white solid (121.2 mg, 0.67 mmol, 93%).

**$R_f$** = 0.86 (EtOAc).

**$^{1}$H NMR (400 MHz, $CDCl_3$)** $\delta_H$ 7.79-7.75 (m, 2H, ArH), 7.46 (tt, *J* = 7.3, 1.3 Hz, 1H, ArH), 7.40-7.35 (m, 2H, ArH), 7.09 (br. s, 1H, CONH), 3.57 (dt, *J* = 6.6, 6.2 Hz, 2H, NH$\underline{CH_2}$), 2.72 (dt, *J* = 8.5, 6.6 Hz, 2H, $\underline{CH_2}$SH), 1.39 (t, *J* = 8.5 Hz, 1H, $CH_2\underline{SH}$) ppm.

***m/z*** **HRMS ($APCI^-$)** calcd. for 180.0478 ($[M-H]^-$): ; found: 180.0481.

### 5. Synthesis of Alkenes

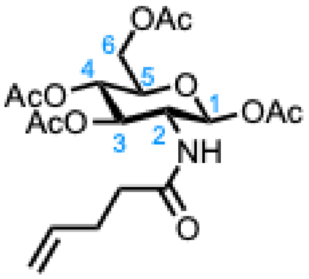

**(2*S*,3*R*,4*R*,5*S*,6*R*)-6-(acetoxymethyl)-3-(pent-4-enamido)tetrahydro-2*H*-pyran-2,4,5-triyl triacetate[5] (2)**

Under $N_2$, 4-pentenoic acid (0.20 mL, 196 mg, 1.66 mmol), EDC•HCl (223 mg, 1.44 mmol), HOBt (194 mg, 1.44 mmol) and DIPEA (2.9 mL, 16.7 mmol) were dissolved in dry DCM (6 mL) at 0 °C. A solution of (2S,3R,4R,5S,6R)-6-(acetoxymethyl)-3-amino-tetrahydro-2H-pyran-2,4,5-triyl triacetate•HCl (500 mg, 1.31 mmol) in dry DCM (8 mL) was added dropwise to this stirred suspension, which was then allowed to warm to rt and stirred for 6 h. The reaction mixture was diluted with EtOAc (20 mL) and washed with $H_2O$ (20 mL), sat. aq. $NaHCO_3$ solution (20 mL), brine (20 mL), dried over $Na_2SO_4$, filtered and concentrated *in vacuo* to afford a white foam. The crude product was purified by flash column chromatography using 25-50% Hex/EtOAc (v/v) to yield the desired product as a white foam (498 mg, 1.16 mmol, 89%).

**$R_f$** = 0.22 (50% Hex/EtOAc (v/v)).

**$^1$H NMR (400 MHz, $CDCl_3$)** $\delta_H$ 5.86 (d, *J* = 9.7 Hz, 1H, CONH), 5.74 (m, 1H, $CH_2\underline{CH}$) 5.68 (d, *J* = 8.9 Hz, 1H, $C_1H$), 5.16 (dd, *J* = 12.0, *J* = 9.7, 1H, $C_4H$), 5.10 (app. t, *J* = 9.7, 1H, $C_3H$), 5.01 (app. dq, 17.1, 1.6 Hz, 1H, $\underline{CH_2}CH$ x1) 4.96 (app. dq, *J* = 10.3, 1.6 Hz, 1H, $\underline{CH_2}CH$ x1), 4.31 (dd, *J* = 19.2, *J* = 9.7, 1H, $C_2H$), 4.25 (dd, *J* = 12.5, 4.8 Hz, 1H, $C_6H'$), 4.11 (dd, *J* = 12.5, 2.3 Hz, 1H, $C_6H$), 3.82 (ddd, *J* = 9.8 Hz, *J* = 4.8 Hz, *J* = 2.3 Hz, 1H, $C_5H$), 2.34-2.27 (m, 2H, $COCH_2\underline{CH_2}$), 2.23-2.17 (m, 2H, $COCH_2$), 2.09 (s, 3H, 3 x $C_1$-Acet $\underline{C}H_3$), 2.07 (s, 3H, 3 x $C_6$-Acet $\underline{C}H_3$), 2.03 (s, 3H, 3 x $C_3$-Acet $\underline{C}H_3$), 2.02 (s, 3H, 3 x $C_4$-Acet $\underline{C}H_3$) ppm.

***m/z* HRMS (ESI$^+$)** calcd. for $C_{19}H_{27}NNaO_{10}$ ($[M+Na]^+$): 452.1527 ; found: 452.1518.

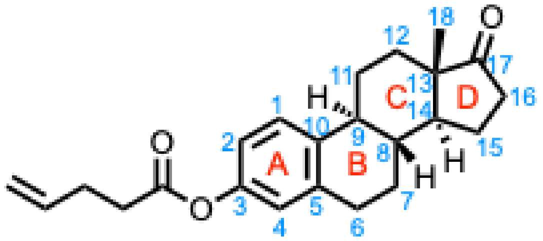

**(8*R*,9*S*,13*S*,14*S*)-13-methyl-17-oxo-7,8,9,11,12,13,14,15,16,17-decahydro-6*H*-cyclopenta[*a*]phenanthren-3-yl pent-4-enoate (7)**

To a solution of 4-pentenoic acid (1.9 mmol, 0.190 g, 0.194 mL) in dry DCM (12 mL) at 0 °C, EDC•HCl (1.9 mmol, 0.364 g) was added and stirred for 15 min. DMAP (0.4 mmol, 0.049 g) and estrone (1.5 mmol 0.406 g) was then added. The solution was warmed to rt and stirred for 24 h. The solvent was then removed *in vacuo*. The product was purified by silica gel flash chromatography using 25% EtOAc/Hexane (v/v) to yield the desired product as a colourless oil (333 mg, 66%). The isolated product was in agreement with the literature literature.[6]

**R$_f$** = 0.76 (50% Hex/EtOAc (v/v)).

**$^1$H NMR (400 MHz, CDCl$_3$)** $\delta_H$ 7.31 (d, *J* = 8.5 Hz, 1H, A-1), 6.89-6.81 (m, A-1, A-4), 5.98-5.87 (m, 1H, CH$_2$<u>CH</u>), 5.17 (qd, *J* = 17.1, 1.4 Hz, 1H, <u>CH$_2$</u>CH x1), 5.10 (qd, *J* = 10.2, 1.4 Hz, 1H, <u>CH$_2$</u>CH x1), 2.97-2.90 (m, 2H, B-2), 2.68 (t, *J* = 7.4 Hz, COCH$_2$), 2.58-2.49 (m, 2H, COCH$_2$<u>CH$_2$</u>), 2.47-2.40 (m, 1H, D-16 x1), 2.37-2.27 (m, 1H, C-11 x1), 2.23-2.10 (m, 2H, B/C-9, D-16 x1), 2.10-1.95 (m, 2H, B-7 x1, D-15 x1), 1.72-1.41 (m, 7H, B-7 x1, B/C-8, C/D-14, D-15 x1, C-12, C-11 x1), 0.93 (s, 3H, C/D-18) ppm.

***m/z* HRMS (APCI$^+$)** calcd. for $C_{23}H_{29}O_3$ ([M+H]$^+$): 353.2111; found: 353.2111.

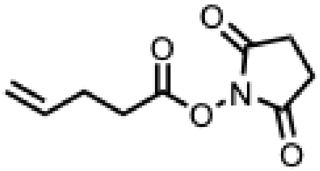

**2,5-dioxopyrrolidin-1-yl pent-4-enoate[7] (S5)**

*N*-hydroxysuccinimide (0.096 g, 0.83 mmol) and DMAP (0.010, 0.08 mmol) were dissolved in dry DCM and DCC (0.137 mL, 0.171 g, 0.83 mmol) was added, followed by pent-4-enoic acid (0.077 mL, 0.075 g, 0.75 mmol). The mix was stirred under Ar for 24 h, filtered, and concentrated *in vacuo*. The product was then purified by silica gel flash chromatography using 0.2-0.4% MeOH/DCM (v/v) to yield the desired product as a white solid (0.1107 g, 75%).

**R$_f$** = 0.41 (0.2% MeOH/DCM (v/v)).

**$^1$H NMR (400 MHz, CDCl$_3$)** $\delta_H$ 5.92-5.80 (m, 1H, CH$_2$<u>CH</u>), 5.16-5.07 (m, 2H, <u>CH$_2$</u>CH), 2.84 (br. s, 4H, 2 x *N*HS CH$_2$), 2.72 (t, *J* = 7.3 Hz, 2H, COCH$_2$), 2.50 (q, *J* = 7.3 Hz, 2H, COCH$_2$<u>CH$_2$</u>) ppm.

***m/z* HRMS (ESI$^+$)** calcd. for $C_9H_{11}NO_4Na$ ([M+Na]$^+$): 220.0580; found: 220.0580.

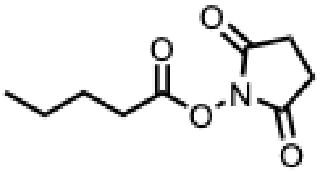

**2,5-dioxopyrrolidin-1-yl pentanoate**[8] **(S6)**

*N*-hydroxysuccinimide (0.096 g, 0.83 mmol) and DMAP (0.010, 0.08 mmol) were dissolved in dry DCM and DCC (0.137 mL, 0.171 g, 0.83 mmol) was added, followed by pentanoic acid (0.081 mL, 0.077 g, 0.75 mmol). The mix was stirred under Ar for 24 h, filtered, and concentrated *in vacuo*. The product was then purified by silica gel flash chromatography using 0.4% MeOH/DCM (v/v) to yield the desired product as a colourless oil (0.0.1307 g, 87%).

**$R_f$** = 0.45 (0.2% MeOH/DCM (v/v)).

**$^1$H NMR (400 MHz, $CDCl_3$)** $\delta_H$ 2.83 (br. s, 4H, 2 x *N*HS $CH_2$), 2.61 (t, *J* = 7.5 Hz, 2H $COCH_2$), 1.73 (quint., *J* = 7.5 Hz, 2H, $COCH_2\underline{CH_2}$), 1.44 (sext., *J* = 7.5 Hz, 2H, $CH_3\underline{CH_2}$), 0.95 (t, *J* = 7.5 Hz, 3H, $CH_3$) ppm.

***m/z* HRMS (ESI$^+$)** calcd. for $C_9H_{13}NO_4Na$ ($[M+Na]^+$): 222.0737; found: 222.0736.

### 6. Synthesis of Labelled Products

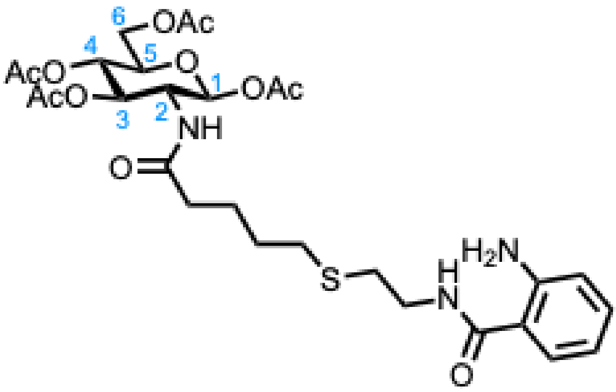

**(2*S*,3*R*,4*R*,5*S*,6*R*)-6-(acetoxymethyl)-3-(5-((2-(2-aminobenzamido)ethyl)thio)pentanamido) tetrahydro-2*H*-pyran-2,4,5-triyl triacetate (4)**

#### HEPES buffer procedure

*N*-(2-mercaptoethyl)benzamide **1** (35 mg, 0.19 mmol, 1.2 equiv.), (2S,3R,4R,5S,6R)-6-(acetoxymethyl)-3-(pent-4-enamido)tetrahydro-2*H*-pyran-2,4,5-triyl triacetate **2** (70 mg, 0.16 mmol, 1.0 equiv.) and DPAP (10.4 mg, 41 µmol) were dissolved in 50 mM HEPES aq. buffer solution containing 150 mM NaCl (12 mL). MeOH (3 mL) was added. The resulting mixture was sonicated for 5 min and then irradiated at 365 nm at rt with stirring for 30 min. MeOH was evaporated under reduced pressure. The aqueous mixture was diluted with DCM (2 x 10 mL). The combined organics were washed with $H_2O$ (2 x 20 mL) and brine (2 x 20 mL), dried over $Na_2SO_4$, filtered and concentrated *in vacuo*. The crude product was purified by flash column chromatography (EtOAc) to yield the desired product as a colourless oil (82 mg, 0.13 mmol, 80%).

#### Gdn•HCl procedure

AbzSH **1** (10 mg, 0.05 mmol, 1 eq.), sugar **2** (21 mg, 0.005 mg, 1 eq.) and DPAP (2.6 mg, 0.01 mmol, 0.2 eq.) were dissolved in $H_2O$ : MeOH 1:1 (5 mL) and Gdn•HCl (0.29 g, 6M) was added. Reaction mixture was sonicated until fully dissolved and stirred under UV-A irradiation for 1 h. Solvent evaporated and reaction mixture partitioned between $H_2O$ : DCM. Organic layer separated, dried and purified by flash column chromatography (8:2 EtOAc:Hexane) to yield the product as a pale yellow oil (7 mg, 22%)

**$R_f$** = 0.76 (EtOAc).

**$^1$H NMR (400 MHz, $CDCl_3$)** $\delta_H$ 7.46 (d, *J* = 7.6 Hz, 1H, ArH), 7.27 (t, *J* =7.5 Hz, 2H, 1 x ArH), 6.85 (d, *J* = 7.9 Hz, ArH) 6.82-6.76 (m, 2H, 1 x ArH, 1 x CONH) 6.34 (d, *J* = 9.5 Hz, 1H, $CONHC_2$), 5.51 (d, *J* = 8.8 Hz, 1H, $C_1H$), 5.06-5.00 (m, 2H, 1 x $C_3H$, 1 x $C_4H$), 4.32-4.26 (m, 1H, $C_2H$), 4.19 (d, *J* = 12.6, 4.5 Hz, 1H, $C_6H$), 4.03 (dd, *J* = 12.6, 2.1 Hz, 1H, $C_6H'$), 3.64-3.53 (m, 2H, CONH$\underline{CH_2}$), 3.44 (ddd, *J* = 9.9, 4.5, 2.1 Hz, 1H, $C_5H$), 2.76-2.70 (m, 2H,$CH_2S$), 2.54 (t, *J* = 7.3 Hz, 2H, $SCH_2$), 2.16 (t, *J* = 7.3 Hz, 2H, $CH_2CO$), 2.08 (s, 3H, 3 x $C_1$-OAcet $CH_3$), 2.07 (s,

3H, 3 x $C_6$-OAcet $CH_3$), 2.01 (s, 6H, 3 x $C_3$-OAcet $CH_3$, 3 x $C_4$-OAcet $CH_3$), 1.74-1.64 (m, 2H, $SCH_2CH_2\underline{CH_2}$), 1.59-1.51 (m, 2H, $SCH_2\underline{CH_2}$) ppm.

**$^{13}$C NMR (151 MHz, CDCl$_3$)** $\delta_C$ 173.1 ($\underline{C}ONHC_2$), 171.1 ($C_3$-Acet $\underline{C}$=O), 170.7 ($C_6$-Acet $\underline{C}$=O), 169.7 ($C_1$-Acet $\underline{C}$=O), 169.6 ($C_4$-Acet $\underline{C}$=O), 169.3 (CONH), 146.3 ($ArNH_2$), 132.5 (ArH), 127.9 (ArH), 118.4 (ArH), 118.3 (ArH), 118.0 (ArC), 92.5 ($C_1$), 72.8 ($C_3$), 72.5 ($C_5$), 67.9 ($C_4$), 61.5 ($C_6$), 52.4 ($C_2$), 39.1 ($CONH\underline{C}H_2$), 35.5 ($CH_2CO$), 31.4 ($\underline{C}H_2S$), 30.7 ($SCH_2$), 28.8 ($SCH_2\underline{C}H_2$), 24.5 ($SCH_2CH_2\underline{C}H_2$), 21.0 ($C_1$-Acet $\underline{C}H_3$), 20.8 ($C_6$-Acet $\underline{C}H_3$), 20.6 ($C_3$-Acet $\underline{C}H_3$), 20.6 ($C_4$-Acet $\underline{C}H_3$) ppm.

***m/z* HRMS (ESI$^+$)** calcd. for $C_{28}H_{40}N_3O_{11}S$ ($[M+H]^+$): 626.2378; found: 626.2366.

**$\nu_{max}$** (film), cm$^{-1}$: 3301 (N–H stretch), 1752 (C=O stretch), 1221 (C–O stretch), 1074 (C–O stretch).

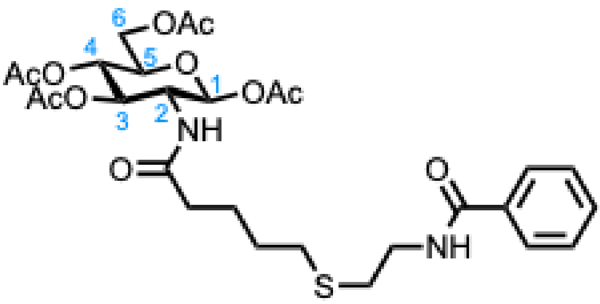

**(2*S*,3*R*,4*R*,5*S*,6*R*)-6-(acetoxymethyl)-3-(5-((2-benzamidoethyl)thio)pentanamido)tetrahydro-2*H*-pyran-2,4,5-triyl triacetate (5)**

*N*-(2-mercaptoethyl)benzamide **1** (35 mg, 0.20 mmol, 1.2 equiv.), (2*S*,3*R*,4*R*,5*S*,6*R*)-6-(acetoxymethyl)-3-(pent-4-enamido)tetrahydro-2*H*-pyran-2,4,5-triyl triacetate **2** (70 mg, 0.16 mmol, 1.0 equiv.) and DPAP (10.4 mg, 41 µmol) were dissolved in 50 mM HEPES aq. buffer solution containing 150 mM NaCl (12 mL). MeOH (3 mL) was added. The resulting mixture was sonicated for 5 min and then irradiated at 365 nm at rt with stirring for 30 min. MeOH was evaporated under reduced pressure. The aqueous mixture was diluted with DCM (2 x 10 mL). The combined organics were washed with $H_2O$ (2 x 20 mL) and brine (2 x 20 mL), dried over $Na_2SO_4$, filtered and concentrated *in vacuo*. The crude product was purified by flash column chromatography (EtOAc) to yield the desired product as a colourless oil (78 mg, 0.12 mmol, 76%).

**$R_f$** = 0.63 (EtOAc).

**$^1$H NMR (400 MHz, CDCl$_3$)** $\delta_H$ 7.84-7.81 (m, 2H, ArH), 7.55-7.50 (m, 1H, ArH), 7.55-7.44 (m, 2H, ArH), 6.72 (t, *J* = 5.2 Hz, 1H, CONH), 6.28 (d, *J* =9.5 Hz, 1H, $CONHC_2$), 5.59 (d, *J* = 8.9 Hz, 1H, $C_1H$), 5.12-5.04 (m, 2H, $C_{3,4}H$), 4.32-4.25 (m, 1H, $C_2H$), 4.21 (dd, *J* = 12.5, 4.5 Hz, 1H, $C_6H$), 4.05 (dd, *J* = 12.5, 2.2 Hz, 1H, $C_6H'$), 3.67-3.60, (m, 2H, $CONH\underline{CH_2}$), 3.53-3.49 (m, 1H, $C_5H$), 2.74 (t, *J* = 6.5 Hz, 2H, $CH_2S$), 2.55 (t, *J* = 6.9 Hz, 2H, $SCH_2$), 2.16 (t, *J* = 7.1 Hz, 2H, $CH_2CO$), 2.10 (s, 3H, 3 x $C_1$-Acet $\underline{C}H_3$), 2.08 (s, 3H, 3 x $C_6$-Acet $\underline{C}H_3$), 2.02 (s, 3H, 3 x $C_3$-Acet $\underline{C}H_3$), 2.02 (s, 3H, 3 x $C_4$-Acet $\underline{C}H_3$), 1.75-1.65 (m, 2H, $SCH_2CH_2\underline{CH_2}$), 1.62-1.54 (m, 2H, $SCH_2\underline{CH_2}$) ppm.

**$^{13}$C NMR (151 MHz, $CDCl_3$)** $\delta_C$ 173.2 ($CONHC_2$), 171.1 ($C_3$-Acet C=O), 170.8 ($C_6$-Acet C=O), 169.7 ($C_1$-Acet C=O), 169.4 ($C_4$-Acet C=O), 168.2 (CONH), 134.5 (ArC), 131.9 (ArH), 128.9 (ArH), 127.2 (ArH), 92.6 ($C_1$), 72.8 ($C_5$), 72.7 ($C_3$), 67.9 ($C_4$), 61.7 ($C_6$), 52.7 ($C_2$), 39.4 ($CONHCH_2$), 35.7 ($CH_2CO$), 31.5 ($CH_2S$), 30.8 ($SCH_2$), 28.8 ($SCH_2CH_2$), 24.5 ($SCH_2CH_2CH_2$), 21.1 ($C_1$-Acet $CH_3$), 20.8 ($C_6$-Acet $CH_3$), 20.8 ($C_3$-Acet $CH_3$), 20.7 ($C_4$-Acet $CH_3$) ppm.

***m/z* HRMS (ESI$^+$)** calcd. for $C_{28}H_{39}N_2O_{11}S$ ($[M+H]^+$): 611.2269; found: 611.2259.

**$\nu_{max}$** (film), $cm^{-1}$: 3300 (N–H stretch), 2935 (C–H stretch), 1751 (C=O stretch), 1220 (C–O stretch).

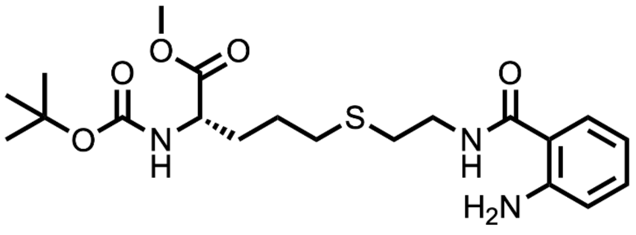

**methyl 5-((2-(2-aminobenzamido)ethyl)thio)-2-((*tert*-butoxycarbonyl)amino)pentanoate (S7)**

From Boc-Agl-OMe **6**, compound **S7** was synthesised on a 0.3 mmol scale as per general procedure A. The product was purified by flash column chromatography (0-2.5% MeOH in DCM) and obtained as a pale yellow oil (37 mg, 30%).

**$R_f$** = 0.31 (0.5% MeOH/DCM (v/v)).

**$^1$H NMR (400 MHz, DMSO-d6)** $\delta_H$ 8.31 (t, *J* = 5.5 Hz, 1H, Ar-CONH), 7.46 (d, *J* = 8.0 Hz, 1H, ArH), 7.28 (d, *J* = 8.0 Hz, 1H, OCONH), 7.13 (t, *J* = 7.5 Hz, 1H, ArH), 6.69 (d, *J* = 8.0 Hz, 1H, ArH), 6.51 (t, *J* = 7.5 Hz, 1H, ArH), 6.39 (s, 2H, $NH_2$), 4.03-3.94 (m, 1H, Agl $\alpha$CH), 3.63 (s, 3H, $OCH_3$), 3.41-3.35 (m, 2H, Ar-$CONHCH_2$), 2.65 (t, *J* = 7.1 Hz, 2H, Ar-$CONHCH_2CH_2$), 2.56-2.49 (m, 2H, Agl $SCH_2$), 1.82-1.72 (m, 1H, Agl $SCH_2CH_2$ x1), 1.71-1.55 (m, 3H, Agl $SCH_2CH_2$ x1, $SCH_2CH_2CH_2$), 1.39 (s, 9H, Boc 3 x $CH_3$) ppm.

**$^{13}$C NMR (100.5 MHz, DMSO-d6)** $\delta_C$ 173.5 ($COOCH_3$), 169.3 (Ar-CO), 156.0 (ArC), 150.1 (ArC), 132.1 (ArCH), 128.4 (ArCH), 116.8 (ArCH), 115.0 (ArCH), 78.7 (Boc CH), 53.6 (Agl $\alpha$CH), 52.2 ($OCH_3$), 39.3 ($CH_2NH$), 30.8 ($SCH_2$), 30.3 (Agl $CH_2$), 28.6 (Boc $CH_3$), 26.0 (Agl $CH_2$) ppm.

***m/z* HRMS (ESI$^+$)** calcd. for $C_{20}H_{31}N_3O_5SNa$ ($[M+Na]^+$): 448.1877; found: 448.1865.

**$\nu_{max}$** (film), $cm^{-1}$: 3408 (N-H stretch), 2968 (C-H stretch), 1667 (C=O stretch).

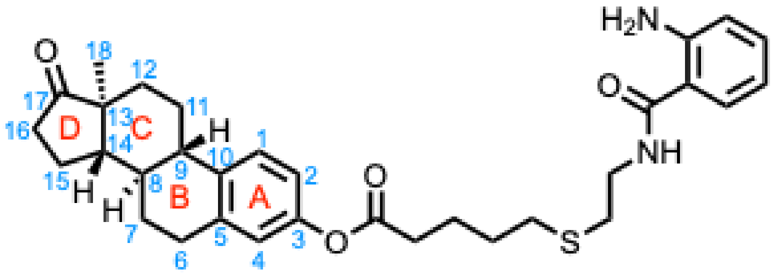

**(8*R*,9*S*,13*S*,14*S*)-13-methyl-17-oxo-7,8,9,11,12,13,14,15,16,17-decahydro-6*H*-cyclopenta[*a*]phenanthren-3-yl 5-((2-(2-aminobenzamido)ethyl)thio)pentanoate (S8)**

From (8*R*,9*S*,13*S*,14*S*)-13-methyl-17-oxo-7,8,9,11,12,13,14,15,16,17-decahydro-6*H*-cyclopenta-[*a*]-phenanthren-3-yl pent-4-enoate **7**, compound **S8** was synthesised on a 0.15 mmol scale as per general procedure A. The product was purified by flash column chromatography (0-50% v/v EtOAc in Hexane) and obtained as a colourless oil (8 mg, 10%).

**$R_f$** = 0.33 (50% Hex/EtOAc (v/v)).

**$^1$H NMR (400 MHz, DMSO-d6)** $\delta_H$ 8.34 (t, *J* = 5.6 Hz, 1H, NH), 7.46 (dd, *J* = 8.1, 1.3 Hz, 1H, Abz ArH), 7.29 (d, *J* = 8.4 Hz, 1H, A-1), 7.14 (td, *J* = 7.7, 1.3 Hz, 1H, Abz ArH), 6.85 (dd, *J* = 8.4, 2.5), 1H, A-2), 6.81 (app. d, 1.9 Hz, 1H, A-4), 6.69 (dd, *J* = 8.1, 1.0 Hz, 1H, Abz ArH), 6.51 (td, *J* = 7.7, 1.0 Hz, 1H, Abz ArH), 6.40 (br. s, 2H, $NH_2$), 3.42-3.35 (m, 2H, $NH\underline{CH_2}$), 2.87-2.81 (m, 2H, B-6), 2.66 (t, *J* = 7.4 Hz, 2H, $\underline{SCH_2}CH_2NH$), 2.59 (m, 4H, $SCH_2$, $COCH_2$), 2.45 (dd, *J* = 18.5, 8.1 Hz, 1H, D-16 x1), 2.39-2.35 (m, 1H, C-11 x1), 2.30-2.19 (m, 1H, B/C-9), 2.13-2.02 (m, 1H, D-16 x1), 2.00-1.90 (m, 2H, B-7 x1, D 15 x1), 1.80-1.69 (m, 3H, $COCH_2\underline{CH_2}$, C-12 x1), 1.68-1.59 (m, 2H, $COCH_2CH_2\underline{CH_2}$), 1.58-1.48 (m, 3H, B/C-8, C/D-14, D-15 x1), 1.43-1.32 (m, 3H, B-7 x1, C-11 x1, C-12 x1), 0.84 (s, 3H, C/D-18) ppm.

**$^{13}$C NMR (100.5 MHz, DMSO-d6)** $\delta_C$ 219.6 (C-17), 171.8 (C=OO), 168.8 (C=OAr), 149.6 (C-$NH_2$), 148.2 (A-3), 137.7 (A/B-5), 137.1 (A/B-10), 131.6 (Abz ArC), 127.9 (Abz ArC), 126.3 (A-1), 121.5 (A-4), 118.9 (A-2), 116.3 (Abz ArC), 114.5 (Abz ArC), 49.5 (C/D-14), 47.3 (C/D-13), 43.5 (B/C-9), 38.9 ($NHCH_2$), 37.4 (B/C-8), 35.3 (D-16), 33.0 ($CO\underline{CH_2}$), 31.3 (C-12), 30.4 ($SCH_2$), 30.3 ($\underline{SCH_2}CH_2NH$), 28.8 (B-6), 28.3 ($SCH_2\underline{CH_2}$), 25.8 (B-7), 25.3 (C-11), 23.5 ($COCH_2\underline{CH_2}$), 21.1 (D-15), 13.4 (C/D-18) ppm.

***m/z* HRMS (ESI$^+$)** calcd. for $C_{32}H_{41}N_2O_4S$ ($[M+H]^+$): 549.2782 found: 549.2783.

**$\nu_{max}$** (film), $cm^{-1}$: 3472 (N-H stretch), 2968 (C-H stretch), 1737 (C=O stretch), 1648 (C=O stretch).

### 7. Biological Labelling

#### 7.1 Buffer solutions

Unless otherwise stated, all solutions described were prepared using deionised water ($dH_2O$) filtered through a Millipore Milli-Q water purification system.

**Homogenate Buffer**: 50 mM Tris-Cl pH 7.4, 250 mM sucrose, 5 mM $MgCl_2$, 1 mM dithiothreitol (DTT)

**PBS**: 8 mM $Na_2HPO_4$, 150 mM NaCl, 3 mM KCl, 2 mM $KH_2PO_4$

**PBST**: 8 mM $Na_2HPO_4$, 150 mM NaCl, 3 mM KCl, 2 mM $KH_2PO_4$, 0.1% Tween 20.

**Reducing Sample Buffer**: 200 mM Tris-Cl pH 6.8, 30% glycerol, 9% SDS, 0.4% β mercaptoethanol, 0.1% bromophenol blue

#### 7.2 SDS-PAGE

Denatured samples were resolved on 10% or 12% acrylamide gel selected according to the mass of the protein(s) of interest. 12% acrylamide gel was composed of resolving gel: 1.3 mL, 1.5 M Tris-Cl pH 6.8, 1.5 mL 40% acrylamide/bis acrylamide (29:1), 2 mL $dH_2O$, 50 µL 10% SDS, 50 µL 10% ammonium persulfate, 5 µL tetramethylenediamine; and stacking gel: 630 µL 0.5 M Tris-Cl pH 6.8, 300 µL acrylamide/bis-acrylamide (29:1), 1.3 mL $dH_2O$, 25 µL 10% SDS, 25 µL ammonium persulfate, 2.5 µL tetramethylenediamine. 10% acrylamide gel was composed of resolving gel: 1.3 mL, 1.5 M Tris-Cl pH 6.8, 1.25 mL 40% acrylamide/bis acrylamide (29:1), 2.25 mL $dH_2O$, 50 µL 10% SDS, 50 µL 10% ammonium persulfate, 5 µL tetramethylenediamine; and stacking gel: 630 µL 0.5 M Tris-Cl pH 6.8, 300 µL acrylamide/bis-acrylamide (29:1), 1.3 mL $dH_2O$, 25 µL 10% SDS, 25 µL ammonium persulfate, 2.5 µL tetramethylenediamine.

Samples were loaded along with Fisher EZ-Run PreStained Recombinant Protein Ladder (Fisher Scientific) and resolution was achieved at 150 V for 80 min in running buffer (25 mM Tris, 190 mM Glycine, 1% SDS) using a PowerPac Basic Power Supply (Bio-Rad, California USA).

#### 7.3 Silver Staining

SDS-PAGE gels were treated with fixative solution (40% EtOH, 10% AcOH) for 1 h at ambient temperature. Gels were washed with 20% EtOH solution (2 x 10 min), then rehydrated in $dH_2O$ (2 x 10 min). Gels were sensitised in 0.02% $Na_2S_2O_3$ solution for 50 s and immediately washed with $dH_2O$ (2 x 1 min). Gels were incubated in a solution of 12 mM $AgNO_3$(aq.) solution containing 0.02% formaldehyde at 4 °C for 1 h. Gels were washed with $dH_2O$ (2 x 20 s) and treated with developer solution (3 potassium carbonate, 0.05% formaldehyde). Development was halted with 5% AcOH. Stained gels were visualised using a Typhoon TLA9500 laser scanner (GE Healthcare, Illinois USA).

### 7.4 Coomassie Staining

SDS-PAGE gels were treated with 2.4 mM Coomassie blue in 10% AcOH, 50% MeOH, for 3 h. Gels were washed with 10% AcOH, 10% MeOH solution (4 x 20 min). Stained gels were visualised using a ChemiDoc MP Imaging System (Bio-Rad, California USA) in the white light channel.

### 7.5 Synthesis of functionalised bovine serum albumin (BSA)

#### 7.5.1 Synthesis of polyolefin-BSA

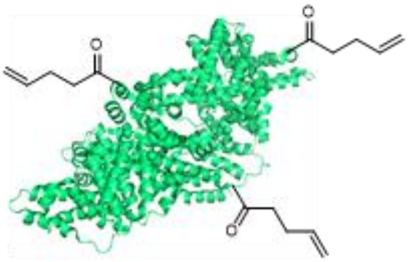

Bovine serum albumin (BSA) (300 µg, 4.51 nmol, 1.0 equiv.) in homogenate buffer pH 10.2 (500 µL) was treated with 20 mM 2,5-dioxopyrrolidin-1-yl pent-4-enoate, **S5** (33.9 µL, 678 nmol, 150 equiv.) in DMSO and incubated at 37 °C for 5 h. The mixture was desalted using a NAP-5 column according to the manufacturer's instructions and eluted with column buffer. The pH was adjusted to pH 6-7 through portion-wise addition of 1 M HCl. Protein concentration of the eluent was measured on a Nanodrop 1000 Spectrophotometer (0.25 mg/mL, 1000 µL).

#### 7.5.2 Synthesis of polyalkyl-BSA

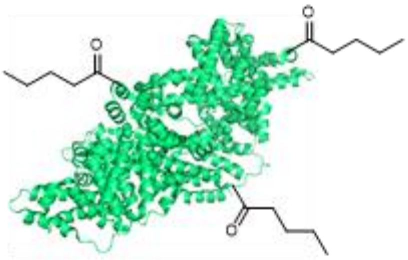

Bovine serum albumin (BSA) (300 µg, 4.51 nmol, 1.0 equiv.) in homogenate buffer pH 10.2 (500 µL) was treated with 20 mM 2,5-dioxopyrrolidin-1-yl pentanoate, **S6** (33.9 µL, 678 nmol, 150 equiv.) in DMSO and incubated at 37 °C for 5 h. The mixture was desalted using a NAP-5 column according to the manufacturer's instructions and eluted with column buffer. The pH was adjusted to pH 6-7 through portion-wise addition of 1 M HCl. Protein concentration of the eluent was measured on a Nanodrop 1000 Spectrophotometer (0.24 mg/mL, 1000 µL).

#### 7.5.3 Thiol-ene labelling of functionalised BSA

Functionalised BSA (16 µL, 0.25 mg/mL in homogenate buffer) was diluted to a total volume of 50 µL with homogenate buffer in glass microvials (Supelco). Samples were treated with a solution of AbzSH **1** (25 mM),

DPAP (15 mM) and MAP (100 mM) in DMSO-d$H_2O$ (5:1, 4.5 µL). Samples were sparged with nitrogen for 30 s and then irradiated at 365 nm for 5 min in a LZC photoreactor oven (Luzchem, Ontario Canada). Upon completion, samples were treated with reducing sample buffer (30 µL) and heated to 95 °C for 5 min. Proteins were resolved by 10% SDS-PAGE and visualised by silver staining and using a Chemidoc XRS+ equipped with a UV-lamp centred at 302 nm (BioRad, California USA).

### 7.6 Genetic incorporation of S-allyl-L-cysteine into LmrR

#### 7.6.1 Selection of the orthogonal synthetase/tRNA pair

Four orthogonal aminoacyl-tRNA synthetase/tRNA pairs (OP1-OP4) were screened for incorporation of S-allyl-L-cysteine (SAC) using an sfGFP reporter containing an in-frame amber codon at position 150 (sfGFP-150TAG). Whole-cell fluorescence was normalised to OD600. OP2 produced the strongest SAC-dependent response and was selected for expression of LmrR V15SAC (***Figure S1***).

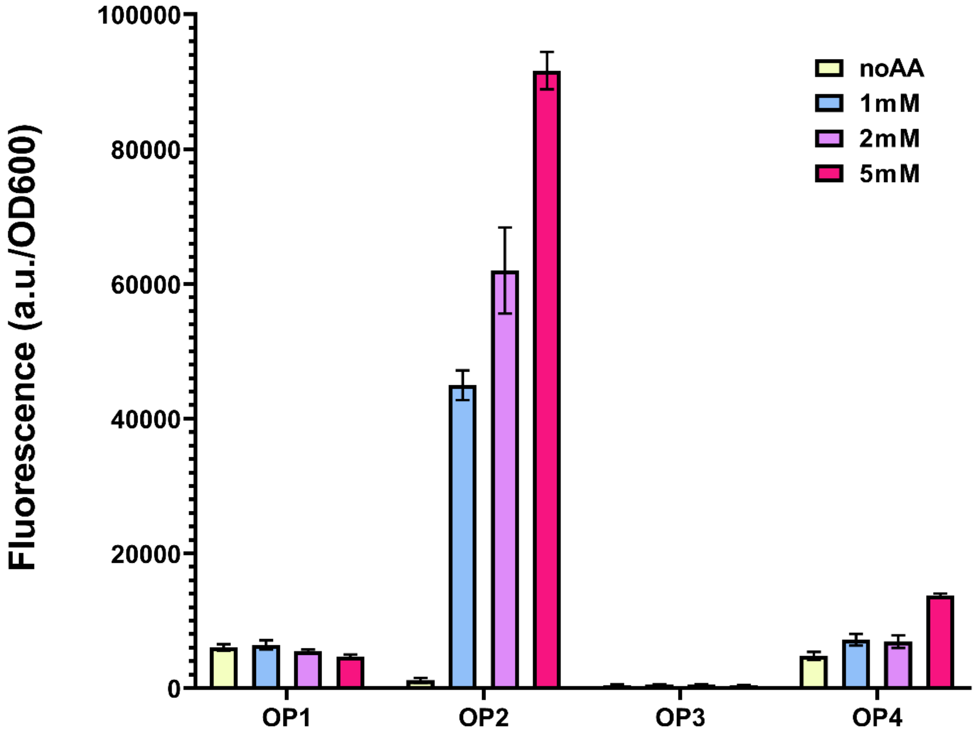

**Figure S1.** Screening of OP1-OP4 for SAC-dependent amber suppression using sfGFP-150TAG. Cultures were analysed without SAC (noAA) or with 1, 2 or 5 mM SAC. Fluorescence was normalised to OD600. The number of biological replicates and the error-bar definition require confirmation before publication.

#### 7.6.2 Expression and purification of LmrR V15SAC

The LmrR V15TAG expression construct was available from previous work in the Drienovska laboratory. Chemically competent Escherichia coli BL21(DE3) cells were co-transformed with the ampicillin-resistant LmrR V15TAG expression plasmid and the spectinomycin-resistant OP2 plasmid. A selected colony was used to inoculate LB medium (5 mL) containing ampicillin (100 µg/mL) and spectinomycin (100 µg/mL) and was grown overnight at 37 °C with shaking at 130 rpm. The preculture was transferred to LB medium (500 mL in a 5 L Erlenmeyer flask) containing the same antibiotics and grown at 37 °C until OD600 reached 0.6-0.8. SAC (10 mM final concentration) and IPTG (1 mM final concentration) were added, and the culture was incubated overnight at 28 °C with shaking at 135 rpm. Wild-type LmrR was expressed in parallel with ampicillin selection but without OP2 or SAC.

Cells were harvested by centrifugation (4000 rpm, 45 min, 4 °C) and resuspended in lysis buffer (50 mM potassium phosphate, 500 mM NaCl, 0.1 mM phenylmethylsulfonyl fluoride, pH 8.0). Cells were disrupted by sonication on ice (5 s on/5 s off, 5 min total sonication time, 35% amplitude), and insoluble material was removed by centrifugation (20000 rpm, 45 min, 4 °C). The clarified lysate was incubated with Strep-Tactin resin, pre-equilibrated with four column volumes of wash buffer (20 mM Tris-HCl, 300 mM NaCl, pH 8.0), for 30 min on ice. The resin was washed with ten column volumes of wash buffer, and bound protein was eluted in 3 mL of 20 mM Tris-HCl, 300 mM NaCl and 2.5 mM desthiobiotin, pH 8.0. The protein was desalted into 50 mM HEPES, 150 mM NaCl, pH 8.0, flash-frozen in liquid nitrogen and stored at -20 °C.

Protein concentration was determined by absorbance at 280 nm using a molar extinction coefficient of 25440 M-1 cm-1, calculated for monomeric LmrR. A typical purified yield of 14 mg L-1 culture was obtained.

### 7.7 Thiol-ene labelling of LmrR V15SAC

#### 7.7.1 Labelling of purified LmrR

Thiol-ene labelling reactions were performed in 50 mM HEPES, 150 mM NaCl, pH 7.4, in a total volume of 30 µL in PCR tubes. Unless stated otherwise, each reaction contained 6 µg purified protein. Stock solutions of AbzSH, 2,2-dimethoxy-2-phenylacetophenone (DPAP) and 4'-methoxyacetophenone (MAP) were prepared at 50 mM in DMSO; Irgacure 2959 was prepared at 14 mM in Milli-Q water. For the initial comparison of photoinitiator systems, LmrR V15SAC or wild-type LmrR was treated with AbzSH (2.0 mM) and either DPAP/MAP (0.7/4.9 mM) or Irgacure 2959 (2.0 mM), followed by irradiation at 365 nm for 30 min using a UVP UVL-15 lamp (Analytik Jena; 4 W, 254/365 nm) operated at 365 nm. PCR tubes were positioned directly beneath the lamp in its holder, at the fixed distance set by the lamp housing, without additional spacing. Control reactions lacking AbzSH, photoinitiator or UV irradiation were prepared in parallel.

#### 7.7.2 Optimisation of probe concentration and irradiation time

For the initial concentration screen, LmrR V15SAC was treated with AbzSH at final concentrations of 0.1, 0.2, 0.5, 1.0 or 2.0 mM. DPAP and MAP were scaled proportionally with AbzSH, and samples were irradiated at 365 nm for 30 min. A two-dimensional optimisation was subsequently performed using AbzSH at 0.2, 0.5, 1.0 or 2.0 mM and irradiation times of 2.5, 5, 10, 20 or 30 min. A concentration of 0.5 mM AbzSH produced a clear band with limited background in the purified-protein screen, whereas 1.0 mM gave a stronger signal and was selected as the working concentration for subsequent experiments. The conditions reported in the manuscript were 1.0 mM AbzSH, 0.4 mM DPAP and 2.5 mM MAP with irradiation at 365 nm for 10 min.

#### 7.7.3 Labelling in E. coli cell-free lysate

Cell-free lysate containing LmrR V15SAC was prepared using the expression and lysis procedure described above. Following sonication and clarification by centrifugation, purification was omitted and the clarified lysate was used directly. Each 30 µL labelling reaction contained 6 µL clarified lysate; AbzSH, DPAP/MAP and reaction buffer were added to the final volume. Lysate samples were treated with AbzSH (0.5, 1.0 or 2.0 mM) and proportionally scaled DPAP/MAP under the selected reaction conditions. Samples lacking AbzSH or UV irradiation were included as negative controls. Approximately equal amounts of LmrR V15SAC were present across the lysate samples, as judged by Coomassie-stained SDS-PAGE.

##### 7.7.4 SDS-PAGE and in-gel fluorescence

After labelling, 10 µL of each reaction (approximately 2 µg purified protein where applicable) was mixed with 2.5 µL of 5x reducing SDS sample buffer and heated at 95 °C for 5 min. The 5x buffer was prepared, per 50 mL, from Tris-Cl (1.875 g), glycerol (25 mL), SDS (5 g), 2-mercaptoethanol (12.5 mL) and bromophenol blue, adjusted to pH 6.8 and brought to volume with Milli-Q water. Samples were resolved on 4-20% gradient precast polyacrylamide gels (12-well) at 140 V for 70 min. Gels were washed three times with deionised water (5 min per wash), and in-gel fluorescence was recorded at 305 nm using an Azure Biosystems 400 imaging system. Proteins were subsequently stained with InstantBlue Coomassie Protein Stain (Abcam).

#### 7.8 Intact-protein LC-MS analysis

##### 7.8.1 LC-MS method

LC–MS (quadrupole). Protein characterisation was performed using an Agilent LC/MSD XT ESI-Quadrupole system equipped with an electrospray ionisation source. Protein samples (5 µL, approximately 10 µM) were injected onto an Agilent AdvanceBio RP-mAb C4 column (2.1 × 50 mm). Mobile phase A consisted of water containing 0.1% formic acid (v/v), while mobile phase B consisted of isopropanol/acetonitrile/water (80:10:10, v/v/v) containing 0.1% formic acid. Proteins were separated using a linear gradient from 0 to 95% B over 15 min at a flow rate of 0.6 mL $min^{-1}$. Mass spectra were acquired over an m/z range of 0–3000 at a scan rate of 1.0 Hz, with a capillary voltage of 3000 V. Spectra were deconvoluted using a maximum-entropy algorithm, with $H^+$ as the charge carrier and a deconvolution mass range of 10,000–100,000 Da.

LC–MS (TOF).

High-resolution intact-protein LC–MS analysis was performed using an Agilent LC/MSD TOF system (G6230B) equipped with an electrospray ionisation source. Protein samples (1 µL, approximately 10 µM) were injected onto an Agilent AdvanceBio RP-mAb C4 column (2.1 × 50 mm). Mobile phases A and B were as described above. Proteins were separated using a linear gradient from 0 to 95% B over 10 min at a flow rate of 0.6 mL $min^{-1}$. The first 2 min were diverted to waste. Mass spectra were acquired over an m/z range of 0–3000 at a scan rate of 1.0 Hz, with a capillary voltage of 3500 V. Spectra were deconvoluted using the maximum-entropy algorithm in Agilent BioConfirm software (version 12.0), with $H^+$ as the charge carrier and a deconvolution mass range of 10,000–100,000 Da.

#### 7.8.2 Confirmation of SAC incorporation

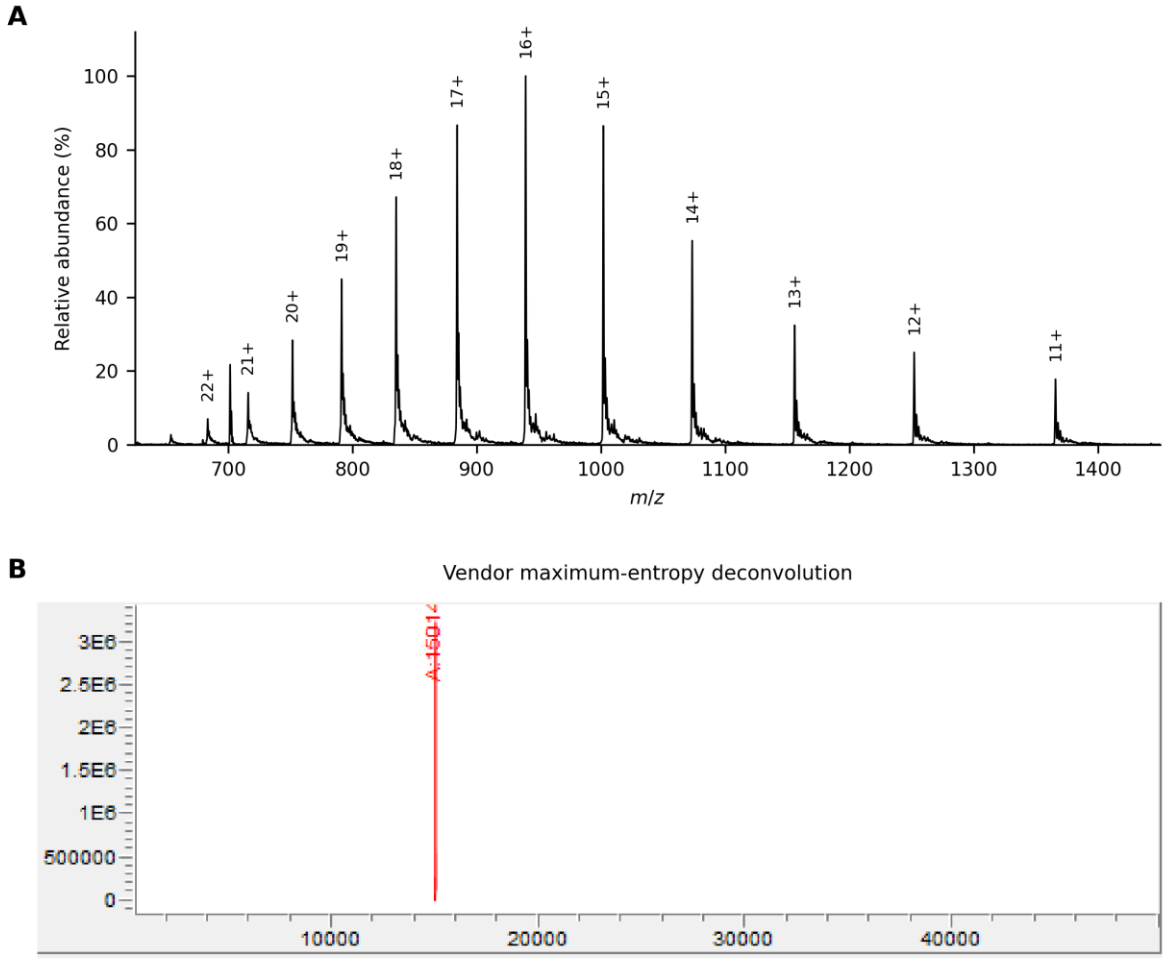

**Figure S2.** Intact-protein LC-MS confirmation of SAC incorporation in purified LmrR V15SAC. (A) Positive-ion ESI-MS charge-state envelope replotted from the machine-exported raw spectrum; charge assignments are shown above the principal peaks. (B) Vendor-generated maximum-entropy deconvoluted mass spectrum of LmrR V15SAC. The observed neutral mass was 15014 Da, in agreement with the calculated mass of 15013.2 Da.

#### 7.8.3 TEC labelling initiated with DPAP/MAP

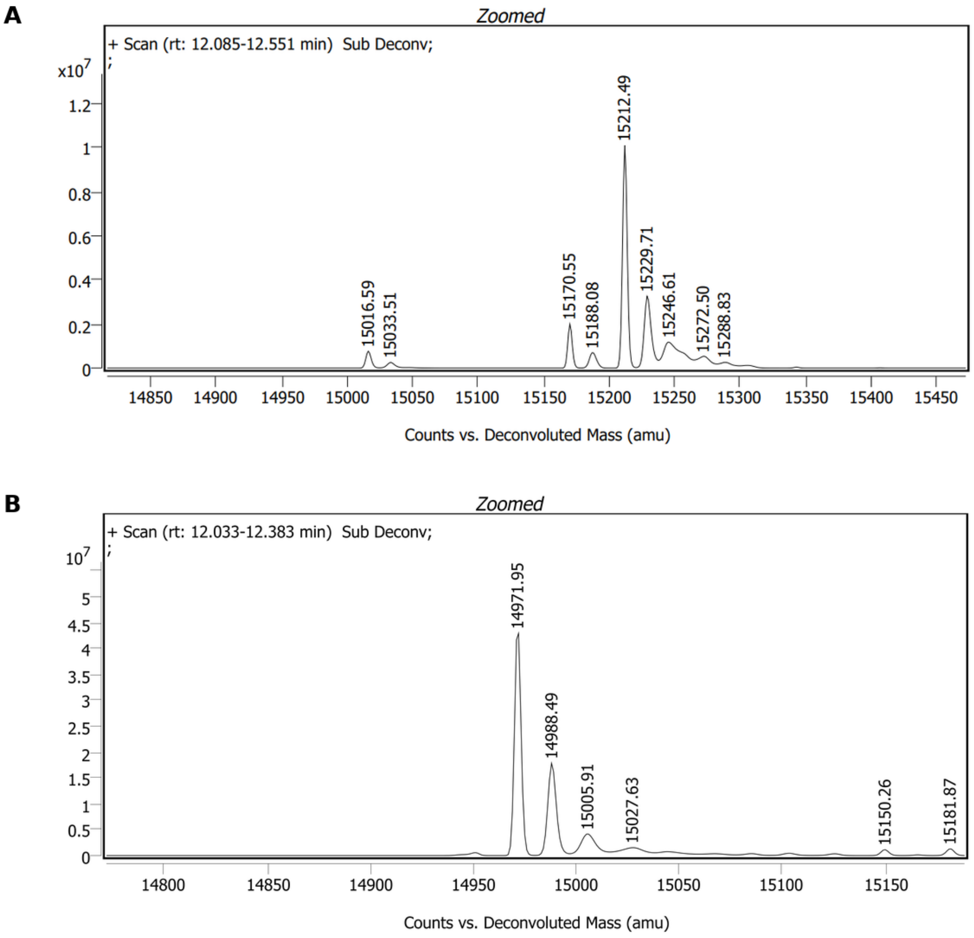

**Figure 3(b). Raw** deconvoluted intact-protein mass spectra following TEC treatment with AbzSH and DPAP/MAP. (A) LmrR V15SAC: signals at 15016 Da and 15212 Da correspond to unmodified and AbzSH-labelled protein, respectively. The +196 Da shift is consistent with attachment of one AbzSH molecule, and the labelled species was the dominant signal. (B) Wild-type LmrR treated under the same conditions: the major signal at 14972 Da corresponds to unmodified protein; no +196 Da product was detected.

#### 7.8.4 TEC labelling initiated with Irgacure 2959

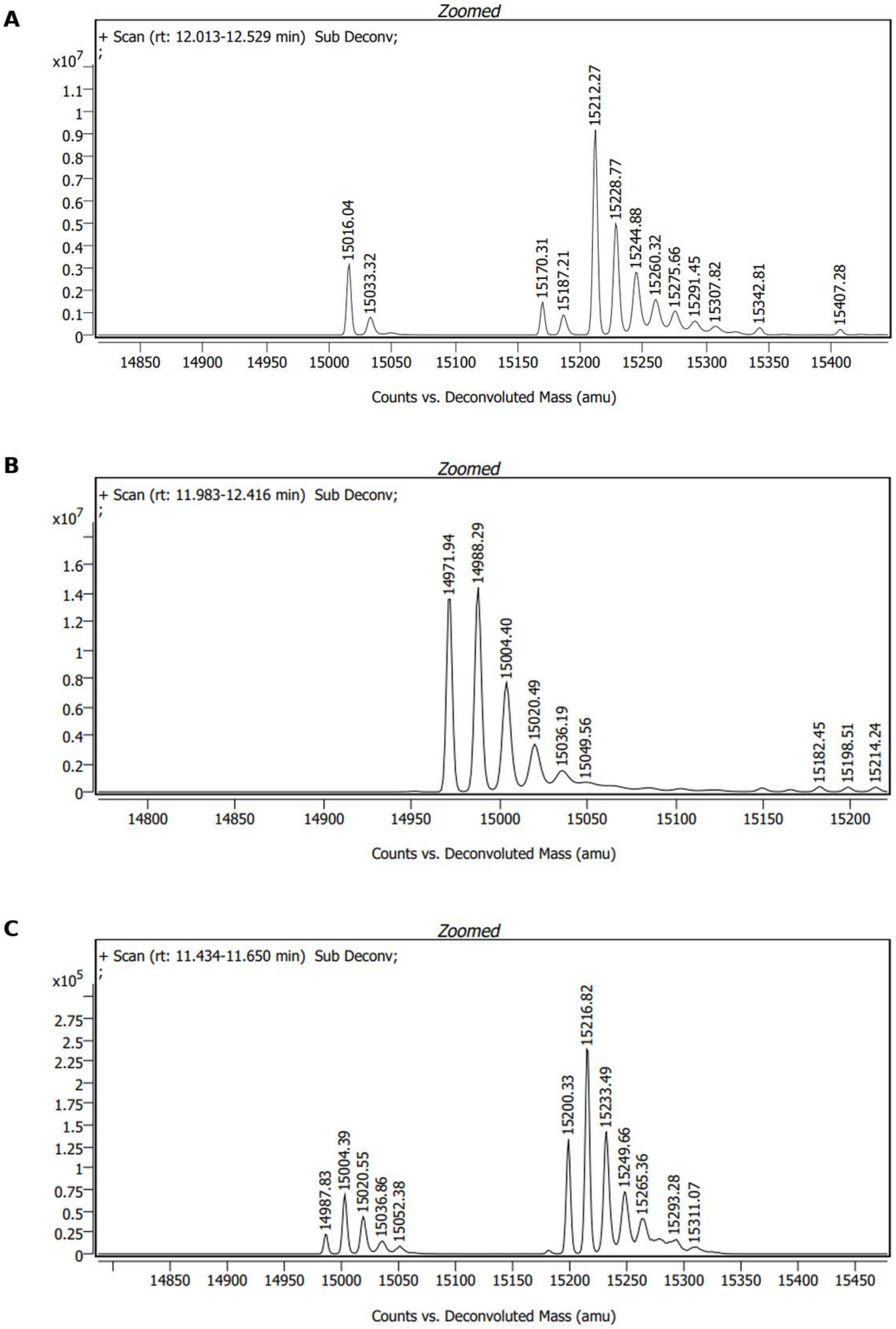

**Figure S3.** Deconvoluted intact-protein mass spectra following Irgacure 2959-initiated TEC treatment. (A) LmrR V15SAC: signals at 15016 Da and 15212 Da correspond to unmodified and AbzSH-labelled protein, respectively; additional lower-intensity species were also present. (B) Wild-type LmrR, principal retention-time window: the dominant signals at 14972 and 14988 Da correspond to unmodified protein and a +16 Da species. (C) A separate lower-intensity retention-time window contained signals at approximately 15200 and 15217 Da. These species do not correspond to a simple +196 Da AbzSH addition to wild-type LmrR and were not assigned.

#### 7.8.5 Positive-ion ESI-MS charge-state envelopes for control samples

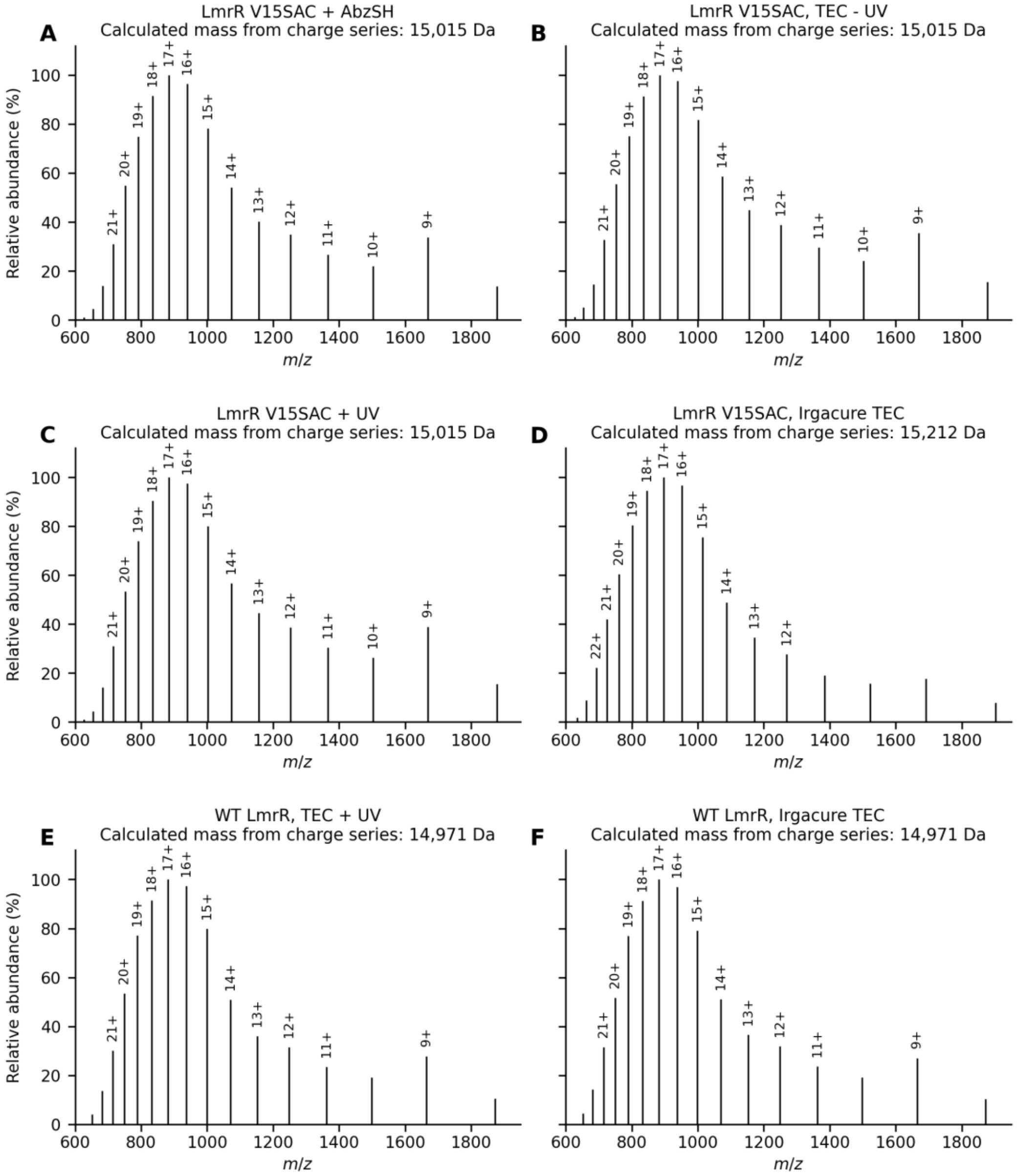

**Figure S4.** Positive-ion ESI-MS charge-state envelopes replotted from machine-exported Agilent BioConfirm centroided peak tables for the LmrR TEC control series. (A) LmrR V15SAC with AbzSH; (B) LmrR V15SAC under TEC conditions without UV irradiation; (C) LmrR V15SAC exposed to UV; (D) LmrR V15SAC following Irgacure 2959-initiated TEC; (E) wild-type LmrR under TEC conditions with UV irradiation; and (F) wild-type LmrR following Irgacure 2959-initiated TEC. Intensities were normalised to the base peak in each spectrum. Consecutive charge states were assigned from each centroided peak series, and the corresponding calculated neutral mass is displayed above each panel. These panels are direct charge-state plots and are not vendor maximum-entropy deconvolutions. Panel D shows the approximately +196 Da mass difference expected for attachment of one AbzSH molecule.

### 7.9 Gel images for LmrR labelling experiments

#### 7.9.1 Initial TEC labelling and reaction controls

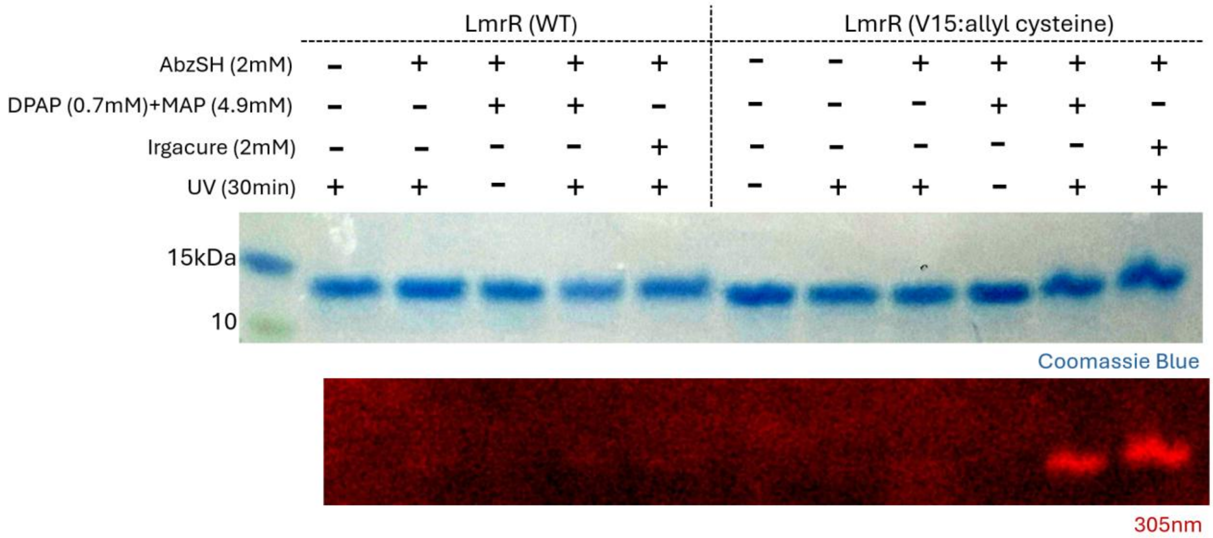

**Figure 3(a).** Annotated gel composite for the initial TEC labelling experiment. The Coomassie-stained gel is shown above the corresponding 305 nm in-gel fluorescence image. Lane conditions are displayed directly above the gels for wild-type LmrR and LmrR V15SAC, including the DPAP/MAP and Irgacure 2959 reaction controls.

#### 7.9.2 AbzSH concentration screen

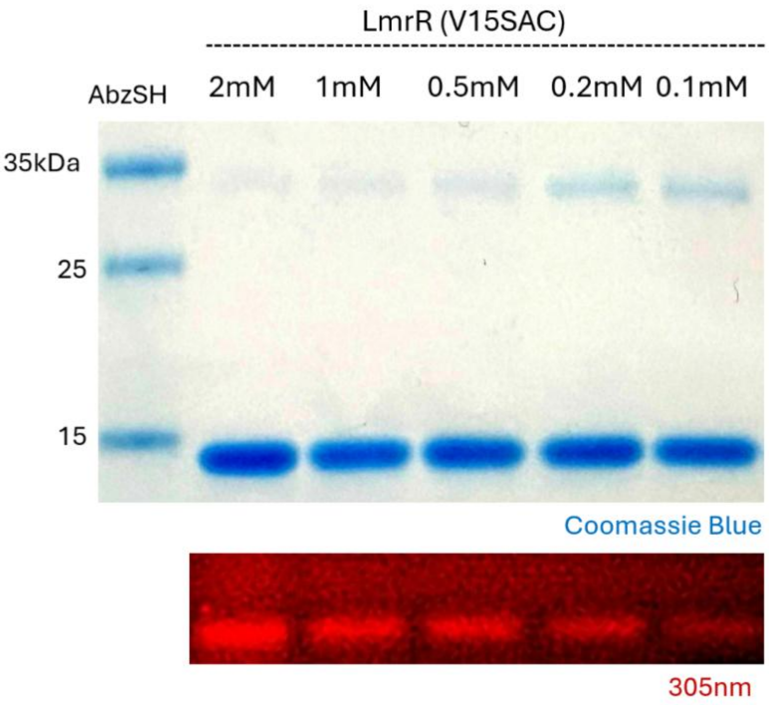

**Figure S5.** Annotated gel composite for the AbzSH concentration screen using purified LmrR V15SAC and 30 min UV irradiation. The Coomassie-stained gel is shown above the corresponding 305 nm in-gel fluorescence image. From left to right after the molecular-weight marker, AbzSH concentrations were 2.0, 1.0, 0.5, 0.2 and 0.1 mM, with DPAP/MAP scaled proportionally.

#### 7.9.3 Probe concentration and irradiation-time optimisation

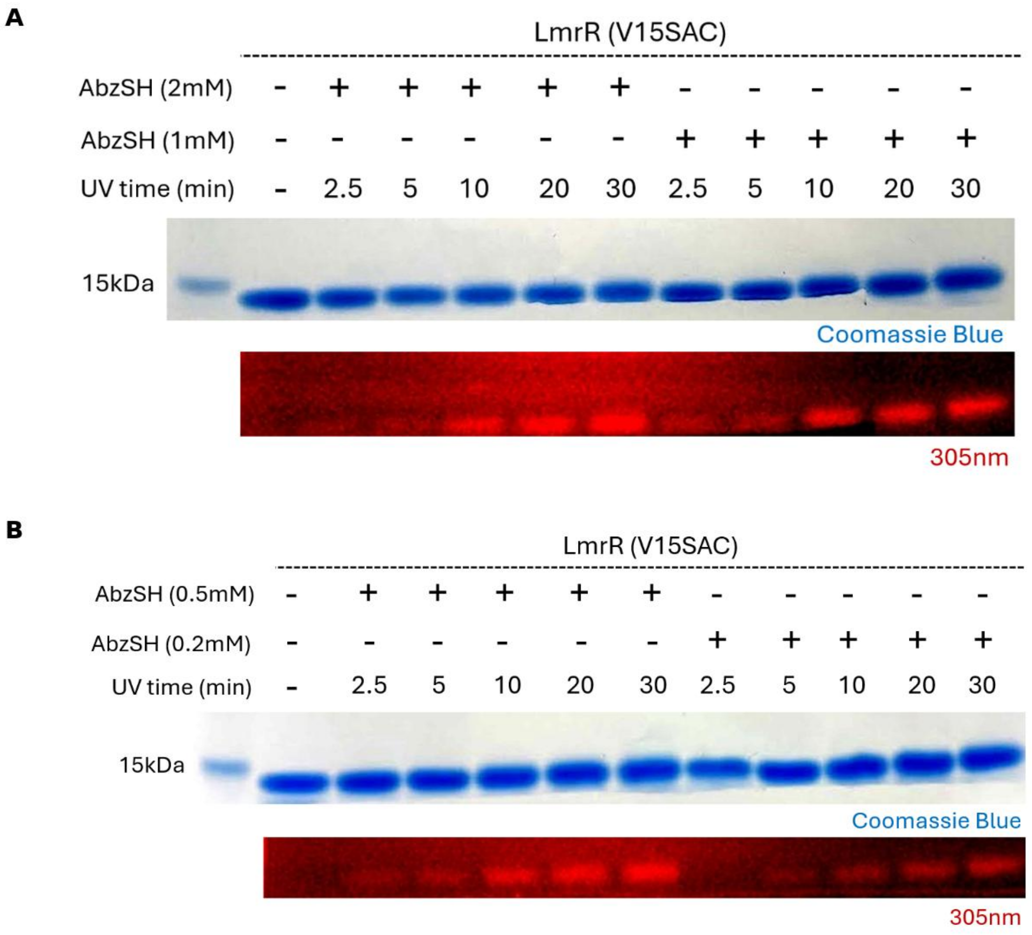

**Figure S6.** Annotated gel composites for the two-dimensional optimisation of AbzSH concentration and irradiation time. Each panel shows the Coomassie-stained gel above its corresponding 305 nm in-gel fluorescence image, with lane conditions displayed directly above the gels. (A) No-AbzSH/no-UV control followed by 2.0 and 1.0 mM AbzSH samples irradiated for 2.5, 5, 10, 20 or 30 min. (B) The corresponding series using 0.5 and 0.2 mM AbzSH. DPAP/MAP was scaled proportionally.

#### 7.9.4 TEC labelling in cell-free lysate

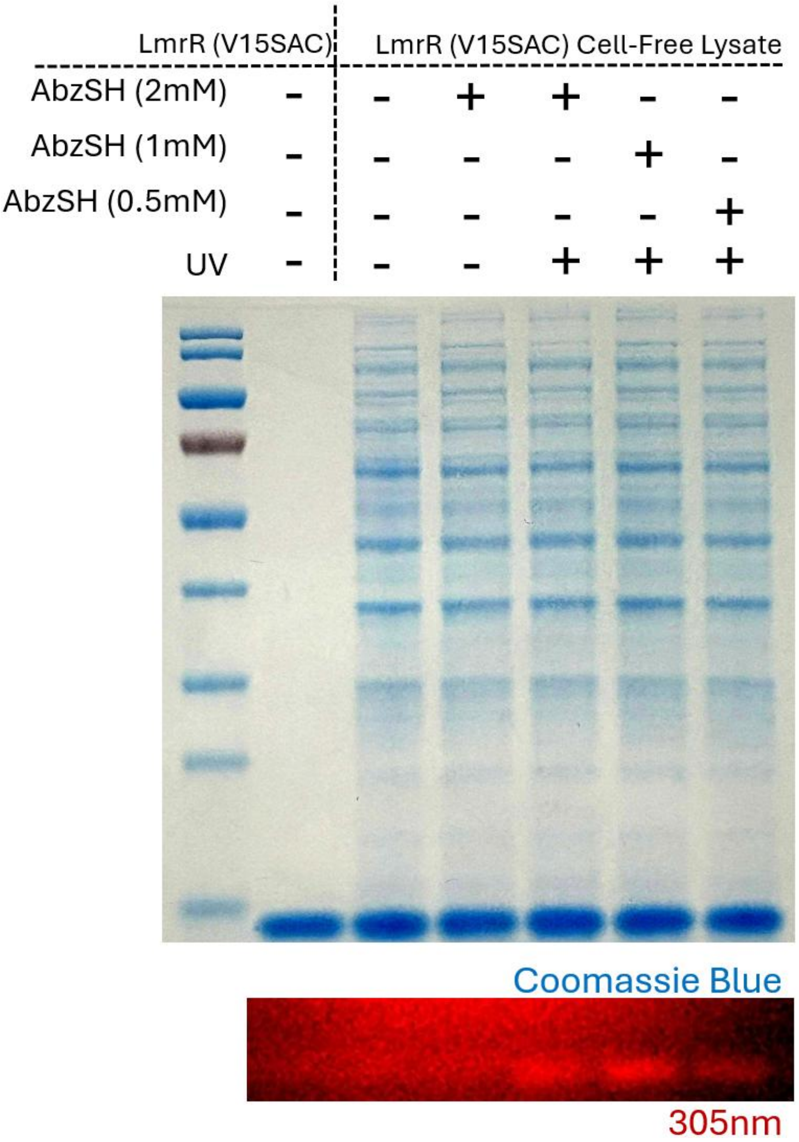

**Figure 4.** Annotated gel composite for TEC labelling in E. coli cell-free lysate containing LmrR V15SAC. The Coomassie-stained gel is shown above the corresponding 305 nm in-gel fluorescence image. From left to right after the molecular-weight marker: purified LmrR V15SAC without AbzSH or UV; lysate without AbzSH or UV; lysate with 2.0 mM AbzSH without UV; and complete lysate reactions with 2.0, 1.0 or 0.5 mM AbzSH and UV. Fluorescence was observed at the expected molecular weight of LmrR V15SAC without detectable labelling of other abundant lysate proteins.

### 7.10 Raw, unedited gel images

Raw, unedited gel images corresponding to the annotated initial TEC and cell-free lysate figures are provided below. The probe-concentration and irradiation-time optimisation data are presented only as annotated composites (***Figure S6***).

#### 7.10.1 Initial TEC labelling

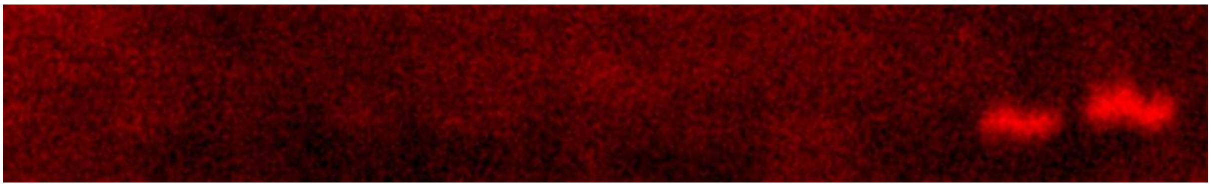

**Figure S7**. Edited picture of 305nm scan of TEC on LmrR (V15SAC) and LmrR WT

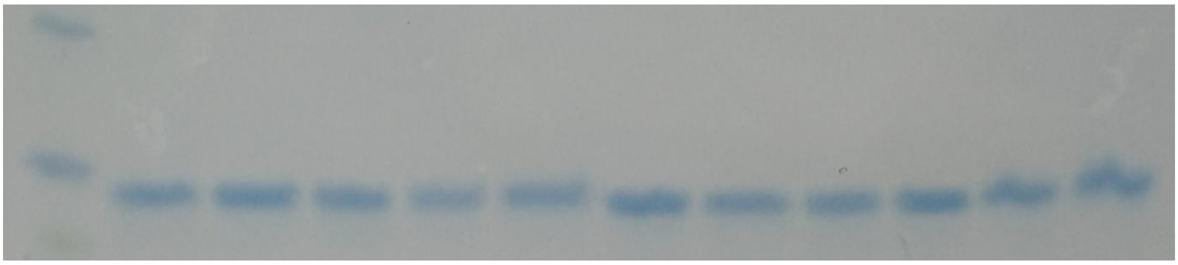

**Figure S8.** Raw, unedited cropped Coomassie-stained gel corresponding to the annotated initial TEC labelling figure above (***Figure 3***).

#### 7.10.2 Cell-free lysate labelling

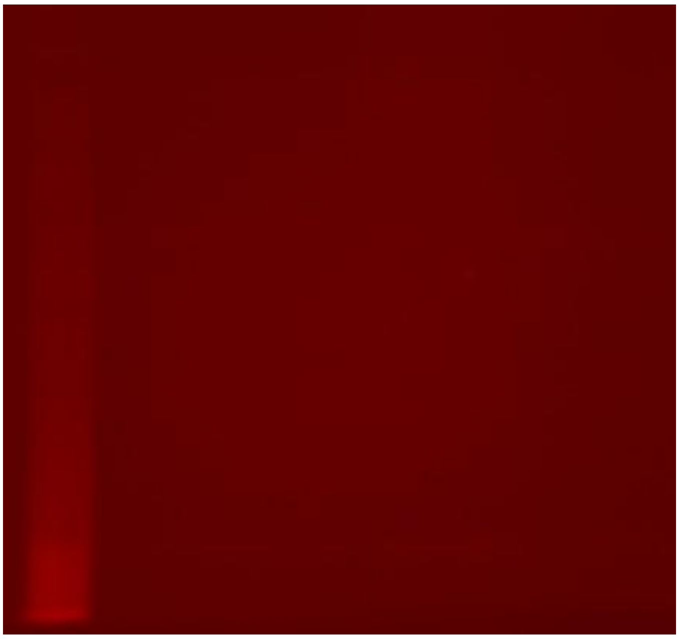

**Figure S9.** Raw, unedited 305 nm fluorescence image corresponding to the annotated cell-free lysate labelling figure above (***Figure 4***).

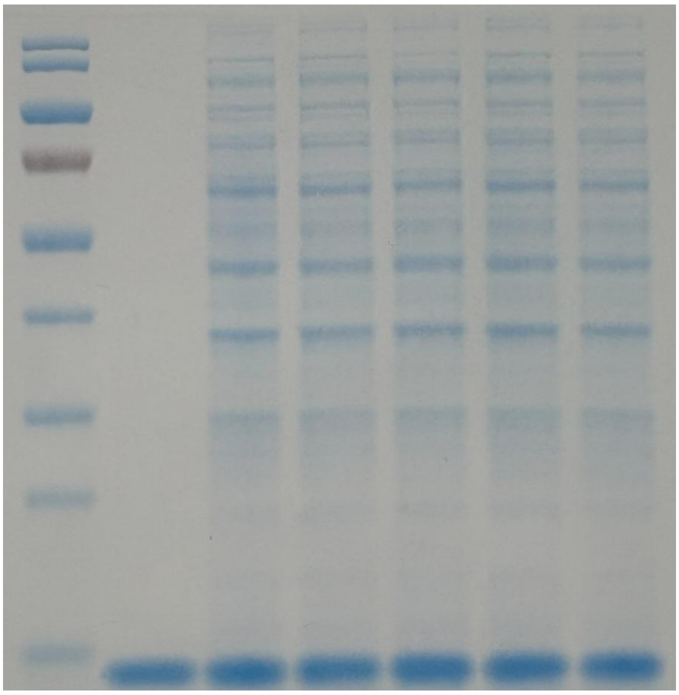

**Figure S10.** Raw, unedited Coomassie-stained gel corresponding to the annotated cell-free lysate labelling figure above (***Figure 4***).

### 8. Spectroscopic Data

**S2** (Boc-Abz)

$^1$H NMR $CDCl_3$

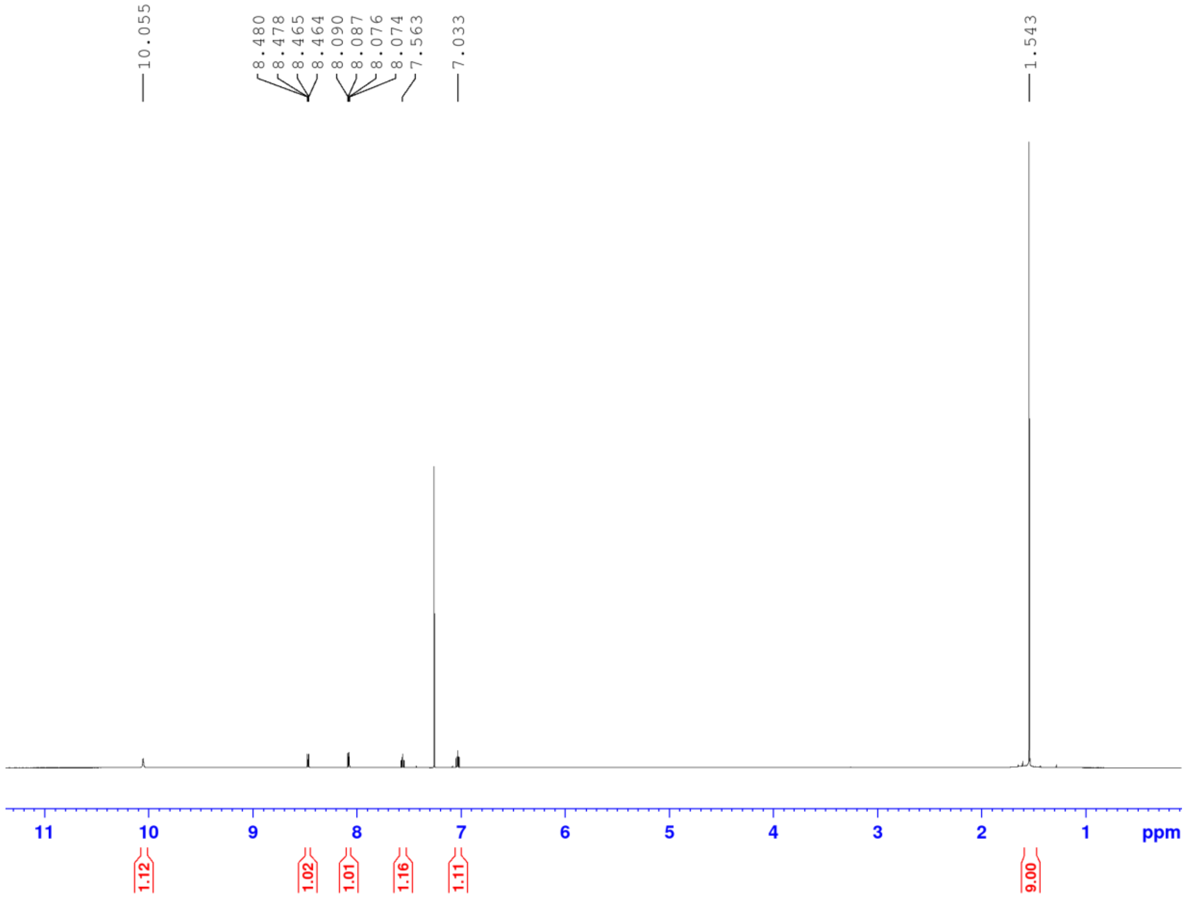

**S3** (Boc-Abz-STrt)

$^{1}$H NMR $CDCl_3$

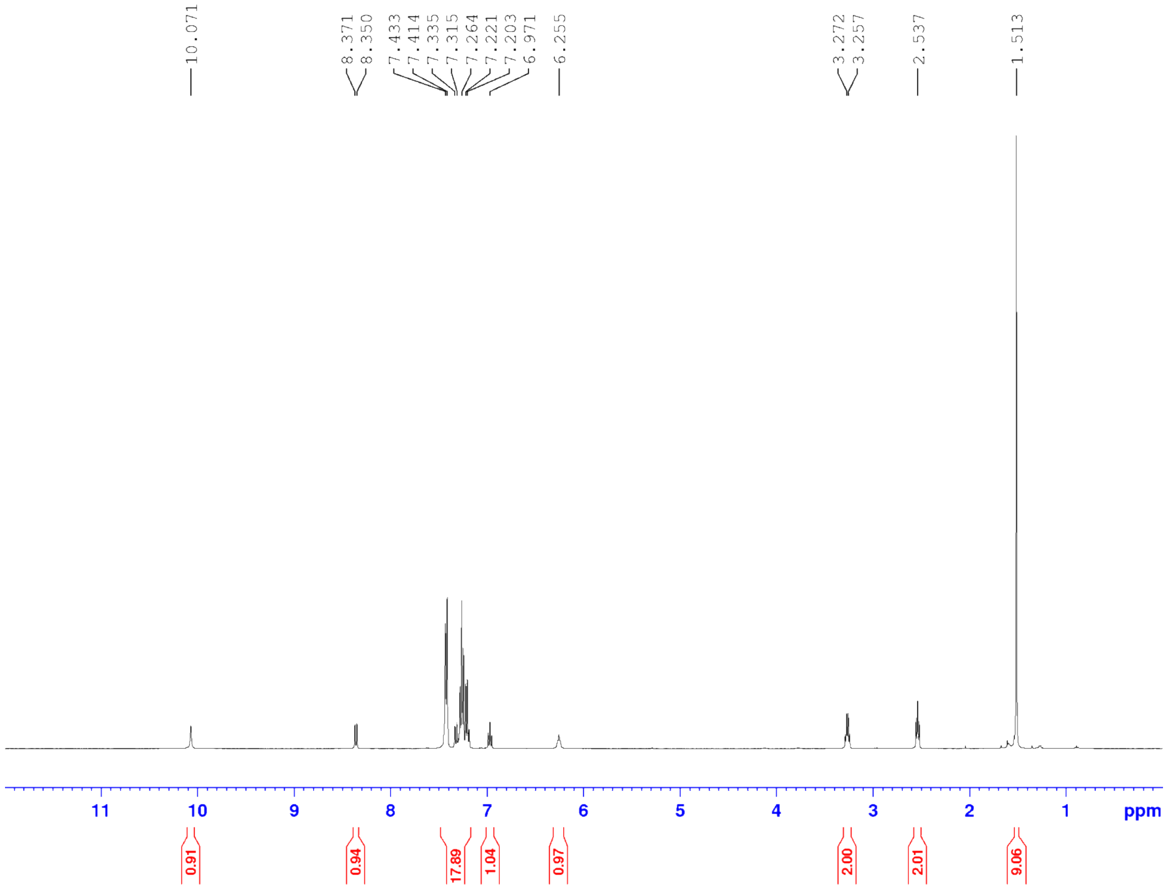

$^{13}$C NMR $CDCl_3$

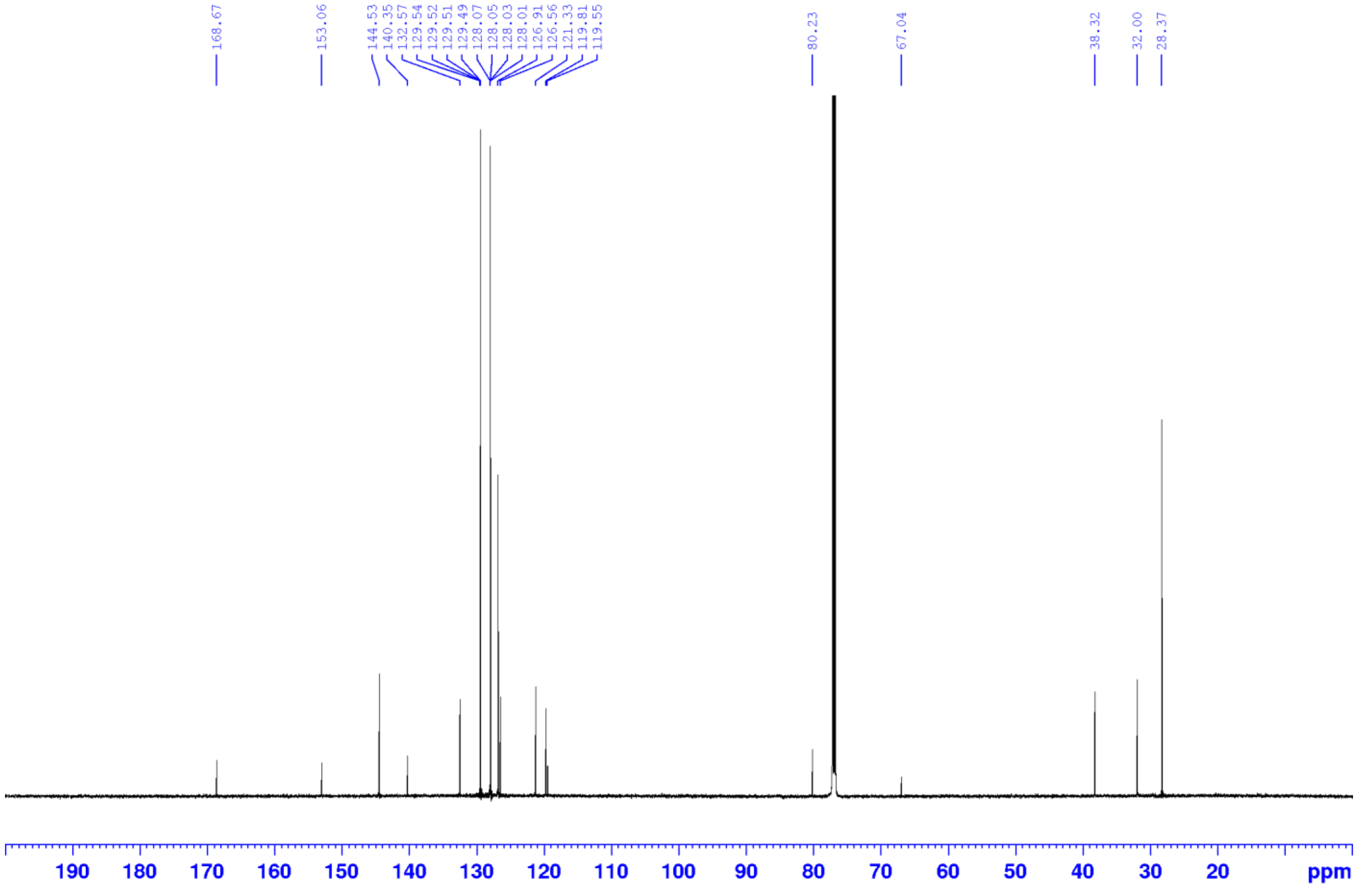

**1** (AbzSH)

$^1H$ NMR $CDCl_3$

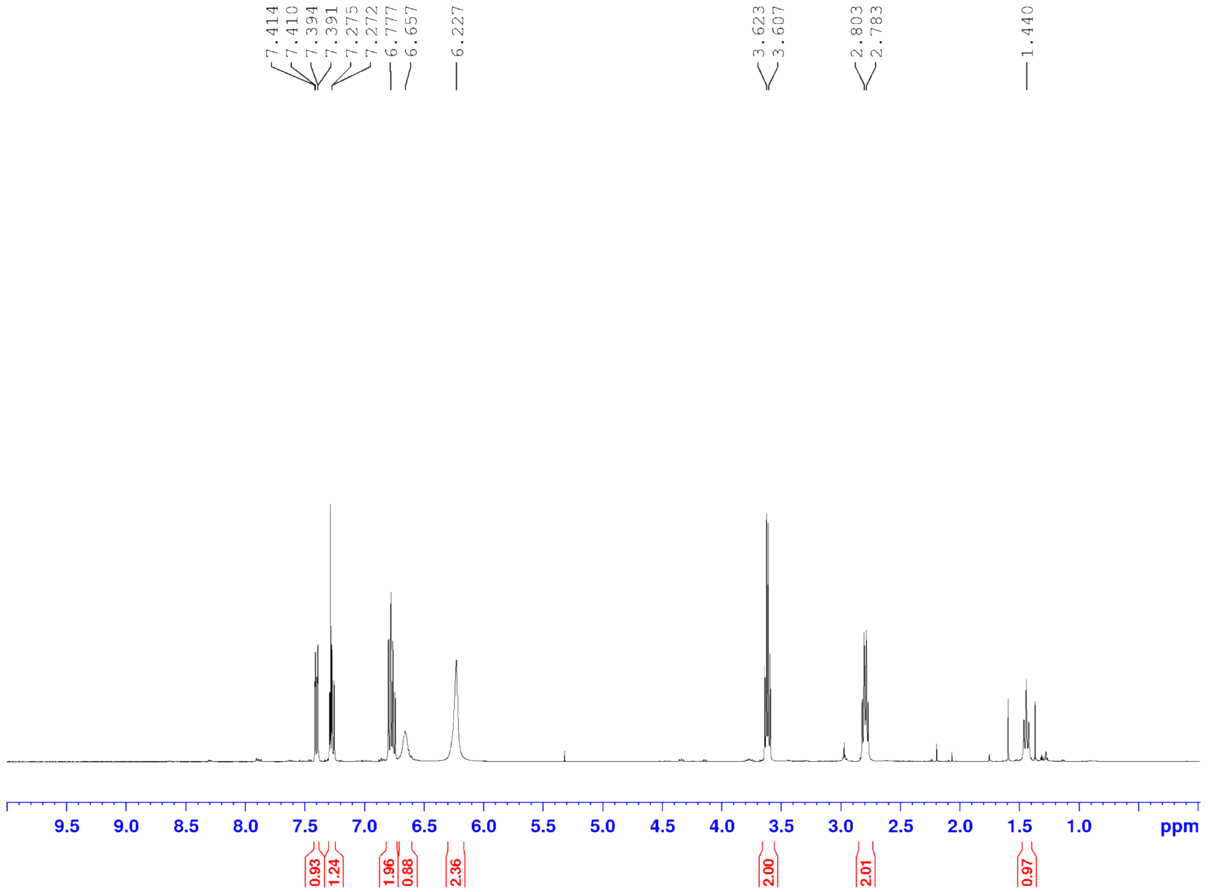

**S4** (Amine-free disulfide)

$^1$H NMR $CDCl_3$

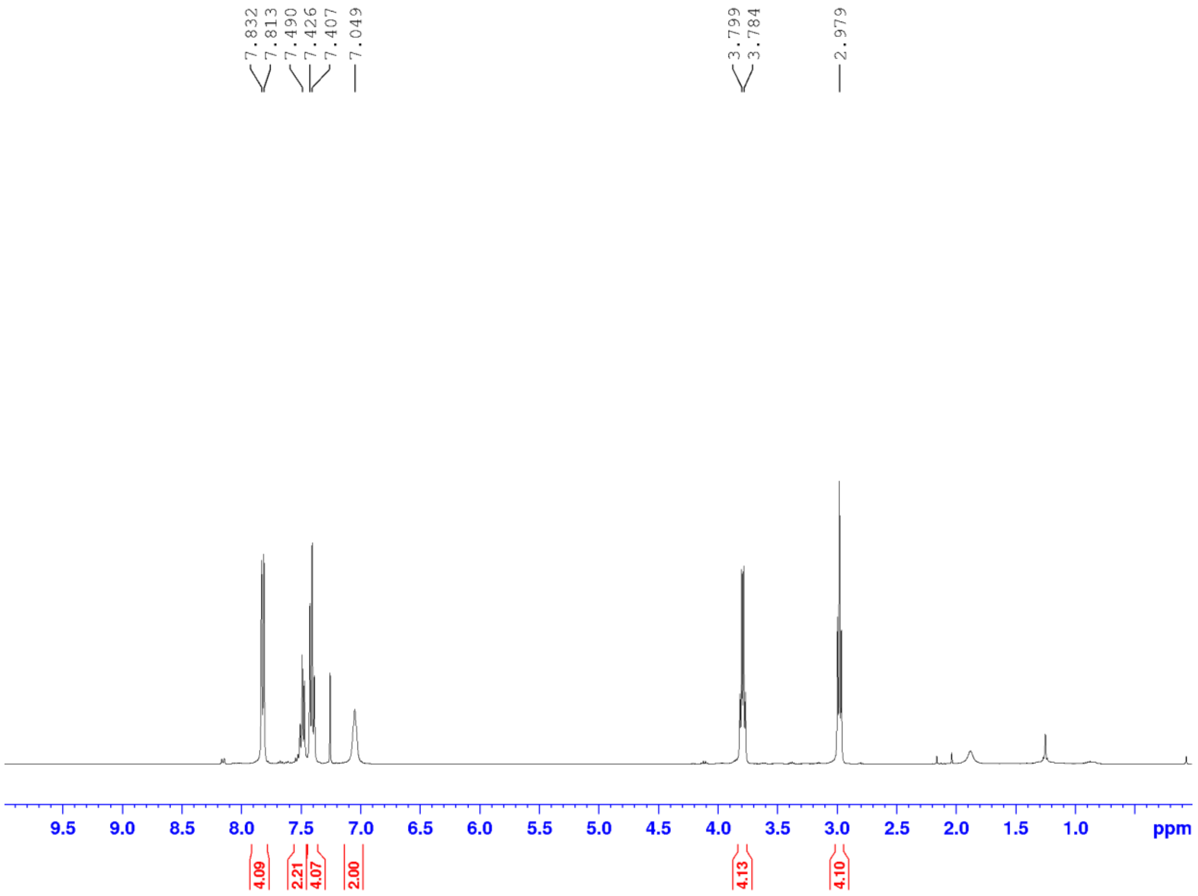

**3** (Amine-free thiol)

$^1H$ NMR $CDCl_3$

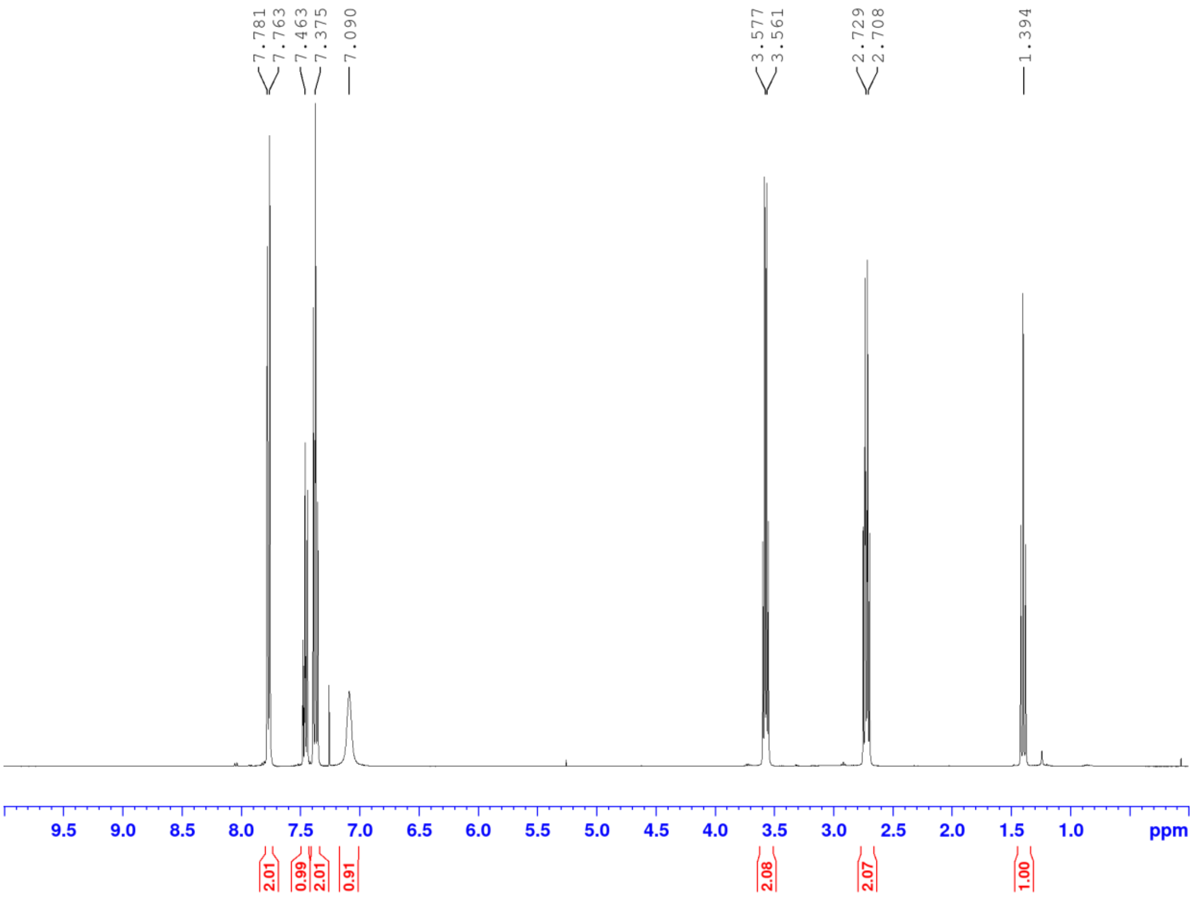

**2** (Sugar)

$^1H$ NMR $CDCl_3$

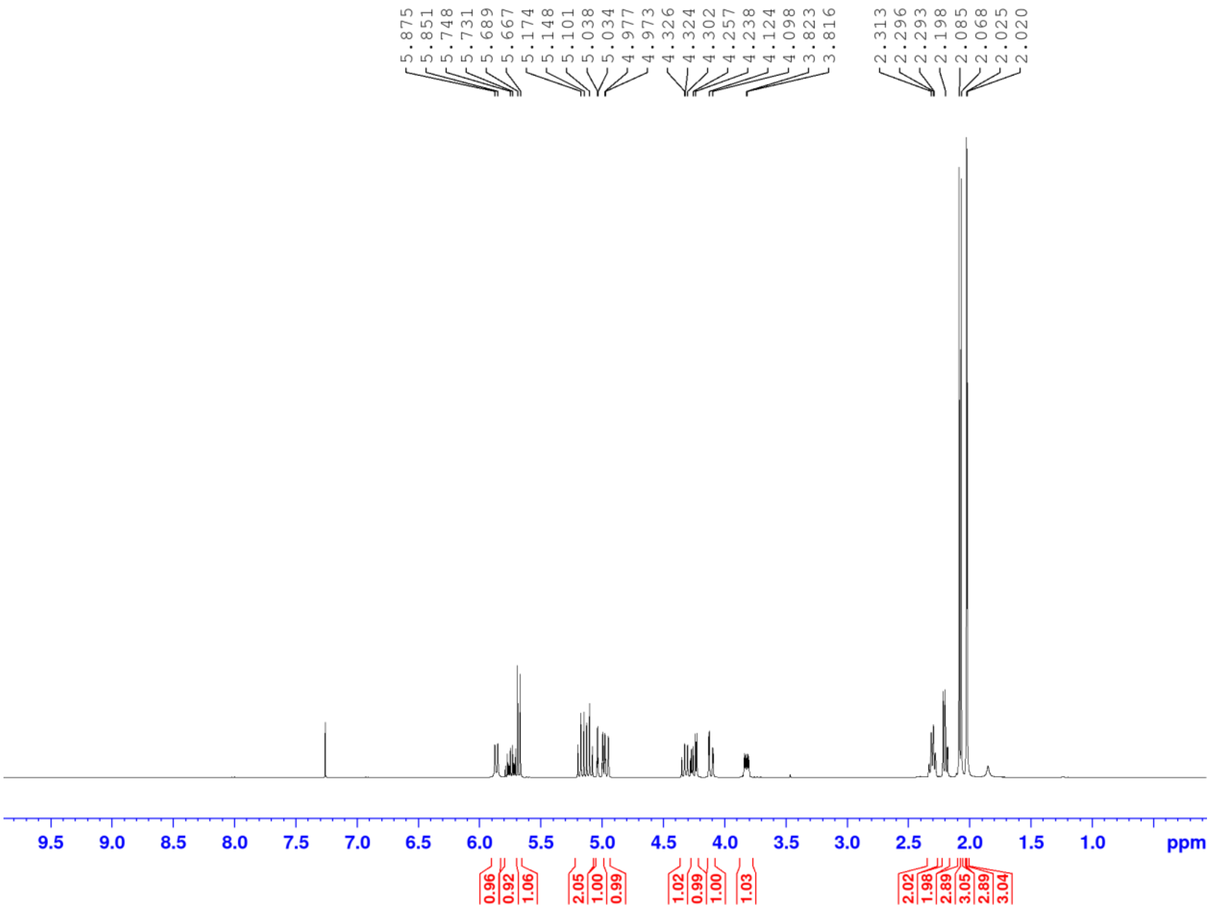

**7** (Estrone)

$^{1}H$ NMR $CDCl_3$

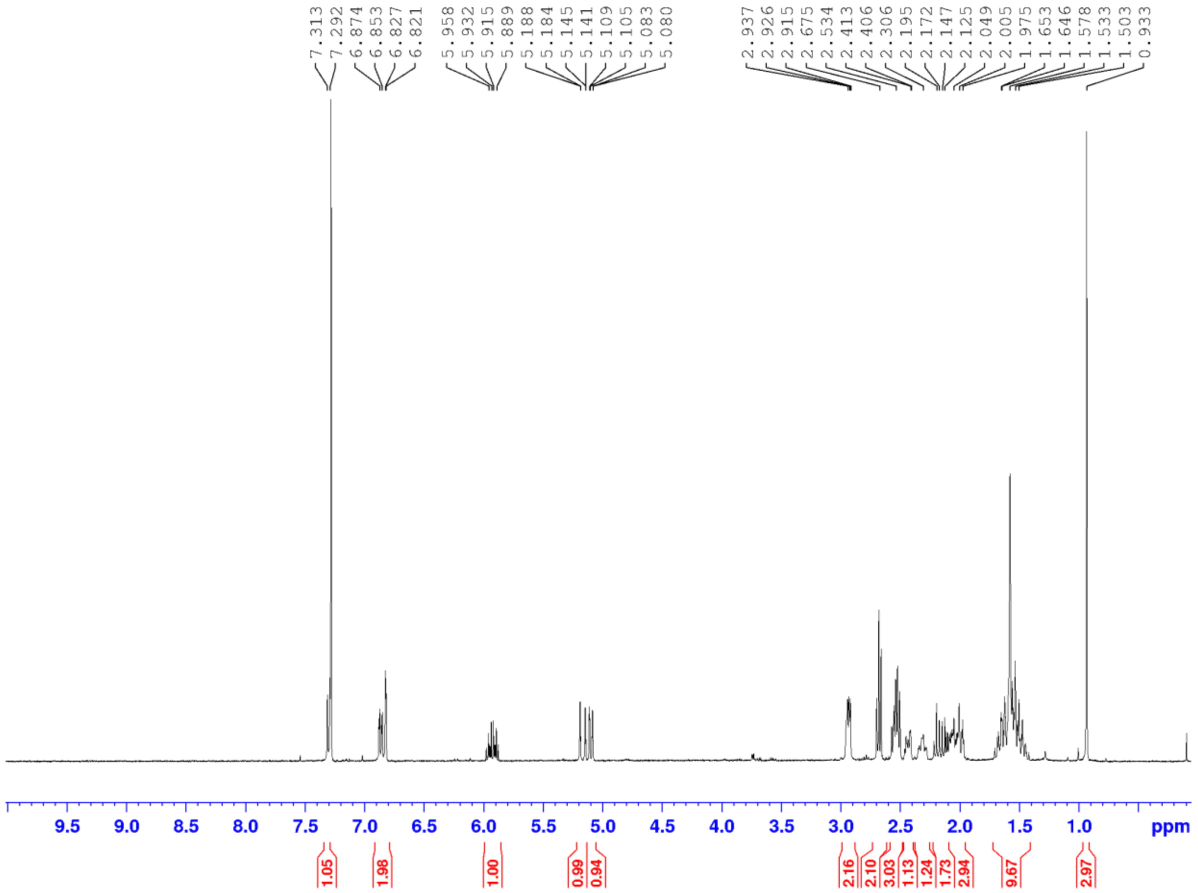

**S5** (NHS-alkene)

$^1H$ NMR $CDCl_3$

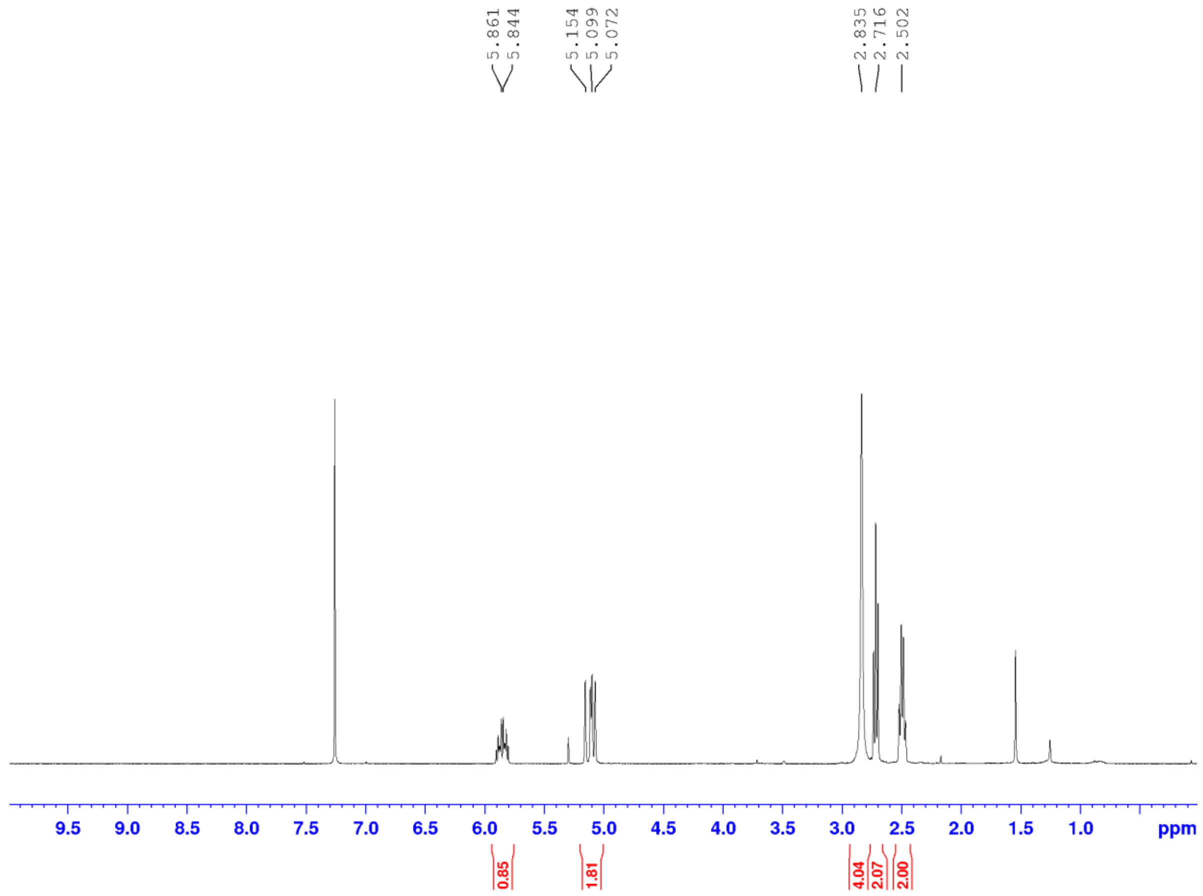

**S6** (NHS-alkane)

$^1H$ NMR $CDCl_3$

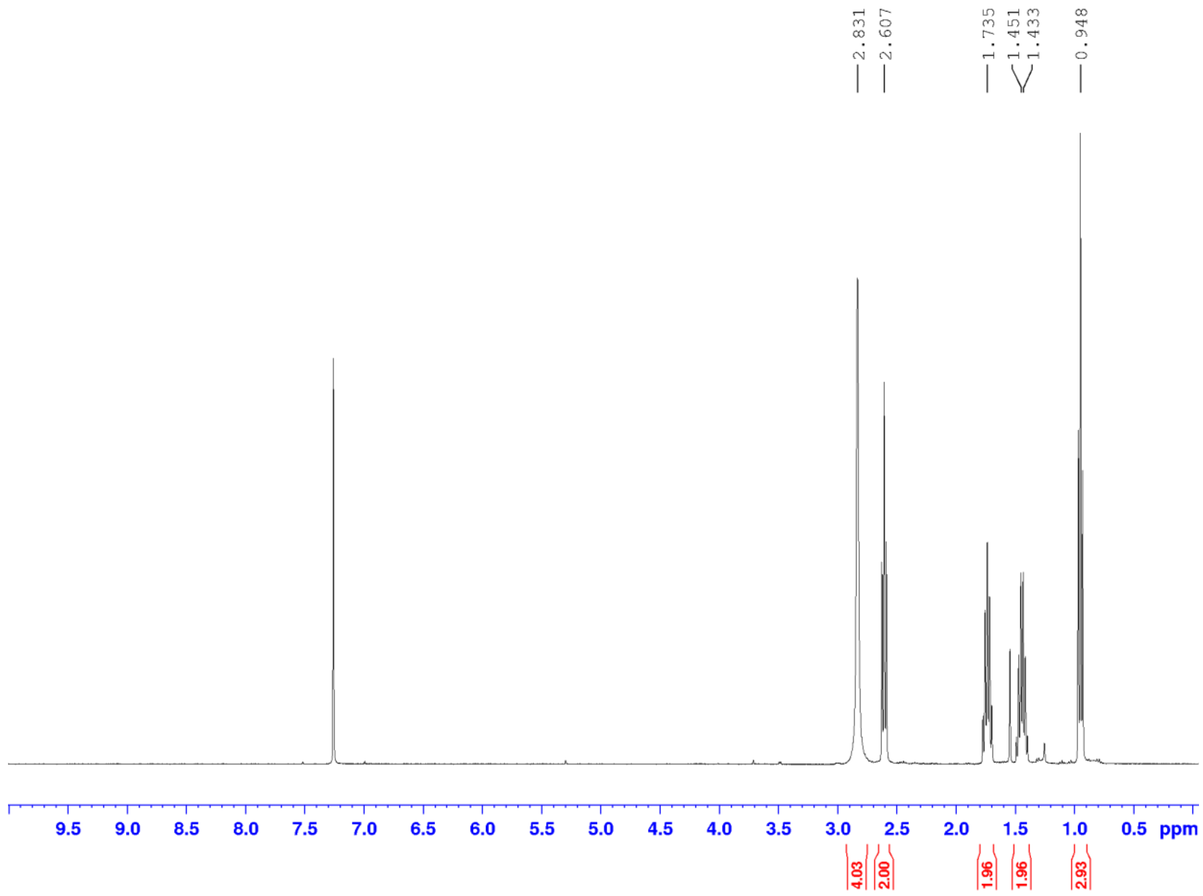

**4** (Sugar-Abz)

$^{1}H$ NMR $CDCl_3$

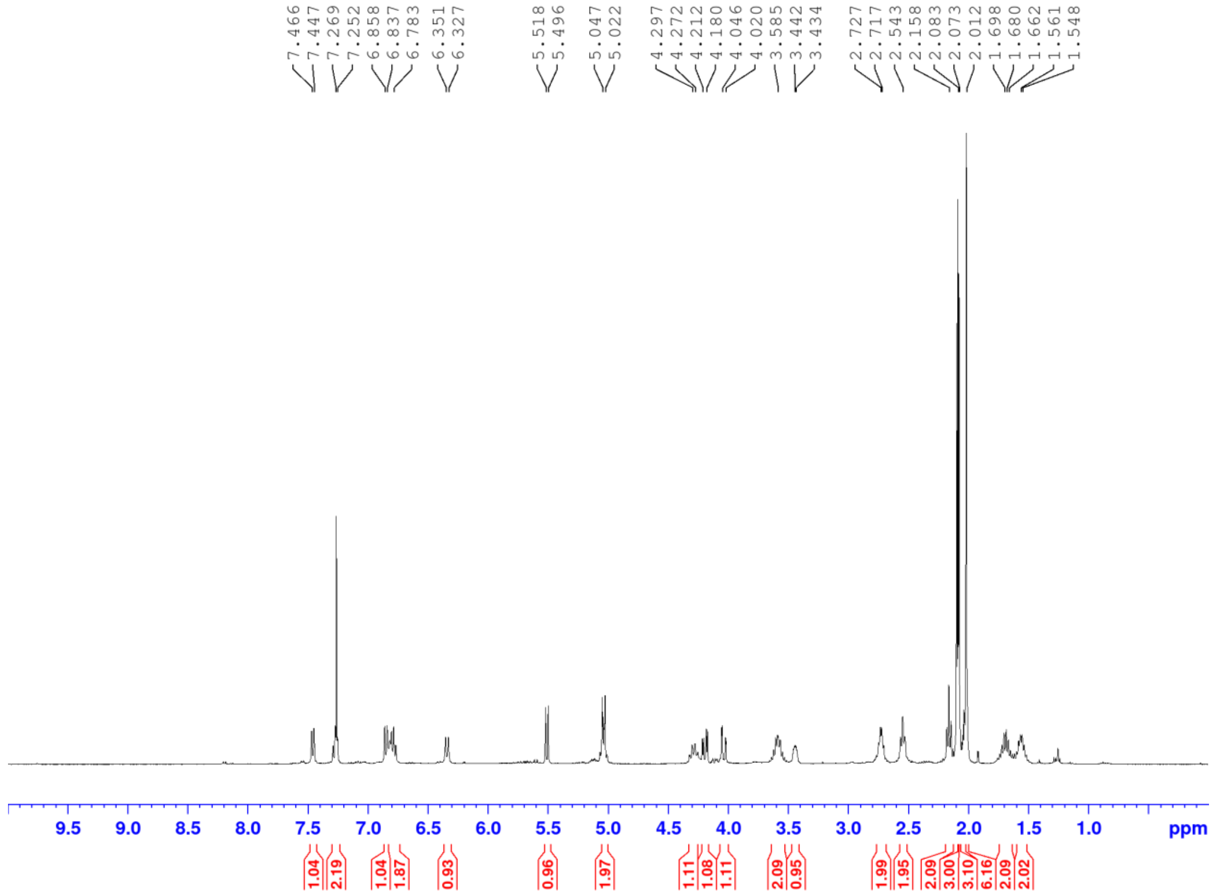

$^{13}C$ NMR $CDCl_3$

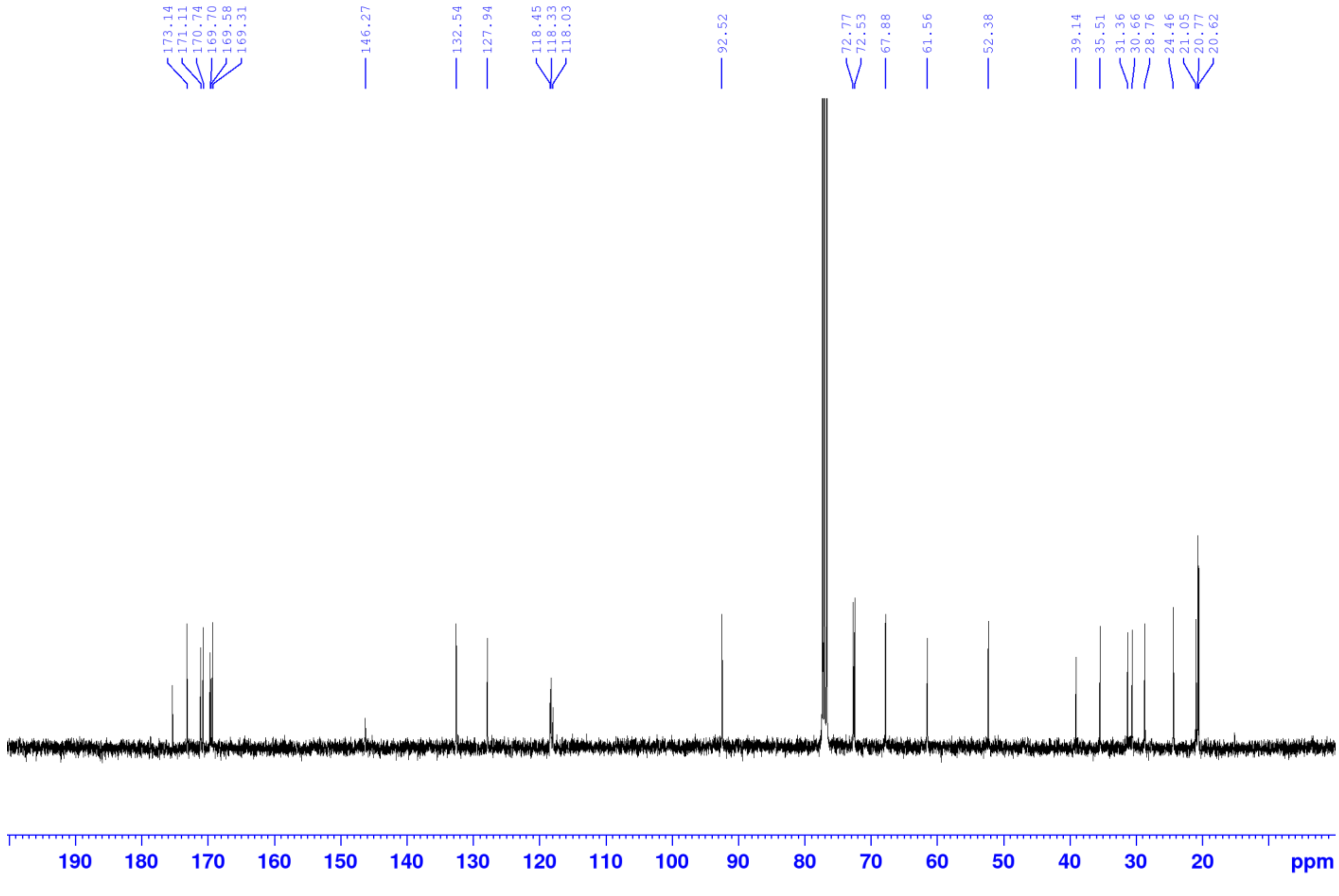

**5** (Sugar-Amine-free-Abz)

$^{1}H$ NMR $CDCl_3$

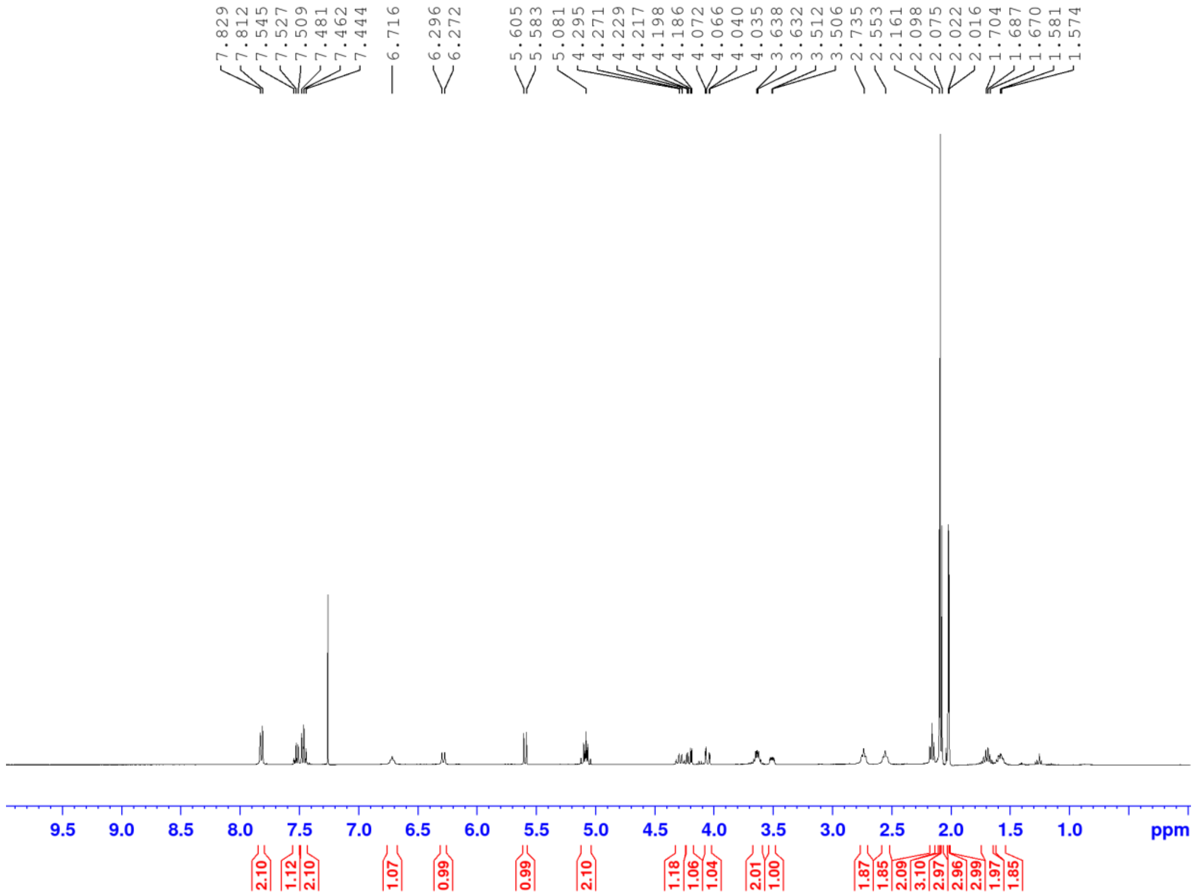

$^{13}C$ NMR $CDCl_3$

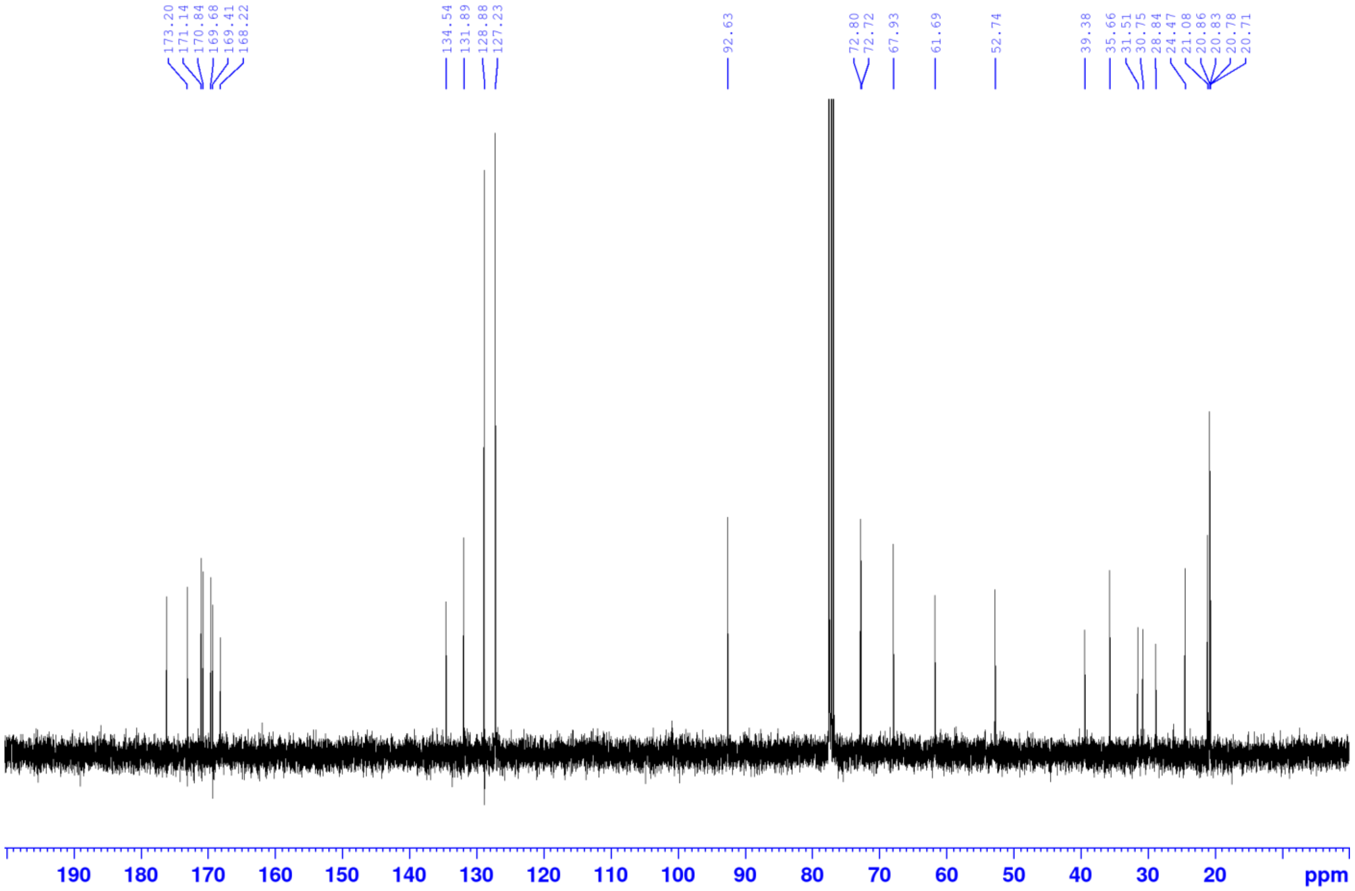

**S7** (Agl-Abz)

$^1$H NMR DMSO-d6

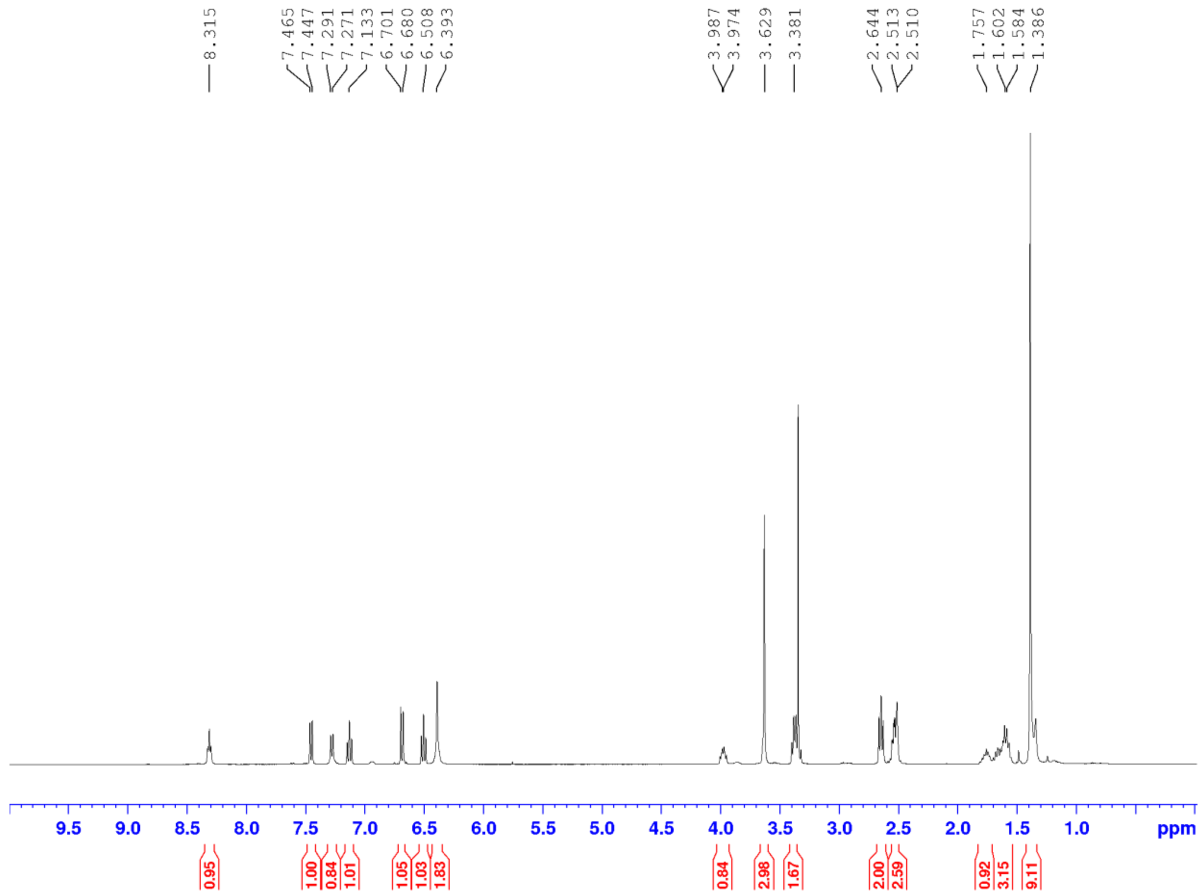

$^{13}$C NMR DMSO-d6

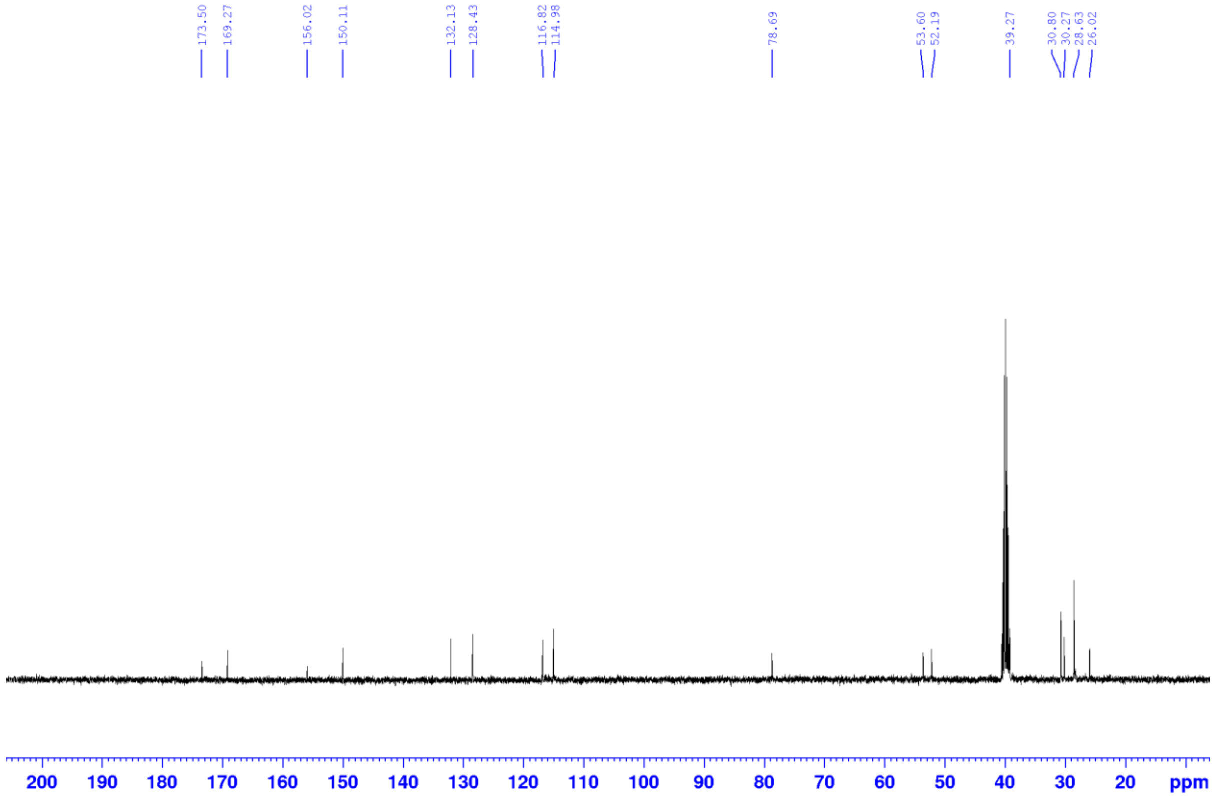

**S8** (Estrone-Abz)

$^{1}H$ NMR DMSO-d6

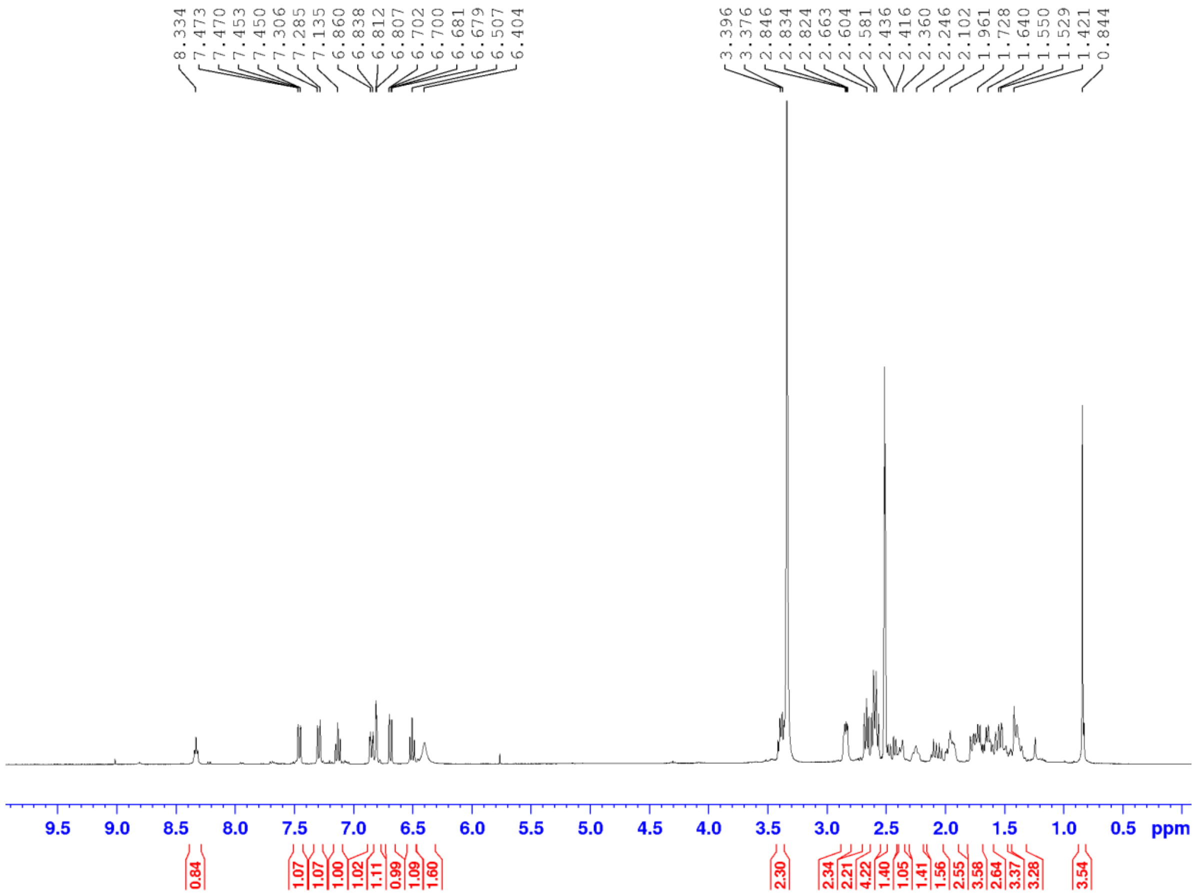

$^{13}C$ NMR DMSO-d6

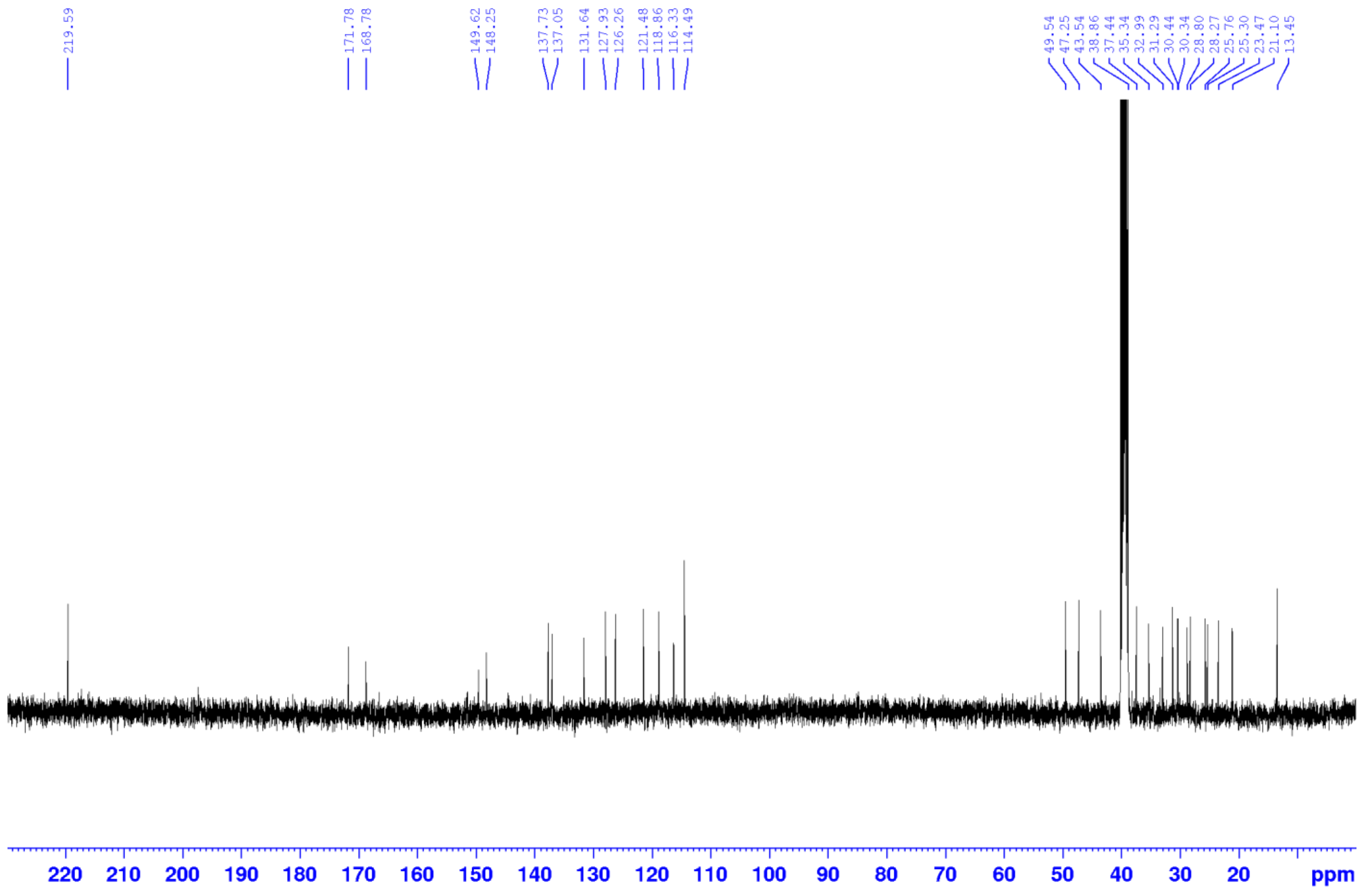

### 9. References

(1) Karmakar, B.; Banerji, J. An Expeditious, Efficient Green Methodology for the Boc Protection of Amines and Silyl Protection of Alcohols over Tungstophosphoric Acid-Doped Mesoporous Silica. *Tetrahedron Lett.* **2010**, *51* (29), 3855–3858. https://doi.org/10.1016/j.tetlet.2010.05.080

(2) Hurst, R. D.; Nieves, A.; Brichacek, M. Expanding Glycomic Investigations through Thiol-Derivatized Glycans. *Molecules* **2023**, *28* (4), 1956. https://doi.org/10.3390/MOLECULES28041956

(3) Baud, M. G. J.; Haus, P.; Leiser, T.; Meyer-Almes, F.-J.; Fuchter, M. J. Highly ligand efficient and selective N-2-(Thioethyl)picolinamide histone deacetylase inhibitors inspired by the natural product psammaplin A. *ChemMedChem* **2013**, *8* (1), 149–156. https://doi.org/10.1002/cmdc.201200450

(4) Li, J.; Liu, J.; Li, X.; Wang, T.; Zhang, Y.; Liang, J.; Zhang, X.; Li, C.; Sun, H.; Song, L.; Liang, D.; Xiong, Z. Design, Synthesis and Biological Evaluation of Cinnamoyl Tethered Betulinic Acid Derivatives as α-Glucosidase Inhibitors. *J. Mol. Struct.* **2025**, *1345*, 143024. https://doi.org/10.1016/j.molstruc.2025.143024

(5) Descotes, G.; Ramza, J.; Basset, J.-M.; Pagano, S.; Gentil, E.; Banoub, J. Synthesis and Structural Characterization of Macrocyclic Carbohydrate Derivatives Obtained from Catalytic Metathesis Reaction with Chloro-Aryloxide Complexes of Tungsten. *Tetrahedron* **1996**, *52* (33), 10903–10920. https://doi.org/10.1016/0040-4020(96)00612-6

(6) Kozłowska, A.; Dranka, M.; Zachara, J.; Pump, E.; Slugovc, C.; Skowerski, K.; Grela, K. Chelating Ruthenium Phenolate Complexes: Synthesis, General Catalytic Activity, and Applications in Olefin Metathesis Polymerization. *Chem. Eur. J.* **2014**, *20* (43), 14120–14125. https://doi.org/10.1002/chem.201403580

(7) Lee, Y. J.; Yadagiri, K.; Yang, Y.; Torres-Kolbus, J.; Deiters, A.; Liu, W. R. Genetically Encoded Unstrained Olefins for Live Cell Labeling with Tetrazine Dyes. *Chem. Commun.* **2014**, *50* (86), 13085–13088. https://doi.org/10.1039/C4CC06435F

(8) Oh, C.; Li, M.; Kim, E. H.; Park, J. S.; Lee, J. C.; Ham, S. W. Antioxidant and Radical Scavenging Activities of Ascorbic Acid Derivatives Conjugated with Organogermanium. *Bull. Korean Chem. Soc.* **2010**, *31* (12), 3513–3514. https://doi.org/10.5012/BKCS.2010.31.12.3513
